# Repeated transcranial ultrasound is associated with altered temporal organization of spontaneous wheel activity during sleep disruption in mice

**DOI:** 10.64898/2026.11.25.769357

**Authors:** Yutaro Hazama, Kodai Kawata, Kaede Takemura, Itsuki Sawada, Reon Konaka, Karin Shimizu, Riki Sato, Shuhei Iida, Maya Goto, Minto Arimura, Sakura Shinozawa, Ayano Matsushita, Soichiro Kimura, Hideo Ueda

## Abstract

**Background:** Motor-cortex-targeted transcranial ultrasound stimulation (tUS) can influence neural activity and motor responses, but its relationship to spontaneous behavior during sleep disruption (SD) remains unclear.

**Objective:** To examine how repeated tUS during sustained SD relates to the temporal organization of spontaneous running and inactivity bouts.

**Methods:** We recorded wheel activity in 36 mice for 17 days. On Days 13–17, mice in control (Ctrl) or wheel-confinement SD housing received bilateral motor-cortex-targeted tUS or sham. Another 32 mice followed the same treatment schedule and completed the retention phase of the novel object recognition test before open-field testing on Day 17. EEG/EMG recordings in 11 untreated mice characterized the SD sleep– wake phenotype. Hydrophone measurements in water and simulations using individual skull models characterized acoustic exposure.

**Results:** During the final three dark phases, SD/tUS mice had lower activity-resumption probability (0.132 vs 0.224) and longer median inactivity bouts (3.81 vs 2.48 min) than SD/Sham mice. Both housing-by-treatment interactions were significant. Within SD, neither total rotations nor running intensity differed significantly. The mean discrimination index was higher with tUS, although the Holm-adjusted comparison was nonsignificant. SD/tUS mice spent longer in the open-field center than SD/Sham mice. Across all 32 behavioral-test mice, lower resumption probability and longer pauses correlated with longer center time the next day. Untreated SD mice slept more and had more frequent sleep–wake transitions.

**Conclusions:** During ongoing SD, mice receiving repeated motor-cortex-targeted tUS were less likely than sham-treated mice to resume wheel running. Their longer pauses between runs may provide opportunities for sustained rest.

## 1. Introduction

Low-intensity transcranial ultrasound stimulation (tUS) is a noninvasive neuromodulation technique that uses acoustic energy transmitted through the skull to modulate neural activity [1,2]. Motor-cortex-targeted tUS can elicit motor responses in rodents [3,4] and alter cortical oscillations depending on stimulation parameters [5].

At a pulse repetition frequency of 40 Hz, tUS elicited depolarization in some cortical neurons and hyperpolarization in others in awake mice [6]. Patterned tUS of the mouse motor cortex can induce persistent changes in motor-evoked potentials and influence subsequent motor-skill acquisition when delivered before training [7]. Repeated motor-cortex-targeted tUS has also been reported to improve motor test performance in an MPTP mouse model of Parkinson’s disease [8]. These findings motivate examining repeated tUS beyond immediate motor responses and task-specific performance.

Spontaneous wheel running comprises bouts of activity separated by periods of inactivity. Motor-cortical circuits contribute to action initiation and execution [9], and neurons in the mouse secondary motor cortex exhibit distinct activity patterns around locomotor onset and cessation compared with sustained locomotion across contexts including wheel running [10]. Wheel-running microstructure can be characterized by within-bout running rate, bout duration, and interbout pauses [11]. In mice, spontaneous activity also exhibits greater day–night differences in rest-bout duration than in activity-bout duration [12]. Thus, similar rotation totals may arise from different patterns of activity resumption and inactivity persistence, even at comparable running intensities. These temporal features may therefore provide neurobiologically relevant information beyond aggregate activity measures and help characterize behavioral changes associated with neuromodulation.

Sleep physiology likewise depends on temporal organization as well as total sleep time. In mice, sleep fragmentation can increase sleep pressure despite relatively preserved sleep quantity [13], and disrupting sleep continuity can impair memory consolidation without reducing total sleep time [14]. In humans, sleep deprivation alters cortical excitability and plasticity-like responses to noninvasive brain stimulation [15,16], suggesting that neuromodulatory responses may depend on sleep-related physiological context. tUS targeting the motor cortex and other regions can alter measures of rapid eye movement (REM) and non-rapid eye movement (NREM) sleep in healthy and sleep-deprived mice [17]; repeated basal-forebrain-targeted tUS has also been investigated during repeated REM sleep deprivation [18]. However, mice alternate between running and non-running periods during sustained wakefulness [19], with differences in motor-cortical activity [20]. Wheel inactivity therefore cannot be equated with sleep. How repeated motor-cortex-targeted tUS relates to the temporal organization of spontaneous wheel activity outside stimulation sessions during ongoing sleep disruption remains unclear.

Wheel-based housing permits longitudinal monitoring of activity during sustained disruption. Wheel confinement to avoid water alters daily activity and sleep–wake patterns [21]; related housing paradigms have been used to induce chronic sleep fragmentation in a mouse model of Alzheimer’s disease [22]. Object recognition deficits and anxiety-like behavior have also been reported in a water-aversion-based model [23], suggesting functional changes beyond wheel activity. Building on these paradigms, we modeled sleep disruption (SD) by restricting cage-floor access without water and maintained the assigned housing conditions throughout the intervention. Concurrent EEG/EMG and wheel recordings in an independent untreated cohort characterized sleep–wake organization and its correspondence with wheel activity.

We examined whether repeated motor-cortex-targeted tUS was associated with altered temporal organization of spontaneous wheel activity during ongoing SD and whether these associations differed between control (Ctrl) and SD housing conditions. Using a Ctrl/SD × sham/tUS design, we assessed state transitions and persistence alongside total rotations and running intensity during dark phases following daytime treatment, without concurrent sonication. Water-field measurements and skull-based acoustic and thermal simulations characterized exposure and predicted heating. In a separate intervention cohort under the same housing and treatment conditions, the novel object recognition test (NORT) and open-field test (OFT) assessed object recognition and spatial exploration; exploratory analyses examined their associations with preceding wheel activity.

## 2. Materials and Methods

### 2.1. Animals and study design

Seventy-nine male C3H/HeNSlc mice (Japan SLC; approximately 8 weeks old at arrival) were housed individually under a 12-h light/dark cycle (lights on at 07:00, ZT0; off at 19:00, ZT12), at 22 ± 2°C and 55 ± 10% relative humidity, with food and water ad libitum. Procedures followed Josai University’s animal-experiment regulations and were approved under protocols JU 22097, JU 23020, JU 24020, JU 25018, and JU 26018.

The three independent cohorts comprised untreated EEG/EMG mice (Ctrl, n = 4; SD, n = 7), wheel-activity mice (n = 36; 9 per Ctrl/Sham, Ctrl/tUS, SD/Sham, and SD/tUS group), and NORT/OFT mice (n = 32; 7 per Ctrl group and 9 per SD group). The experimenter assigned mice with the aim of balancing group sizes, without deliberate selection by mouse characteristics. No a priori sample-size calculation was performed.

### 2.2. SD conditions and experimental timeline

All mice initially had floor and wheel access. At ZT8 on Day 10, SD mice were confined to the wheel, while Ctrl mice retained access to both. Intervention cohorts remained in SW-15S (Ctrl) or SW-15-SD (SD) cages (Melquest) through Day 17, with daily tUS or sham on Days 13–17 at ZT2–4. Wheel recording spanned Days 1–17 in the wheel-intervention and EEG/EMG cohorts (Figure 1, Supplementary Methods S1, S5 and S9– S10).

**Figure 1.**
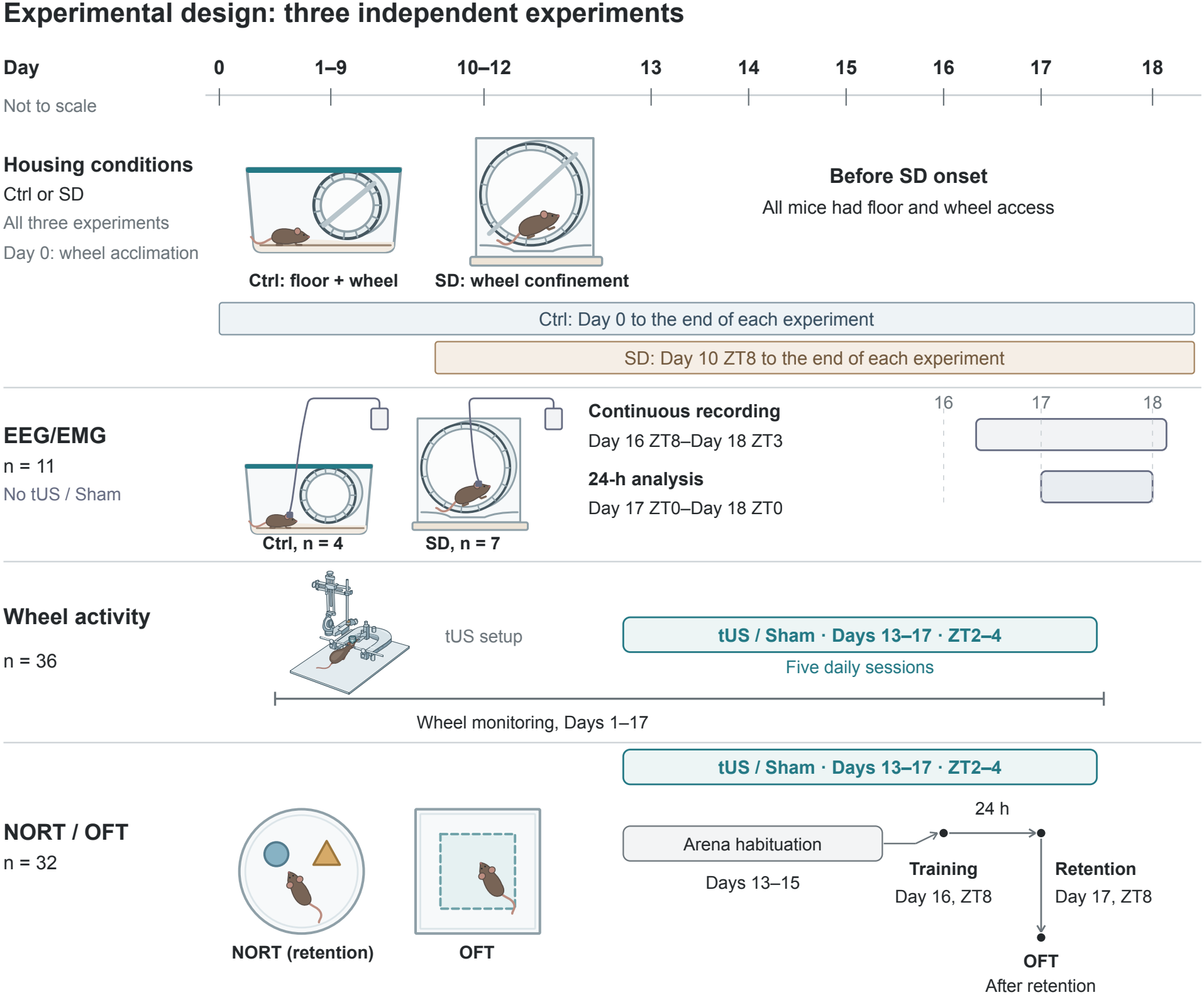
Experimental design and timelines for three independent cohorts. The timeline is not drawn to scale. Wheel acclimation began on Day 0. Ctrl mice had access to both the cage floor and the wheel from Day 0 through the end of each experiment. SD mice had the same access until Day 10 at ZT8, when they were confined to the wheel for the remainder of each experiment. ZT0 denotes lights on, and ZT12 denotes lights off. The independent EEG/EMG cohort (n = 11; Ctrl, n = 4; SD, n = 7) received neither tUS nor sham. Wired EEG/EMG was recorded in the assigned Ctrl and SD conditions. Continuous EEG/EMG recording extended from Day 16 at ZT8 to Day 18 at ZT3. The 24-h analysis interval extended from Day 17 at ZT0 to Day 18 at ZT0. The wheel-activity and NORT/OFT intervention cohorts comprised Ctrl/Sham, Ctrl/tUS, SD/Sham, and SD/tUS groups. The wheel-activity cohort included nine mice per group (n = 36). The NORT/OFT cohort (n = 32) included seven mice per Ctrl group and nine mice per SD group. Bilateral motor-cortex-targeted transcranial ultrasound stimulation (tUS) or sham procedures were administered daily on Days 13–17 between ZT2 and ZT4. Wheel activity was monitored on Days 1–17 in the wheel-activity cohort. In the separate behavioral-testing cohort, mice were habituated to the empty novel-object-recognition test (NORT) arena on Days 13–15, followed by training on Day 16 at ZT8 and retention testing on Day 17. On Day 17, NORT retention testing began at ZT8 after the daily tUS/sham procedure and was followed by the open-field test (OFT).

### 2.3. Ultrasound stimulation and sham procedures

tUS used a 20-mm planar piezoelectric transducer (Honda Electronics) with a resin collimator (2-mm outlet). After scalp hair removal on Day 10 in both groups, tUS mice were anesthetized with isoflurane (5% induction, 1.5–2.5% maintenance), secured stereotaxically, and targeted bilaterally at motor cortex (AP +1.5 mm, ML ±1.5 mm relative to bregma). Gel provided scalp–collimator coupling.

Ultrasound was delivered at 1.078 MHz, with 40-Hz pulse repetition and a 50% duty cycle (12.5 ms ON/OFF), in six alternating 5-min left/right blocks. Excluding repositioning, the blocks totaled 30 min (15 min per hemisphere), including 15 min of ultrasound emission. Sham mice received matched isoflurane conditions and anesthesia duration, but were placed in a separate box without head fixation, transducer contact, gel, or ultrasound. Both groups were warmed during anesthesia. Apparatus and exposure details are in Supplementary Methods S2 and Table S1 [24].

### 2.4. Acoustic measurements and modeling

Free-field pressure was measured in three orthogonal water planes using a calibrated needle hydrophone (HNR-0500; Onda Corporation). Temporal-average intensity included pulse-OFF periods; the sampled XY maximum served as the intensity-ratio denominator. Source amplitude was calibrated against hydrophone waveforms along a water depth profile.

Ten independent C3H/HeJ CT skull models from MusMorph were combined with a common MRI template and approximate brain-atlas registration [25–27]. The source was reconstructed from the measured water-plane intensity and carrier phase. Left and right acoustic fields were simulated separately using a k-space model (120-µm grid) [28,29]. In modeled brain and effective cerebrospinal fluid (CSF), the peak-connected region at ≥−6 dB relative to the pressure-amplitude peak was extracted for each side. For each side, motor-area composition was the percentage of atlas-labeled voxels in the predicted exposure region assigned to the primary motor area (MOp) or secondary motor area (MOs). Composition and peak intensity were each averaged across sides within skulls. Acoustic heat sources entered a Pennes bioheat model (80-µm grid), with temperature rise estimated against matched no-sonication simulations [30] (Supplementary Methods S3–S4 and Table S1).

### 2.5. EEG/EMG recording and wheel–vigilance correspondence

After electrode implantation under isoflurane, EEG/EMG mice recovered for one week and underwent tether habituation. We recorded at 400 Hz in cable-compatible wheel cages (RWC-15/RWC-15-SD, Melquest) from Day 16 ZT8 to Day 18 ZT3 and analyzed the 24 h beginning at Day 17 ZT0.

FASTER2 (v0.5.5) scored 8-s epochs as Wake, NREM or REM without group labels or manual correction. Sleep comprised NREM and REM. Occupancy was the fraction of valid time in each state. We counted Sleep↔Wake transitions per valid hour. We calculated conditional probabilities per mouse from valid seconds of overlap with 1-min wheel bins (Section 2.6), weighting mice equally. Permutation tests compared mouse-level absolute EEG band power from 4-s wake windows between housing groups, with Holm adjustment across four bands (Supplementary Methods S1.2 and S9–S10).

### 2.6. Wheel recording and behavioral-state analysis

We recorded 1-min wheel counts with CIF-4A (intervention cohorts) or TUSB-S03CN3 (independent EEG/EMG cohort). Counts indicated running (R, ≥1) or inactivity (I, zero). Missing bins were excluded. For valid adjacent bins, the two-state transition probabilities [12] were P(I→R) = N(IR)/[N(II) + N(IR)] and P(R→I) = N(RI)/[N(RR) + N(RI)]. No pairs crossed missing data or analysis boundaries.

Dark-phase (ZT12–24) outcomes used Days 15–17, selected after inspection of the daily group trajectories (Figure 4A). Within each mouse, daily dark-phase rotation totals, transition probabilities, running intensity (rotations per active minute), and median durations of contiguous inactive bins were averaged equally across those three days. Median durations retained window-edge bouts at their observed lengths.

### 2.7. Behavioral tests

NORT followed the habituation, training, and retention sequence described by Lueptow [31]. Mice explored two identical objects on Day 16 and underwent retention testing 24 h later after tUS or sham. Object positions were fixed (familiar on the left, novel on the right). DeepLabCut scoring used the first 300 s of cumulative observation excluding time on objects during the 600-s test. All 32 mice completed this window with ≥20 s of exploration. The discrimination index was (T_novel_ − T_familiar_)/(T_novel_ + T_familiar_).

Approximately 2 h later, the same mice underwent a 20-min OFT in a 50 × 50 cm arena. Outcomes were total distance and time in the central 30 × 30 cm. Both tests were scored blind to group (Supplementary Methods S6–S7).

### 2.8. Statistical analysis

Each mouse was the experimental unit. Tests were two-sided (α = 0.05). Reported 95% confidence intervals were pointwise. In the EEG/EMG cohort, unpaired equal-variance Student’s t tests compared 24-h Sleep occupancy and Sleep↔Wake transition frequency between groups. Welch’s t tests compared dark-minus-light occupancy differences, with Holm adjustment across Wake, NREM and REM.

For the intervention cohorts, two-factor linear models included housing, treatment and their interaction, with HC3 standard errors (transformations in Supplementary Methods S8.1). Holm adjustment covered three wheel or four NORT/OFT group contrasts per outcome. Interactions were tested separately without cross-outcome adjustment. In the behavioral-test cohort, we pooled all 32 mice for Spearman correlations between two-night wheel averages (Day 15–16 dark phases) and Day 17 NORT/OFT outcomes (Supplementary Methods S8.3).

## 3. Results

### 3.1. Acoustic exposure and predicted heating

Within the measured water XY plane, the maximum peak negative pressure was 0.300 MPa, and the maximum temporal-average intensity was 1.194 W/cm^2^ (Figure 2A). The transverse −6-dB intensity widths were 1.59 mm along X and 1.57 mm along Y, whereas the axial intensity profiles remained above threshold at the scan boundary (Supplementary Figure S1).

**Figure 2.**
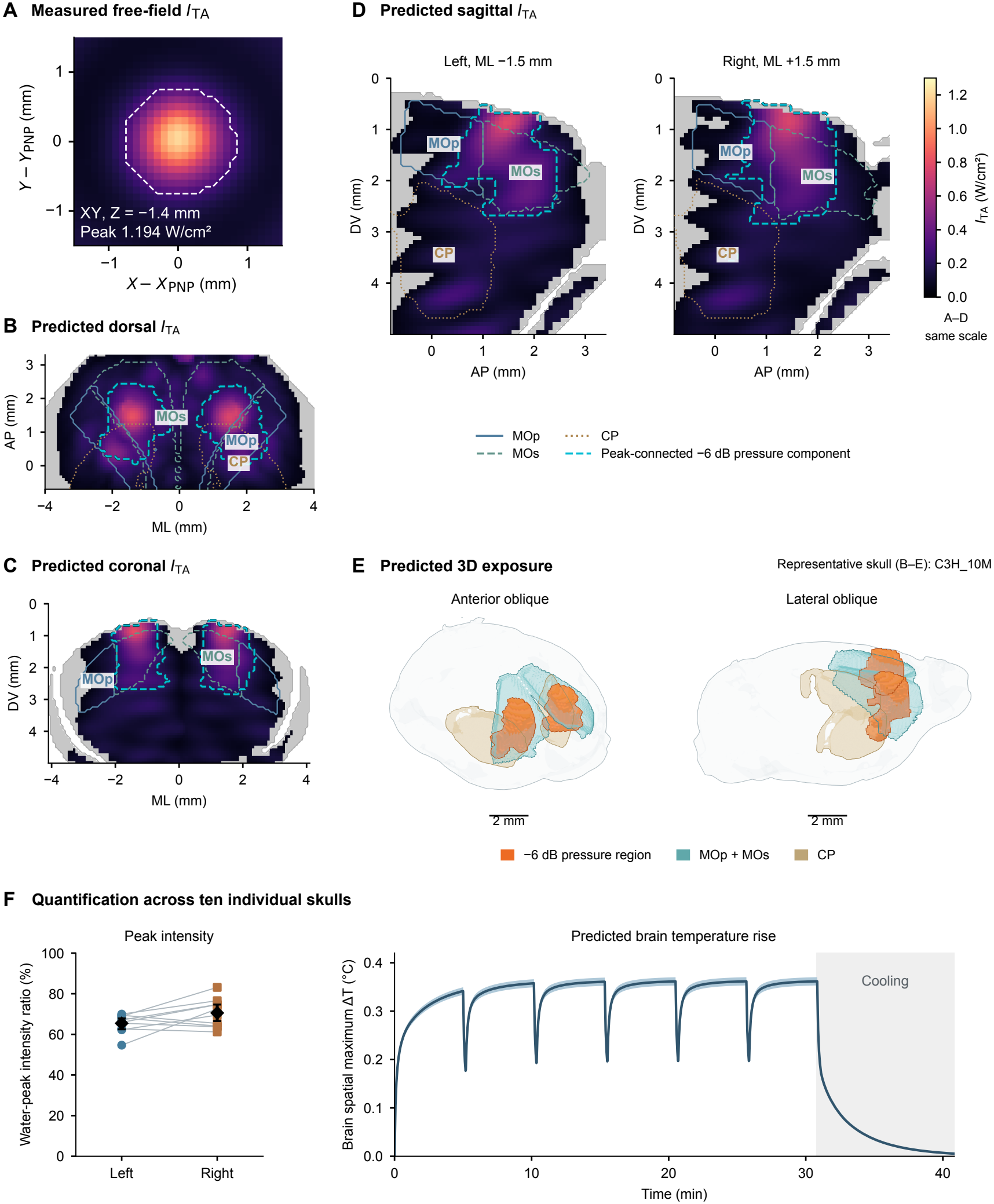
Measured water intensity, predicted transcranial exposure, and brain heating. (A) Measured water temporal-average intensity (I_TA) in the XY plane at Z = −1.4 mm (maximum, 1.193585 W/cm^2^). The transverse axes are centered on the measured peak negative pressure maximum. The white dashed contour encloses the four-neighbor peak-connected −6-dB intensity region (≥25.12% of the measured XY maximum). A–D share an absolute linear 0–1.3 W/cm^2^ color scale, without panelwise normalization. (B–E) Predicted fields for representative skull C3H_10M (selection rule in Supplementary Methods S3.5). For each independently simulated stimulation side, the peak-connected −6-dB pressure-amplitude component comprises voxels within brain plus effective CSF with |p| ≥50.12% of that side’s pressure-amplitude peak, connected by six-neighbor adjacency. Bilateral displays do not represent simultaneous sonication. (B) Dorsal maximum-intensity projection. (C) Coronal section at AP +1.5 mm. B and C show voxelwise maxima of separately modeled left/right intensities and spatial unions of their pressure components (projected across depth in B). (D) Left/right sagittal sections at ML −1.5/+1.5 mm show each corresponding side separately. In B–D, cyan dashed contours indicate pressure-defined components, not thresholds of the displayed intensity maps. Anatomical outlines use approximate atlas registration, and intensities below 0.0455 W/cm^2^ are hidden only for display. (E) Anterior and lateral oblique views (2-mm scale bars). Orange shows the pressure components, teal the primary and secondary motor areas (MOp + MOs), tan the caudoputamen (CP), and the pale shell the brain plus effective CSF. Surface colors do not encode intensity. (F) Ten-skull summaries. Left: the peak-intensity ratio is 100 × each side’s maximum intracranial I_TA (brain plus effective CSF) divided by the measured water-XY maximum (1.193585 W/cm^2^), not total acoustic-power transmission. Connected circles (left) and squares (right) show paired skull values; black diamonds and error bars indicate side-specific means and skull-bootstrap 95% confidence intervals (n = 10). The bilateral mean was 68.00 ± 4.48% (mean ± between-skull SD), after averaging sides within skulls. Right: predicted brain temperature rise during six alternating 5-min blocks and cooling (gray). At each time point, the spatial brain maximum was averaged over both starting orders within each skull, then across skulls (line, mean; band, between-skull SD). The mean curve peaked at 0.362°C; Supplementary Figure S3 instead summarizes maxima taken over space and time for each skull and starting order before averaging.

Across ten skulls, the predicted spatial peak of the intracranial pressure amplitude, averaged across sides within each skull, was 0.2261 ± 0.0064 MPa. The predicted peak temporal-average intensity was 0.8116 ± 0.0535 W/cm^2^, or 68.00 ± 4.48% of the measured water-XY maximum (mean ± SD; Figure 2F, left). The primary motor area (MOp) and secondary motor area (MOs) together accounted for 73.37 ± 6.12% of the atlas-labeled predicted exposure volume (mean ± SD; Supplementary Figure S2). Predicted exposure overlapped the caudoputamen in all ten models (e.g., Figure 2E). The mean brain temperature-rise curve peaked at 0.362°C (Figure 2F, right). Peak rises calculated separately for each skull and stimulation order averaged 0.3759 ± 0.0105°C (Supplementary Figure S3).

### 3.2. Sleep–wake characteristics of the SD model

In the independent EEG/EMG experiment without tUS (Ctrl, n = 4; SD, n = 7), mean 24-h Sleep occupancy was 61.23% in SD mice and 49.47% in Ctrl mice (between-group difference [SD − Ctrl], 11.76 percentage points; 95% confidence interval [CI], 8.83 to 14.69; P = 7.94 × 10^−6^; Figure 3A). The dark-minus-light difference in Wake occupancy was smaller in SD mice, at 15.50 percentage points versus 24.53 in Ctrl mice (between-group difference, −9.03 percentage points; 95% CI, −13.84 to −4.22; Holm-adjusted P = 0.00693; Supplementary Table S2).

**Figure 3.**
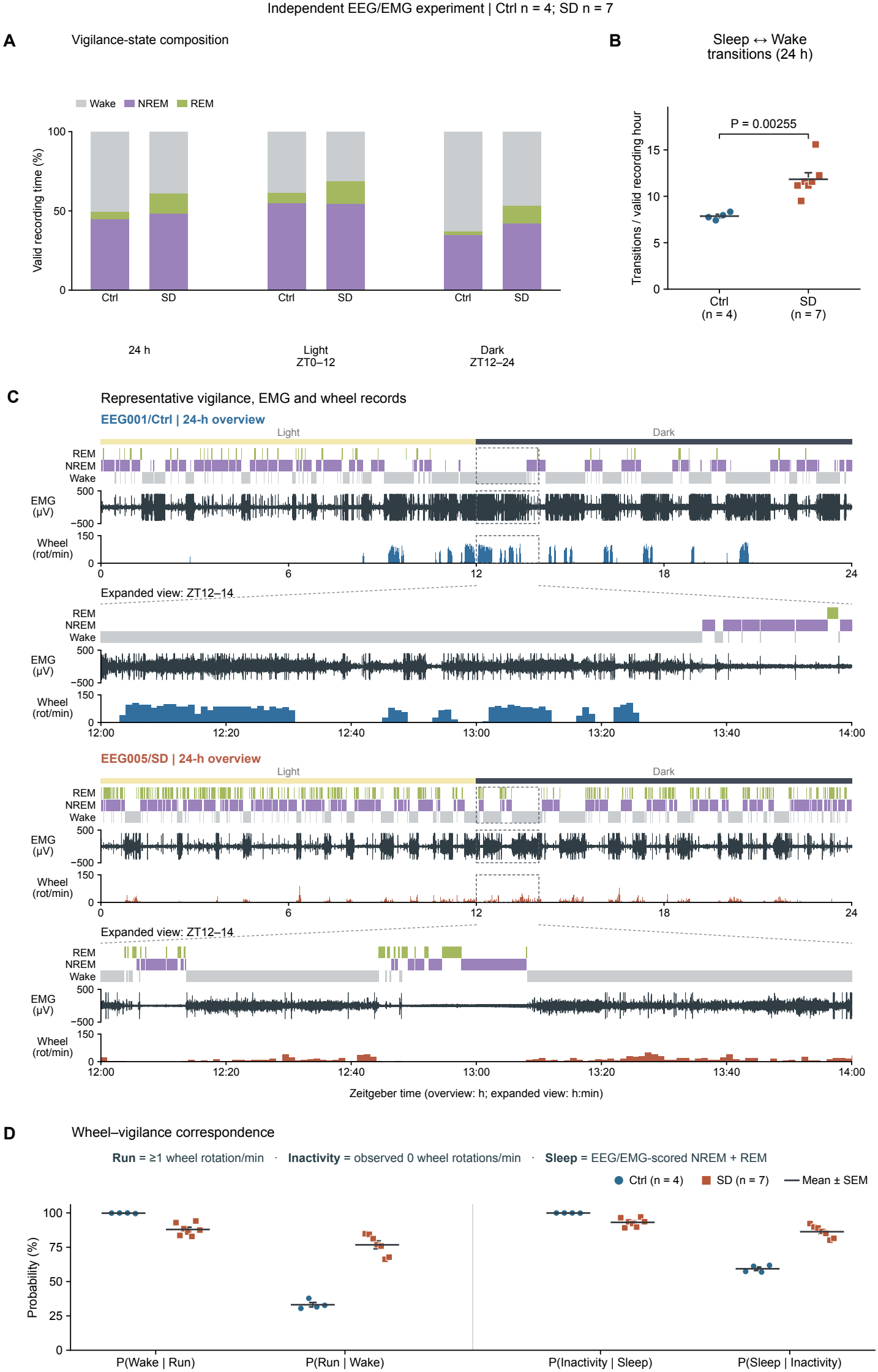
Sleep–wake organization and wheel–vigilance correspondence in the SD model. (A) Group-mean Wake, NREM and REM composition across 24 h and the light (ZT0–12) and dark (ZT12–24) phases. (B) Bidirectional Sleep↔Wake transitions per valid recording hour over 24 h; Sleep combines NREM and REM. Points show individual mice, short horizontal lines show group arithmetic means, and error bars show SEM at the group center. The bracket shows the P value from a two-sided unpaired equal-variance Student t test comparing Ctrl and SD.(C) Representative Ctrl (EEG001) and SD (EEG005) records, each with a 24-h overview and expanded ZT12–14 view. Vigilance used 8-s epochs, raw EMG was acquired at 400 Hz, and wheel counts used 1-min bins. (D) Conditional percentages relating Wake or Sleep to wheel activity, calculated from valid overlapping time. Run denotes at least one wheel rotation per minute, and Inactivity denotes an observed zero. Within each pair, the two probabilities share the same overlap duration in the numerator (Wake with Run, or Sleep with Inactivity) and have different denominators defined by the conditioning state. Points show individual mice, horizontal lines show group means, and error bars show SEM. Ctrl n = 4; SD n = 7. These mice received no tUS. The 24-h interval was Day 17 ZT0 to Day 18 ZT0.

Over 24 h, the bidirectional Sleep↔Wake transition frequency was higher in SD mice, at 11.83 transitions per valid recording hour versus 7.86 in Ctrl mice (between-group difference, 3.97 transitions/h; 95% CI, 1.80 to 6.14; P = 0.00255; Figure 3B). Sleep bouts were shorter in SD than Ctrl mice in the light phase (mean of individual medians, 303.4 vs 403.0 s, P = 0.0306). The dark-phase difference was not significant (280.0 vs 310.0 s, P = 0.602). In both phases, SD mice had shorter Wake bouts and more frequent Sleep↔Wake transitions (Supplementary Figure S5 and Table S2). Waking delta power was 2.25 times as high in SD as in Ctrl mice (geometric means, 1,216 vs 540 µV^2^; Holm-adjusted P = 0.0242; Supplementary Figure S8A).

Representative 24-h records and expanded views of vigilance states, raw EMG, and wheel activity are shown in Figure 3C. Group means of mouse-level percentages showed that Wake occupied 87.97% of the time within 1-min bins classified as wheel-active in SD mice and 99.85% in Ctrl mice. Conversely, wheel-active bins accounted for 76.79% and 33.13% of Wake time, respectively. Sleep occupied 86.34% of wheel-inactive time in SD mice and 59.34% in Ctrl mice; wheel-inactive bins accounted for 93.15% and 99.95% of Sleep time, respectively (Figure 3D). Wheel-inactive bins contained both Sleep and Wake, and Wake was not restricted to wheel-active bins. Individual records and phase-specific correspondence appear in Supplementary Figures S4–S7; wheel and rhythm profiles, in Supplementary Figures S9–S12.

### 3.3. Longitudinal wheel activity and analysis window

In the wheel intervention cohort, daily rotations increased before SD onset and remained lower in SD than in Ctrl mice thereafter (Figure 4A). The 24-h totals exclude recording gaps during procedures and transport. On Days 15–17, mean totals were lower in SD/tUS than in SD/Sham mice. We selected these days after inspecting the group trajectories. Hourly profiles showed lower dark-phase activity in SD than in Ctrl mice, with generally lower values in SD/tUS than in SD/Sham mice (Figure 4B). We therefore examined running intensity and the temporal organization of running and inactivity bouts during ZT12–24.

**Figure 4.**
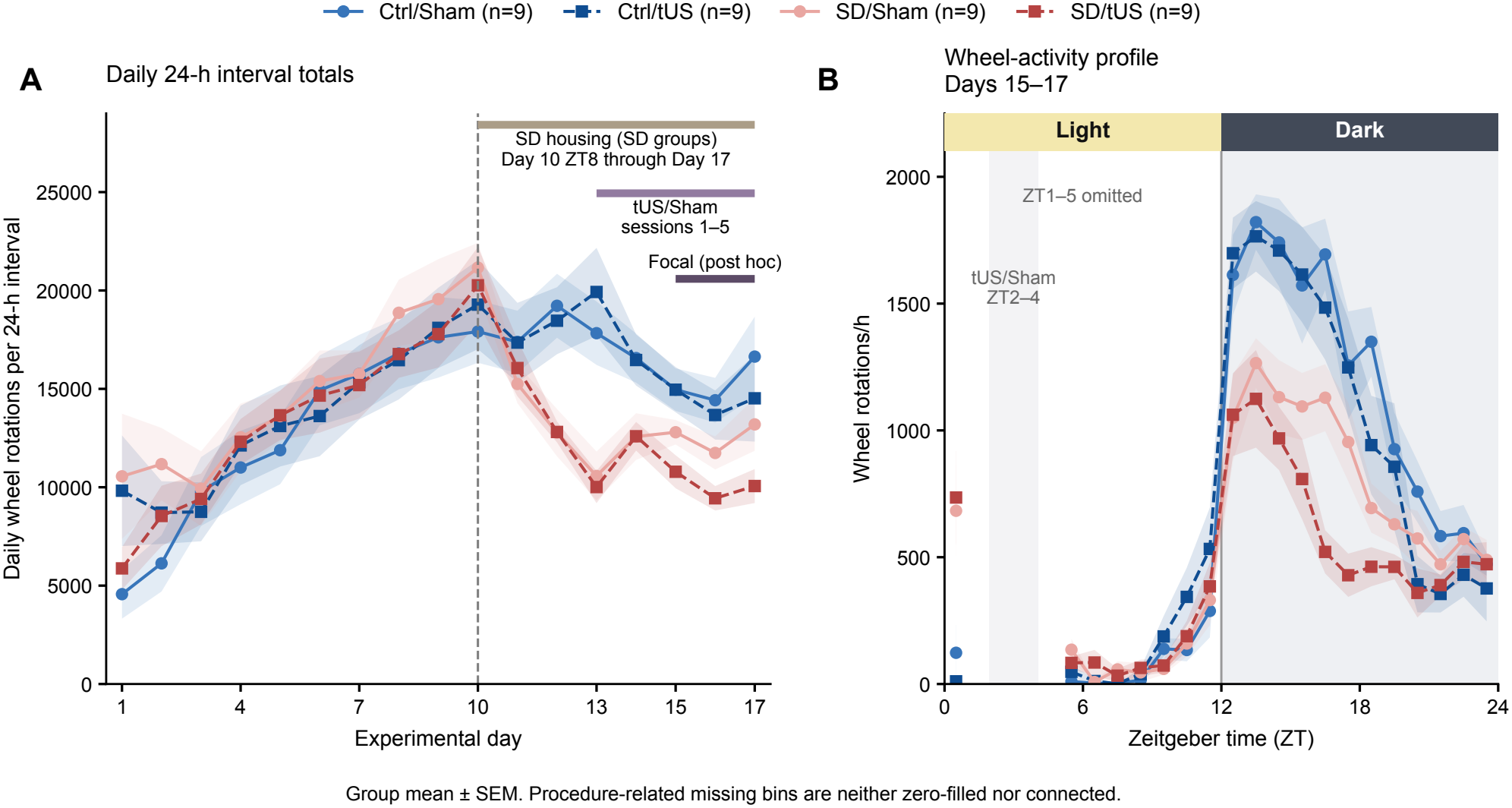
Longitudinal and diurnal wheel-activity profiles. (A) Calendar 24-h interval rotation totals (ZT0–24; 07:00 to 07:00), Days 1–17, mean ± SEM (n = 9/group). These intervals include recording gaps and are not 24 h of uninterrupted observation. The SD condition began on Day 10 at ZT8 and continued through Day 17; tUS/sham sessions occurred on Days 13–17 at ZT2–4. (B) Days 15–17 hourly profiles aligned to the 07:00 day boundary, using complete observed hours. Values were averaged across three days within each mouse, then across mice as mean ± SEM. Gray marks scheduled ZT2–4 procedures. Hourly bins spanning ZT1–5 were omitted because of recording unavailability associated with procedures and transport. No values, connecting curves, or SEM bands were added across this interval. Neither panel includes time-specific tests. The Days 15–17 window was selected after inspection of the daily wheel-activity trajectories in A and was used for the dark-phase summaries in Figure 5B–G.

Mean dark-phase rotation totals were 10,054 in SD/Sham mice and 7,541 in SD/tUS mice, but the between-group difference was nonsignificant after Holm adjustment (P = 0.0504). The housing condition × treatment condition interaction was also nonsignificant (P = 0.3834; Figure 5B). Dark-phase rotation totals were lower in SD/Sham mice than in Ctrl/Sham mice (Holm-adjusted P = 0.0324). Running intensity was 23.04 rotations per active minute in SD/Sham mice and 22.56 in SD/tUS mice, with no significant between-group difference (difference [SD/tUS − SD/Sham], −0.48; 95% CI, −5.74 to 4.77 rotations per active minute; Holm-adjusted P = 0.887; Figure 5C). All comparisons are reported in Supplementary Table S3.

**Figure 5.**
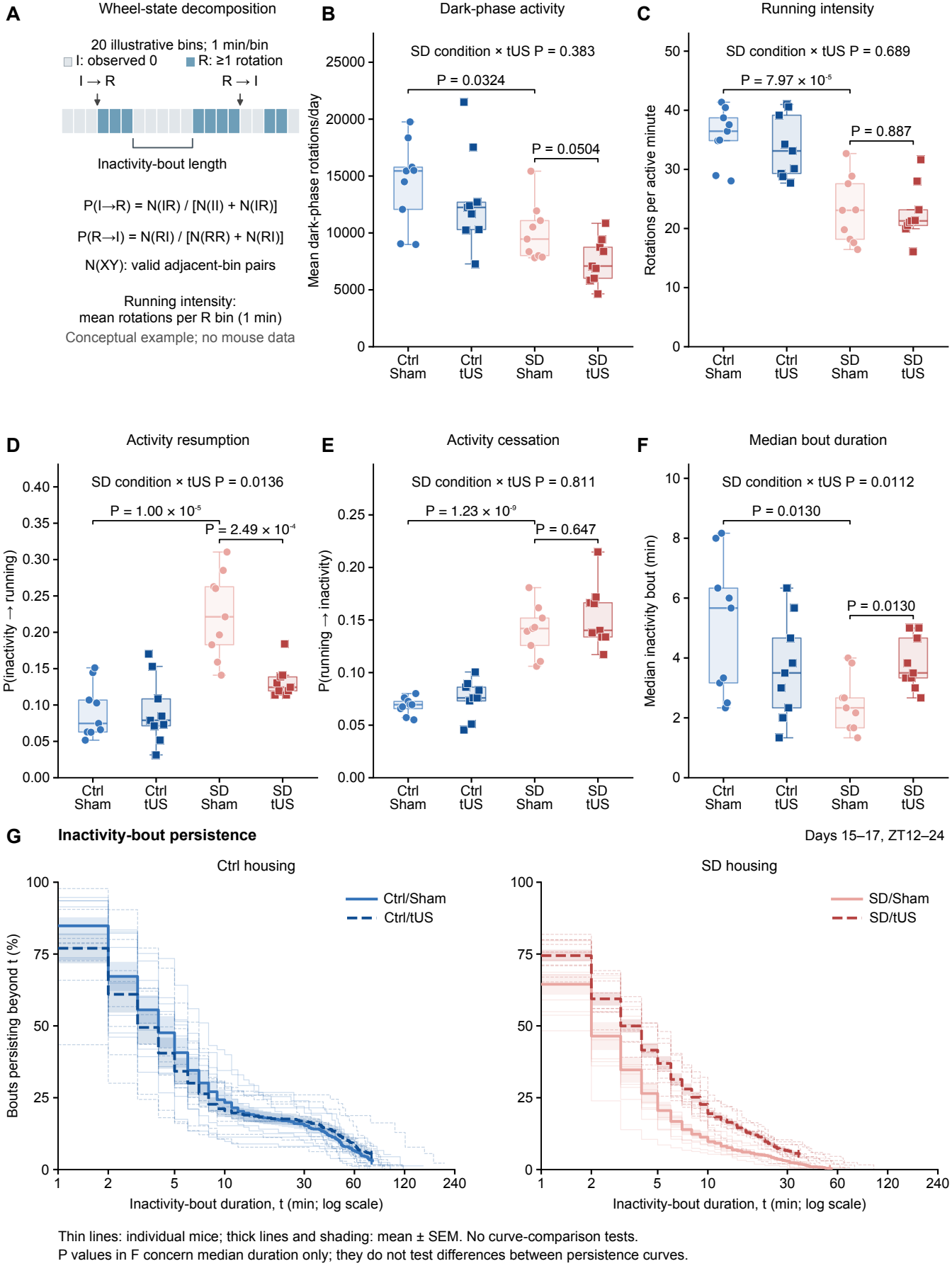
Conceptual decomposition, dark-phase wheel activity, and inactivity-bout persistence. (A) Conceptual illustration of twenty 1-min wheel-state bins, not derived from mouse data. Running (R) denotes at least one rotation and inactivity (I) denotes an observed zero. N(XY) denotes the number of valid adjacent-bin pairs with states X and Y. P(I→R) = N(IR)/[N(II)+N(IR)], and P(R→I) = N(RI)/[N(RR)+N(RI)]; staying in the starting state is therefore included in each denominator. Missing bins are not treated as inactivity, and adjacent pairs do not cross missing bins or analysis-window boundaries. An inactivity bout comprises consecutive inactive bins; running intensity is the mean number of rotations per active 1-min bin. (B) Total dark-phase rotations. (C) Running intensity. (D) P(I→R). (E) P(R→I). (F) Median inactivity-bout duration. Panels B–F summarize Days 15–17, ZT12–24. Each point represents one mouse (n = 9 per group). Boxes span the 25th–75th percentiles with median lines, whiskers extend to the most extreme observations within 1.5 times the interquartile range. All observations are shown, including those beyond the whiskers. For F, each mouse’s plotted value is the average of its three daily medians of window-bounded inactivity runs. Statistical results were obtained with HC3 linear models on the specified analysis scales. Brackets show P values for the displayed group contrasts, with Holm adjustment across three contrasts per wheel outcome. Labels above panels B–F give the housing × tUS interaction P values. Ctrl/tUS versus Ctrl/Sham comparisons are reported in Supplementary Table S3. (G) Inactivity-bout persistence during the same three dark phases, shown as the estimated percentage of bouts lasting longer than duration t. Kaplan–Meier-type curves were calculated separately for each mouse and day. Three daily curves were averaged equally within each mouse, and mice were weighted equally rather than pooling their bouts. Thin lines show individual-mouse curves; thick lines and shading show the mean ± SEM across nine mice. Sham and tUS are shown by solid and dashed lines, respectively. The duration axis is logarithmic. Each mouse curve is shown only where all three daily curves are estimable; the group mean is shown only where all nine mouse curves are estimable. End marks on the thick curves identify the end of this group support, not the maximum bout length. Group-comparison P values apply only to the median endpoint in F; the persistence curves in G were not compared inferentially. F retains window-edge bouts, whereas G excludes unknown-onset bouts and right-censors known-start bouts at dark-phase end (Supplementary Methods S5.3).

### 3.4. Temporal organization of activity and inactivity

Activity-resumption probability P(I→R) was higher in SD/Sham mice than in Ctrl/Sham mice (0.224 versus 0.091; Holm-adjusted P = 1.00 × 10^−5^). Within SD, activity-resumption probability was lower in SD/tUS mice than in SD/Sham mice (0.132 versus 0.224; Holm-adjusted P = 0.000249), with a significant housing condition × treatment condition interaction (P = 0.0136; Figure 5D). Ctrl/Sham and Ctrl/tUS group means were nearly identical. Activity-cessation probability P(R→I) was higher in SD/Sham mice than in Ctrl/Sham mice, but no significant difference was detected between tUS and Sham within SD (Holm-adjusted P = 0.647; Figure 5E).

Group means for the mouse-level median inactivity-bout duration endpoint were 2.48 min in SD/Sham mice and 3.81 min in SD/tUS mice, with longer durations in the SD/tUS group (Holm-adjusted P = 0.0130). The housing condition × treatment condition interaction was also significant (P = 0.0112; Figure 5F). The corresponding value in SD/Sham mice was shorter than the 5.06 min observed in Ctrl/Sham mice (Holm-adjusted P = 0.0130). The persistence curves estimated that the probability of an inactivity bout lasting beyond 5 min was 36.9% in SD/tUS mice and 20.5% in SD/Sham mice (Figure 5G). Short inactivity bouts remained common in both groups. Supplementary Figure S13 shows daily within-SD contrasts. Supplementary Table S3 reports all estimates and 95% CIs.

Mean body weight initially fell in both SD groups and approached the Day 10 baseline earlier in SD/tUS than in SD/Sham mice. Percentage increases in food intake from baseline were larger in SD than in Ctrl groups (Supplementary Figure S14).

### 3.5. Object recognition and open-field behavior

In the separate behavioral-test cohort, the mean NORT discrimination index (DI) was 0.1869 in Ctrl/Sham mice, −0.0067 in SD/Sham mice, and 0.360 in SD/tUS mice. The within-SD difference was nonsignificant after Holm adjustment (P = 0.119), as was the housing condition × treatment condition interaction (P = 0.1459; Figure 6A). Figure 6D–G and Supplementary Figure S15 show representative exploration maps and individual exploration times for familiar and novel objects. Supplementary Table S4 lists all four contrasts per outcome.

**Figure 6.**
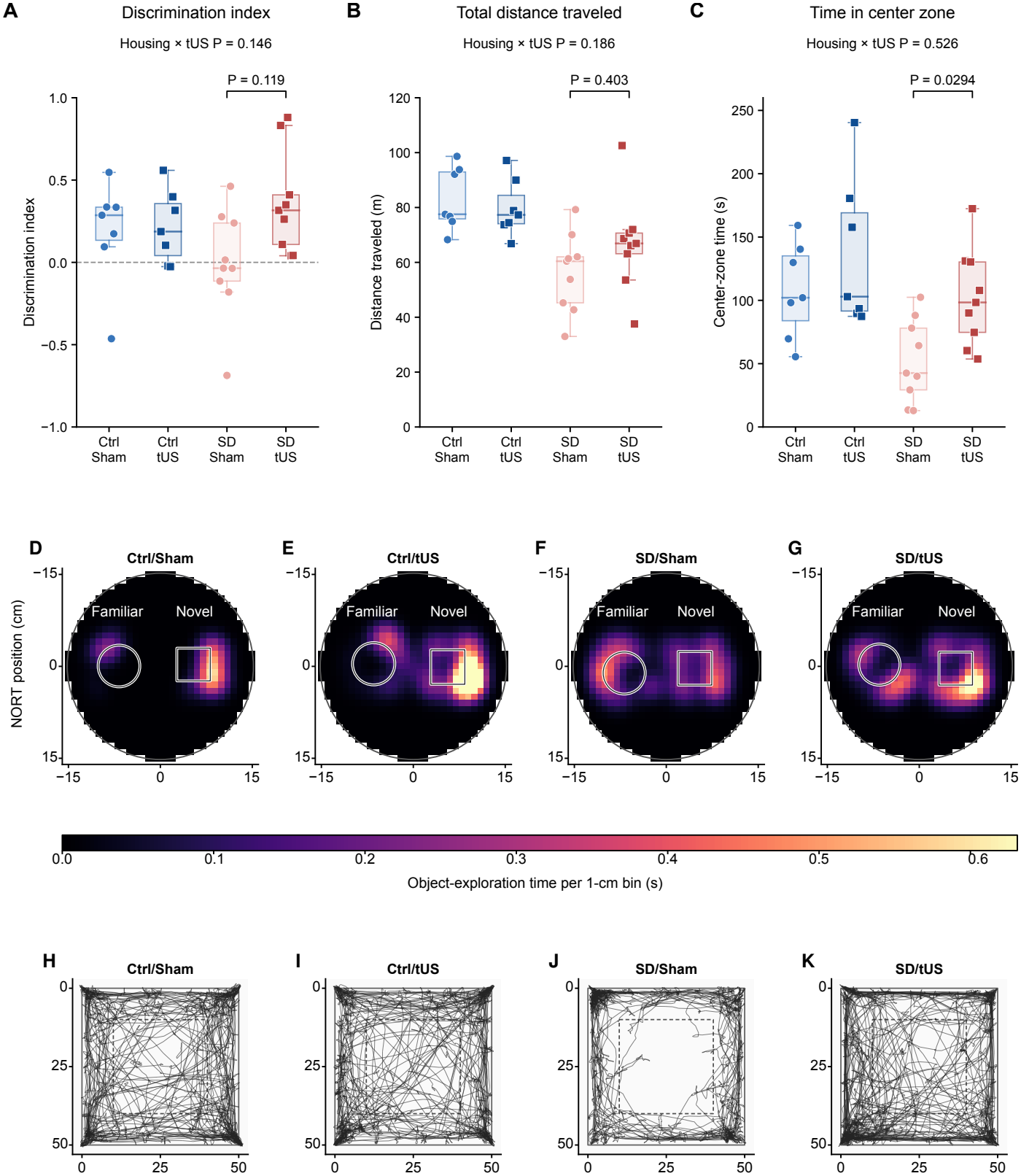
Novel-object-recognition and open-field behavior. (A) NORT discrimination index, 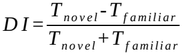. (B) OFT total distance. (C) OFT center time. (D–G) Illustrative NORT exploration heatmaps over the 300-s cumulative observation window, with Gaussian smoothing for display and a shared color scale. The outlines mark the object locations used for image analysis. (H–K) Illustrative OFT trajectories. Coordinates are in centimeters and the dashed square marks the central 30 × 30 cm zone. In A–C, points show individual mice; boxes span the 25th–75th percentiles with median lines, whiskers extend to the most extreme observations within 1.5 times the interquartile range. All observations are shown, including those beyond the whiskers. The same 32 mice contributed to both tests (Ctrl/Sham and Ctrl/tUS, n = 7 each; SD/Sham and SD/tUS, n = 9 each). Within each group, the NORT heatmap and OFT trajectory show different mice. In A–C, the bracket shows SD/tUS versus SD/Sham. Its two-sided HC3 P value is Holm-adjusted across all four contrasts for that outcome. The P value above each panel is for the housing × tUS interaction. Supplementary Table S4 reports all four contrasts.

SD/Sham mice spent less time in the OFT center and traveled less than Ctrl/Sham mice. SD/tUS mice spent longer in the center than SD/Sham mice (102.08 vs 52.36 s; mean difference, 49.73 s; 95% CI, 13.47 to 85.98 s; Holm-adjusted P = 0.0294; Figure 6C). Total distance did not differ significantly within SD (Holm-adjusted P = 0.403; Figure 6B). Neither outcome showed a significant housing × treatment interaction (center time and distance, P = 0.5261 and 0.1858, respectively). Figure 6H–K shows representative trajectories. Across all 32 mice, longer OFT center time on Day 17 was associated with lower activity-resumption probability (ρ = −0.616), longer median inactivity bouts (ρ = 0.383), and fewer switches per hour (ρ = −0.554) during the preceding two dark phases (Supplementary Figure S16D–F). Higher running intensity was also associated with longer center time (ρ = 0.449), and more wheel rotations with greater OFT distance (ρ = 0.445; Supplementary Figure S16J,K). None of the five wheel measures showed a clear association with NORT DI (Supplementary Figure S16A–C, G, H).

## 4. Discussion

During ongoing SD, mice receiving repeated motor-cortex-targeted tUS had a lower probability of resuming wheel running and longer median inactivity bouts than Sham mice. For both measures, the housing condition × treatment interaction indicated that the association with repeated tUS differed between housing conditions. In the persistence of pauses between runs, SD/tUS mice more closely resembled Ctrl/Sham mice than did SD/Sham mice (Figure 5D, F). These longer pauses (Figure 5G) may have provided opportunities for sustained rest while the sleep-disrupting conditions continued.

With the transducer in this position, the targeted motor cortex was predicted to be the main site of exposure through the skull. In water, on-axis time-averaged intensity peaked 1.4 mm beyond the collimator outlet, where the lateral −6 dB intensity widths were approximately 1.6 mm along both X and Y (Supplementary Figure S1D–F). Across ten skull models calibrated to these measurements, the primary and secondary motor areas accounted for a mean of 73.4% of the atlas-labeled predicted exposure volume. The predicted field also overlapped part of the caudoputamen in all ten models, a subcortical extension relevant to interpreting the behavioral response (Figure 2B–E, Supplementary Figure S2). The predicted peak rise in brain temperature during one session averaged approximately 0.38°C across skull models and sonication orders (Supplementary Figure S3). This falls below the 2°C rise that ITRUSST uses as one criterion for nonsignificant thermal risk in human tUS at 200–800 kHz [32]. In mice undergoing remyelination, Olmstead et al. delivered 1.09-MHz ultrasound in 30-min daily sessions for five days and found no histologically evident neuronal injury at the sonication site in any of the seven mice examined [33]. Our temperature estimates and their histological findings together suggest that overt thermal injury to the brain parenchyma was unlikely under our exposure conditions.

The motor cortex, where the predicted exposure was concentrated, also contributes to the initiation of running. In awake mice, motor-cortical neurons show different firing patterns during wheel running and nonrunning periods [20]. In a goal-directed treadmill task, Zhang et al. found that transient pharmacological inactivation of the secondary motor area (MOs) reduced the frequency of running bouts and prolonged pauses [34]. Longer pauses were also a feature of the present study. SD/tUS and SD/Sham mice ran at similar intensities, but SD/tUS mice were less likely to resume running after a pause (Figure 5C, D), linking repeated tUS to the transition from inactivity to spontaneous running. In mouse primary motor cortex, tUS increased or decreased motor-evoked potentials depending on the burst pattern, and these changes persisted after sonication [7]. At the 40-Hz pulse repetition frequency used here, cortical neurons in awake mice showed both depolarizing and hyperpolarizing responses [6]. Together with the predicted exposure distribution, these findings raise the possibility that a change in motor-cortical responsiveness after stimulation contributed to the lower probability of resuming running in SD mice.

In the independent EEG/EMG cohort that received no tUS, SD mice spent more time asleep than Ctrl mice, yet switched more frequently between sleep and wakefulness and had higher absolute EEG δ power during wakefulness (Figure 3A, B, Supplementary Figure S8A). In mice, slow-wave activity during quiet wakefulness increases as acute sleep disruption continues [35]. Under chronic sleep fragmentation, mice return to sleep sooner after awakening even when total sleep time is largely preserved [13]. Sleep pressure may thus have remained high in our SD mice despite the greater time spent asleep. Matching wheel activity to EEG/EMG in this cohort showed that wheel inactivity included both sleep and wakefulness (Figure 3C, D, Supplementary Figure S7). Some of the longer pauses in the intervention cohort may likewise have provided opportunities to fall asleep or remain asleep.

In the separate behavioral-testing cohort, the mean NORT discrimination index was higher in SD/tUS than in SD/Sham mice, although the difference was not significant after Holm adjustment (Figure 6A). The object-recognition deficit reported in C3H/HeN mice confined to a wheel to avoid water [23] was not evident in our SD mice. Our findings therefore offer limited insight into the preservation of recognition memory after repeated tUS reported in 5xFAD mice [36] or the relationship between sleep continuity and memory retention [14]. In the OFT, SD/tUS mice spent more time in the center than SD/Sham mice, with no significant difference in total distance traveled between the two groups (Figure 6B, C). This pattern of exploration is consistent with reduced anxiety-like behavior. The housing condition × treatment interaction for center time was not significant, so this interpretation applies to the comparison within SD.

The two measures of running pauses that changed with repeated tUS under SD in the wheel cohort also related to exploration of a novel environment the next day in the separate behavioral-testing cohort. Across the 32 mice from all four groups, a lower probability of resuming running and longer median inactivity bouts during the two dark phases before testing were each associated with more time in the OFT center the following day (Supplementary Figure S16D–F). Running intensity was also positively associated with center time (Supplementary Figure S16J). The NORT discrimination index showed no clear association with these wheel measures (Supplementary Figure S16A–C, G, H). Both how long mice paused between runs and how vigorously they ran captured individual differences in subsequent center exploration.

Exploration of the center of a novel arena also relates to sleep conditions before testing. In the chronic sleep fragmentation model of Xie et al., mice spent less time in the OFT center despite traveling farther overall [37]. After acute social defeat stress, Feng et al. found that mice given an opportunity to sleep spent more time in the center than mice kept awake for the same period, with no significant difference in total distance traveled [38]. The longer pauses observed here may have afforded opportunities for sustained rest during the dark phase that, in turn, were relevant to center exploration the following day.

Pauses between runs may provide a measure of spontaneous behavior after repeated tUS in mice living under sleep-disrupting conditions. In freely behaving mice, EEG/EMG detection of NREM sleep has been used to trigger ultrasound stimulation of the medial prefrontal cortex [39]. Recording wheel activity and EEG/EMG concurrently during the dark phase after treatment in the intervention mice would establish how much of each prolonged pause is spent in quiet wakefulness and, when mice fall asleep, how long sleep persists.

## Declarations

CRediT: Yutaro Hazama: Conceptualization, Methodology, Software, Data curation, Visualization, Funding acquisition, Investigation, Formal analysis, Writing – original draft, Supervision. Kodai Kawata: Investigation, Formal analysis. Kaede Takemura: Investigation, Formal analysis. Itsuki Sawada: Investigation. Reon Konaka: Investigation, Formal analysis. Karin Shimizu: Investigation, Formal analysis. Riki Sato: Investigation. Shuhei Iida: Investigation. Maya Goto: Formal analysis. Minto Arimura: Investigation. Sakura Shinozawa: Investigation. Ayano Matsushita: Investigation. Soichiro Kimura: Writing – review & editing, Supervision. Hideo Ueda: Writing – review & editing, Supervision.

## Data availability

Wheel-activity data, EEG/EMG recordings and derived measures, and behavioral data from the novel object recognition test (NORT) and open-field test (OFT) are available on Zenodo (https://doi.org/10.5281/zenodo.23996096). A README and figure-to-file map accompany the data. Author-created data are licensed under CC BY 4.0.

## Code availability

Analysis code and utilities for reproducing the supplied numerical summaries and figure display data are available on Zenodo (https://doi.org/10.5281/zenodo.23996126). The README describes the dependencies, instructions, and numerical results covered by the code. Author-created code is licensed under MIT.

Declaration of generative AI and AI-assisted technologies in the manuscript preparation process: The authors used ChatGPT and Codex (OpenAI) to assist with English translation and editing of parts of the manuscript and with the development of analysis and visualization code. The authors reviewed and revised the AI-assisted material and take full responsibility for the content of the published article.

## References

[1] Blackmore J, Shrivastava S, Sallet J, Butler CR, Cleveland RO. Ultrasound Neuromodulation: A Review of Results, Mechanisms and Safety. Ultrasound Med Biol. 2019;45(7):1509–1536. 10.1016/j.ultrasmedbio.2018.12.015

[2] Pellow C, Pichardo S, Pike GB. A systematic review of preclinical and clinical transcranial ultrasound neuromodulation and opportunities for functional connectomics. Brain Stimul. 2024;17(4):734–751. 10.1016/j.brs.2024.06.005

[3] Tufail Y, Matyushov A, Baldwin N, Tauchmann ML, Georges J, Yoshihiro A, et al. Transcranial pulsed ultrasound stimulates intact brain circuits. Neuron. 2010;66(5):681–94. 10.1016/j.neuron.2010.05.008

[4] Lee W, Croce P, Margolin RW, Cammalleri A, Yoon K, Yoo SS. Transcranial focused ultrasound stimulation of motor cortical areas in freely-moving awake rats. BMC Neurosci. 2018;19(1):57. 10.1186/s12868-018-0459-3

[5] Wang X, Yan J, Wang Z, Li X, Yuan Y. Neuromodulation Effects of Ultrasound Stimulation Under Different Parameters on Mouse Motor Cortex. IEEE Trans Biomed Eng. 2020;67(1):291–297. 10.1109/tbme.2019.2912840

[6] Bortz E, San Antonio E, Sherman J, Tseng HA, Raiff L, Han X. Transcranial ultrasound stimulation modulates neuronal membrane potentials across broad timescales in the awake mammalian brain. Brain Stimul. 2025;18(6):1726–1740. 10.1016/j.brs.2025.09.004

[7] Kim HJ, Phan TT, Lee K, Kim JS, Lee SY, Lee JM, et al. Long-lasting forms of plasticity through patterned ultrasound-induced brainwave entrainment. Sci Adv. 2024;10(8):eadk3198. 10.1126/sciadv.adk3198

[8] Zhou H, Niu L, Xia X, Lin Z, Liu X, Su M, et al. Wearable Ultrasound Improves Motor Function in an MPTP Mouse Model of Parkinson’s Disease. IEEE Trans Biomed Eng. 2019;66(11):3006–3013. 10.1109/TBME.2019.2899631

[9] Takahashi N, Moberg S, Zolnik TA, Catanese J, Sachdev RNS, Larkum ME, et al. Thalamic input to motor cortex facilitates goal-directed action initiation. Curr Biol. 2021;31(18):4148–4155.e4. 10.1016/j.cub.2021.06.089

[10] Sun G, Yu C, Cai R, Li M, Fan L, Sun H, et al. Neural representation of self-initiated locomotion in the secondary motor cortex of mice across different environmental contexts. Commun Biol. 2025;8(1):725. 10.1038/s42003-025-08169-7

[11] Hoffman DJ, Newland MC. A microstructural analysis distinguishes motor and motivational influences over voluntary running in animals chronically exposed to methylmercury and nimodipine. Neurotoxicology. 2016;54:127–139. 10.1016/j.neuro.2016.04.009

[12] Ananthasubramaniam B, Meijer JH. Regulation of Rest, Rather Than Activity, Underlies Day-Night Activity Differences in Mice. Front Physiol. 2020;11:268. 10.3389/fphys.2020.00268

[13] Baud MO, Magistretti PJ, Petit JM. Sustained sleep fragmentation induces sleep homeostasis in mice. Sleep. 2015;38(4):567–579. 10.5665/sleep.4572

[14] Rolls A, Colas D, Adamantidis A, Carter M, Lanre-Amos T, Heller HC, et al. Optogenetic disruption of sleep continuity impairs memory consolidation. Proc Natl Acad Sci U S A. 2011;108(32):13305–10. 10.1073/pnas.1015633108

[15] Kuhn M, Wolf E, Maier JG, Mainberger F, Feige B, Schmid H, et al. Sleep recalibrates homeostatic and associative synaptic plasticity in the human cortex. Nat Commun. 2016;7:12455. 10.1038/ncomms12455

[16] Salehinejad MA, Ghanavati E, Reinders J, Hengstler JG, Kuo MF, Nitsche MA. Sleep-dependent upscaled excitability, saturated neuroplasticity, and modulated cognition in the human brain. eLife. 2022;11:e69308. 10.7554/eLife.69308

[17] Wang T, Wang X, Tian Y, Gang W, Li X, Yan J, et al. Modulation effect of low-intensity transcranial ultrasound stimulation on REM and NREM sleep. Cereb Cortex. 2023;33(9):5238–5250. 10.1093/cercor/bhac413

[18] Wu Y, Wang H, Wang M, Ren P, Zhang S. Modulation of rapid eye movement sleep deprivation via transcranial ultrasound stimulation of the basal forebrain in mice. Ultrasonics. 2027;169:108282. 10.1016/j.ultras.2026.108282

[19] Vyazovskiy VV, Tobler I. The temporal structure of behaviour and sleep homeostasis. PLoS One. 2012;7(12):e50677. 10.1371/journal.pone.0050677

[20] Fisher SP, Cui N, McKillop LE, Gemignani J, Bannerman DM, Oliver PL, et al. Stereotypic wheel running decreases cortical activity in mice. Nat Commun. 2016;7:13138. 10.1038/ncomms13138

[21] Miyazaki K, Itoh N, Ohyama S, Kadota K, Oishi K. Continuous exposure to a novel stressor based on water aversion induces abnormal circadian locomotor rhythms and sleep-wake cycles in mice. PLoS One. 2013;8(1):e55452. 10.1371/journal.pone.0055452

[22] Minakawa EN, Miyazaki K, Maruo K, Yagihara H, Fujita H, Wada K, et al. Chronic sleep fragmentation exacerbates amyloid β deposition in Alzheimer’s disease model mice. Neurosci Lett. 2017;653:362–369. 10.1016/j.neulet.2017.05.054

[23] Sakamoto K, Higo-Yamamoto S, Egi Y, Miyazaki K, Oishi K. Memory dysfunction and anxiety-like behavior in a mouse model of chronic sleep disorders. Biochem Biophys Res Commun. 2020;529(2):175–179. 10.1016/j.bbrc.2020.05.218

[24] Martin E, Aubry JF, Schafer M, Verhagen L, Treeby B, Pauly KB. ITRUSST consensus on standardised reporting for transcranial ultrasound stimulation. Brain Stimul. 2024;17(3):607–615. 10.1016/j.brs.2024.04.013

[25] Devine J, Vidal-García M, Liu W, Neves A, Lo Vercio LD, Green RM, et al. MusMorph, a database of standardized mouse morphology data for morphometric meta-analyses. Sci Data. 2022;9(1):230. 10.1038/s41597-022-01338-x

[26] Hikishima K, Komaki Y, Seki F, Ohnishi Y, Okano HJ, Okano H. In vivo microscopic voxel-based morphometry with a brain template to characterize strain-specific structures in the mouse brain. Sci Rep. 2017;7(1):85. 10.1038/s41598-017-00148-1

[27] Wang Q, Ding SL, Li Y, Royall J, Feng D, Lesnar P, et al. The Allen Mouse Brain Common Coordinate Framework: A 3D Reference Atlas. Cell. 2020;181(4):936–953.e20. 10.1016/j.cell.2020.04.007

[28] Treeby BE, Cox BT. k-Wave: MATLAB toolbox for the simulation and reconstruction of photoacoustic wave fields. J Biomed Opt. 2010;15(2):021314. 10.1117/1.3360308

[29] Treeby BE, Jaros J, Rendell AP, Cox BT. Modeling nonlinear ultrasound propagation in heterogeneous media with power law absorption using a k-space pseudospectral method. J Acoust Soc Am. 2012;131(6):4324–36. 10.1121/1.4712021

[30] Pennes HH. Analysis of tissue and arterial blood temperatures in the resting human forearm. J Appl Physiol. 1948;1(2):93–122. 10.1152/jappl.1948.1.2.93

[31] Lueptow LM. Novel Object Recognition Test for the Investigation of Learning and Memory in Mice. J Vis Exp. 2017;(126):55718. 10.3791/55718

[32] Aubry JF, Attali D, Schafer ME, Fouragnan E, Caskey CF, Chen R, et al. ITRUSST consensus on biophysical safety for transcranial ultrasound stimulation. Brain Stimul. 2025;18(6):1896–1905. 10.1016/j.brs.2025.10.007

[33] Olmstead TA, Chiarelli PA, Griggs DJ, McClintic AM, Myroniv AN, Mourad PD. Transcranial and pulsed focused ultrasound that activates brain can accelerate remyelination in a mouse model of multiple sclerosis. J Ther Ultrasound. 2018;6:11. 10.1186/s40349-018-0119-1

[34] Zhang CL, Koukouli F, Allegra M, Ortiz C, Kao HL, Maskos U, et al. Inhibitory control of synaptic signals preceding locomotion in mouse frontal cortex. Cell Rep. 2021;37(8):110035. 10.1016/j.celrep.2021.110035

[35] Grønli J, Clegern WC, Schmidt MA, Nemri RS, Rempe MJ, Gallitano AL, et al. Sleep homeostatic and waking behavioral phenotypes in Egr3-deficient mice associated with serotonin receptor 5-HT2 deficits. Sleep. 2016;39(12):2189–2199. 10.5665/sleep.6324

[36] Yoo SS, Reddy A, Carroll W, Ploypradith K. Repeated application of transcranial ultrasound maintains spatial and recognition memory in 5xFAD mice with reduction of amyloid-β burden. PLoS One. 2025;20(11):e0336114. 10.1371/journal.pone.0336114

[37] Xie Y, Ba L, Wang M, Deng SY, Chen SM, Huang LF, et al. Chronic sleep fragmentation shares similar pathogenesis with neurodegenerative diseases: Endosome-autophagosome-lysosome pathway dysfunction and microglia-mediated neuroinflammation. CNS Neurosci Ther. 2020;26(2):215–227. 10.1111/cns.13218

[38] Feng X, Zhao HY, Shao YJ, Lou HF, Zhu LY, Duan S, et al. Anxiolytic effect of increased NREM sleep after acute social defeat stress in mice. Neurosci Bull. 2020;36(10):1137–1146. 10.1007/s12264-020-00473-y

[39] Jo Y, Lee SM, Jung T, Park G, Lee C, Im GH, et al. General-purpose ultrasound neuromodulation system for chronic, closed-loop preclinical studies in freely behaving rodents. Adv Sci (Weinh). 2022;9(34):e2202345. 10.1002/advs.202202345

