## Supplementary Information for "Repeated transcranial ultrasound is associated with altered temporal organization of spontaneous wheel activity during sleep disruption in mice"

#### Supplementary Methods

##### S1. Animals and experimental housing

###### S1.1. Animals and SD conditions

The analyses included 79 male C3H/HeNSlc mice across three independent cohorts. The independent EEG/EMG cohort underwent recording without tUS or sham to characterize the SD model. Mice in the wheel-activity and NORT/OFT intervention cohorts received tUS or sham. Mice were purchased from Japan SLC, Inc. and were approximately 8 weeks old upon arrival. They were housed individually under a 12-h light/12-h dark cycle, with lights on at 07:00 (Zeitgeber time [ZT] 0) and off at 19:00 (ZT12), at 22 ± 2 °C and 55 ± 10% relative humidity. Food and water were available ad libitum. All animal procedures were approved by the Institutional Animal Care and Use Committee of Josai University under protocols JU 22097, JU 23020, JU 24020, JU 25018, and JU 26018 and were conducted in accordance with the Josai University Regulations for Animal Experiments.

The wheel-activity and behavioral-testing intervention cohorts were kept separate because NORT and OFT required removing mice from their wheel cages, interrupting continuous wheel monitoring and potentially altering subsequent spontaneous activity.

We adapted the wheel-housing procedure of Minakawa et al. [1]. Across all three cohorts, mice first acclimated for 10 days in individual cages with free access to the floor and running wheel. On Day 10 at ZT8, mice assigned to the sleep-disruption (SD) condition were transferred to wheel-confinement cages, in which they were required to remain on the wheel without access to a conventional cage floor. In contrast, control (Ctrl) mice remained in their original cages, retaining free access to both the floor and wheel. Wood-chip bedding was placed in the bottom tray of the SD apparatus, and no water-avoidance procedure was employed. Food and drinking water remained available ad libitum in both conditions. The assigned housing conditions were maintained through the end of recording or behavioral testing in each cohort. Behavioral testing on Day 17, seven days after SD onset, was timed with reference to Sakamoto et al., who conducted NORT or OFT after one week of sleep disruption [2]. In the two tUS/sham intervention cohorts, SW-15S and SW-15-SD cages (Melquest, Toyama, Japan) were used for the Ctrl and SD conditions, respectively. Housing for the EEG/EMG cohort is described in S1.2.

###### S1.2. EEG/EMG-compatible wheel housing

The independent EEG/EMG cohort was housed in RWC-15 (Ctrl) or RWC-15-SD (SD) cages (Melquest, Toyama, Japan). These cages had the same basic geometry as the corresponding SW-series cages used in the intervention cohorts, except that one side of the running wheel was open to accommodate the EEG/EMG recording cable. A custom guard was installed beneath the wheel to prevent access to the space below or beyond it and reduce cable-entanglement risk during tethered recording. The guard was designed in Autodesk Fusion 360 and 3D-printed. This modification was used only in the independent EEG/EMG cohort and was not applied to either tUS/sham intervention cohort.

##### S2. Ultrasound apparatus and procedures

Transcranial ultrasound stimulation was delivered using a custom-made planar piezoelectric transducer (20-mm diameter) and matching box manufactured by Honda Electronics Co., Ltd. (Toyohashi, Aichi, Japan). The transducer was equipped with a custom collimator comprising a 2-mm-diameter outlet and a 10-mm-long tapered section. The collimator was fabricated from Clear Resin using a Form 3B three-dimensional printer (Formlabs, Somerville, MA, USA), and the transducer assembly was mounted in an F44 slide lens holder

(Suruga Seiki, Japan). The system used neither geometric nor electronic focusing. The driving waveform was generated using a multifunction generator (WF1973; NF Corporation, Yokohama, Japan) and amplified using a high-speed bipolar amplifier (HSA4101; NF Corporation). Acoustic exposure parameters and reporting details are provided in Supplementary Table S1, Part A, following the ITRUSST consensus recommendations [3].

Scalp hair was removed in both tUS and sham groups on Day 10 using electric clippers followed by depilatory cream (Epilat; Kracie, Japan). Hair removal was not repeated during stimulation. Mice assigned to active tUS were anesthetized with isoflurane (5% for induction and 1.5–2.5% for maintenance) and positioned in a stereotaxic apparatus (Model 68801; RWD Life Science). Before head fixation, the collimator tip was aligned with the interaural-zero reference of the stereotaxic apparatus. After fixation, the incisor-bar height was adjusted using the stereotaxic atlas of Franklin and Paxinos [4] so that bregma and lambda were approximately level. Nominal bilateral motor-cortex targets were defined relative to bregma at anteroposterior +1.5 mm and mediolateral ±1.5 mm. Ultrasound gel (Pro Jelly, Normal Type; Jex, Japan) was used for acoustic coupling: the tapered section of the collimator was filled with gel, and additional gel was applied to the scalp. During anesthesia, mice in both groups were kept warm using an incandescent lamp. After tUS, residual scalp gel was wiped off.

Ultrasound was delivered at a carrier frequency of 1.078 MHz with a pulse repetition frequency of 40 Hz. Each 25-ms pulse cycle comprised 12.5 ms of ultrasound emission followed by 12.5 ms without emission, corresponding to a 50% duty cycle. In each daily session, the left and right motor-cortex targets were stimulated alternately in six consecutive 5-min blocks, for a total of 30 min of stimulation blocks excluding repositioning time, 15 min per hemisphere. At 50% duty cycle, ultrasound was on for 15 min across the two hemispheres combined. Stimulation was administered once daily on five consecutive days (Days 13–17), between ZT2 and ZT4, while mice remained under their assigned Ctrl or SD conditions.

Mice assigned to the sham groups received isoflurane anesthesia under the same conditions as mice undergoing active tUS (5% for induction and 1.5–2.5% for maintenance), with matched anesthesia duration and the same daily treatment schedule (Days 13–17, ZT2–ZT4). During sham procedures, mice were placed in a separate box without stereotaxic head fixation. No ultrasound was delivered, and neither a transducer nor ultrasound gel was applied to the scalp.

#### S3. Acoustic measurements and modeling

##### S3.1. Acoustic measurements and reference intensity

Free-field acoustic measurements were performed in water using a calibrated needle hydrophone (HNR-0500; Onda Corporation, Sunnyvale, CA, USA). The three-axis scanning tank was based on the design described by Clinard et al. [5]. The measured carrier frequency was approximately 1.078 MHz, with a pulse repetition frequency of 40 Hz and nominal ON and OFF durations of 12.5 ms each. Recorded hydrophone voltages were baseline-corrected and converted to acoustic pressure using the effective calibration sensitivity [6]. Peak negative pressure was calculated as the magnitude of the most negative pressure within each retained pulse, then averaged across pulses at each sampled position. Its maximum within the sampled XY plane was 0.300 MPa, at the same sampled position as the maximum temporal-average intensity. Local temporal-average intensity (I_TA) was calculated by integrating the squared acoustic pressure over complete pulse-repetition intervals, including both ON and OFF periods, and dividing by the pulse period and acoustic impedance of water. The final XY, XZ, and YZ measurement maps contained 3,147 retained cells in total. The XY plane contained 1,845 sampled positions. Figure 2A shows a 3 × 3 mm portion of this plane. The maximum I_TA within the sampled XY plane at $Z = -1.4$ mm was 1.193585 $W/cm^2$ and was used as the water reference for reporting the intracranial peak-intensity ratio. This reference represented the maximum within the measured plane, rather than an exhaustively determined three-dimensional maximum. Measured pulse-average and temporal-average intensities were retained separately without forcing their ratio to equal the nominal duty cycle. Additional water-field maps and line profiles are shown in Supplementary Figure S1. In

Supplementary Figure S1, X = 0 mm and Y = 0 mm denote the collimator center; scanner Z = 0 mm was aligned with the collimator outlet plane during hydrophone measurements. For each profile, the −3 and −6 dB intensity thresholds were 50.12% and 25.12%, respectively, of its maximum $I_{TA}$. Widths were determined between the linearly interpolated crossings bounding the contiguous interval containing that maximum. Complete widths were reported only when both crossings fell within the measured range. The three planes were scanned sequentially without drift correction. Analyses included all final measurement positions, including those flagged during quality control. Grid positions are spatial samples, not independent experimental replicates. These water measurements do not constitute an exhaustive three-dimensional scan or a direct measurement of intracranial exposure.

**Definition**

$$I_{\text{TA,water}}(r)=\frac{1}{T_p}\int_0^{T_p}\frac{p(r,t)^2}{\rho_w c_w}\,dt$$

Here, p is acoustic pressure (Pa), t is time (s), $T_p$ is the pulse period (s), $\rho_w$ is water density (kg/m³), and $c_w$ is sound speed in water (m/s). The equation yields W/m² in SI units. Intensities are reported in W/cm².

##### S3.2. CT-derived anatomy and approximate atlas registration

Anatomical models were constructed using CT-derived skull geometries from ten independent postnatal day 90 (P90) male C3H/HeJ mice obtained through the MusMorph/FaceBase Strain Comparison resource [7,8]. Because individual brain MR images were unavailable for these skulls, intracranial brain and effective cerebrospinal fluid (CSF) compartments were estimated using a common C3H/He population MRI template [9]. Anatomical labels, including the primary motor area (MOp), secondary motor area (MOs), and caudoputamen (CP), were approximately mapped from the Allen Mouse Brain Common Coordinate Framework version 3 (CCFv3) [10]. Atlas labels were mapped to the common MRI template using brain-shape-based scaling and translation. For each CT skull, a local coordinate system was defined using dataset-provided bregma, lambda and paired cranial landmarks. The MRI template was positioned in this coordinate system and translated dorsoventrally to match the CT-estimated intracranial roof. The resulting tissue compartments and atlas labels were therefore treated as model-dependent anatomical approximations, not verified individual boundaries. Overlap with atlas-assigned motor areas described the predicted spatial distribution of acoustic exposure. It was not interpreted as evidence of selective neuronal activation.

##### S3.3. Acoustic propagation and source calibration

Acoustic propagation was simulated separately for the left and right stimulation positions using a three-dimensional, heterogeneous, linear compressional-wave model implemented with a k-space pseudospectral method [11,12]. The computational grid spacing was 120 μm, with 128 temporal samples per acoustic cycle at the measured carrier frequency of 1.078 MHz. Density, sound speed, and the attenuation coefficient at the carrier frequency were spatially uniform within each modeled tissue compartment, with the same tissue-specific values used across all ten skull models. Attenuation was represented by a Stokes-type frequency-squared loss model matched to the specified attenuation coefficient at the carrier frequency; attenuation was treated as absorption for heat-source calculations. Using the principle of acoustic holography [13], we constructed a complex field at 1,845 hydrophone positions 1.4 mm from the collimator outlet using the square root of measured temporal-average intensity and carrier phase relative to the drive monitor. The XY maximum defined the transverse origin. The forward-propagating components were back-propagated to an equivalent numerical source plane 2.64 mm proximal to the outlet using a zero-padded angular spectrum and a 6-mm source radius. To calibrate source amplitude against the measured axial pressure profile in water [14], we used the XZ scan at 31 depths from 0 to 6 mm along the profile shown in Supplementary Figure S1E. The YZ scan provided a repeat measurement along the same line (Supplementary Figure S1F). At each depth, carrier amplitudes from sinusoidal fits over 20 cycles starting 160 μs after nominal arrival were averaged across pulses. The ratio of measured to modeled peak pressure along the XZ line (0.26051 and 0.26328 MPa, respectively) gave a calibration factor of 0.98946. For both sides of all ten skulls, we scaled the linear model outputs after simulation, multiplying pressure by this factor and intensity, heat deposition, and temperature rise by its square (0.97903). The model did not include elastic shear-wave propagation in the skull. Material parameters and their underlying transfer assumptions are summarized in Supplementary Table S1, Part B.

S3.4. Acoustic intensity and atlas-based exposure metrics

Local temporal-average acoustic intensity was estimated from the simulated peak-amplitude pressure phasor as $I_{TA}(r)=D|\hat{p}(r)|^2/[2\rho(r)c(r)]$, where $D=0.5$ was the duty cycle and $\rho$ and $c$ were the local tissue density and sound speed. The duty cycle was applied once. For each stimulation side, the spatial-peak temporal-average intensity was defined as the maximum of this pressure-derived quantity within the physical brain and effective cerebrospinal fluid compartments, excluding other tissues and perfectly matched layers. The water-peak intensity ratio was calculated by dividing this intracranial peak by the measured maximum temporal-average intensity in the sampled water XY plane. This ratio compared two spatial peaks and was not interpreted as transmitted acoustic power or energy.

For each stimulation side, the pressure-amplitude peak was identified within the combined region comprising physical brain and effective cerebrospinal fluid. The peak amplitude was the maximum modulus of the complex pressure phasor in this region and ranged from 0.205 to 0.244 MPa across the 20 side-specific fields. In the linear harmonic model, this amplitude also represents the magnitude of the negative pressure peak; it is a model prediction rather than a measured intracranial waveform. Voxels with pressure amplitudes at least −6 dB relative to that side-specific peak were identified within the same combined region, and the six-neighbor connected component containing the peak was retained. Using the approximate Allen CCFv3 registration, the share assigned to MOp, MOs, caudoputamen (CP), or any other annotated category was calculated by dividing the number of voxels assigned to that category within the retained component by the total number of annotated voxels in the component, including annotated fiber tracts. The unannotated fraction was calculated separately using all voxels in the component as its denominator. These metrics described the atlas-assigned composition of the predicted exposure region, not the proportion of each anatomical structure exposed or the selectivity of neural activation. Side-specific metrics were averaged within each of the ten skull models before group summaries were calculated. Skull models, rather than individual sides, were the independent units.

**Definitions**

$$I_{TA}(r)=\frac{D|\hat{p}(r)|^2}{2\rho(r)c(r)}, D=0.5$$

$$|\hat{p}(r)|\geq 10^{-6/20}|\hat{p}|_{max}\approx 0.5012|\hat{p}|_{max}$$

$$\text{Overlap}_{MOp}(\%)=100\frac{N_{MOp}}{N_{\text{all annotated}}}$$

The overlap denominator is the number of atlas-annotated voxels **within the peak-connected component**, rather than the voxel count of the anatomical structure as a whole.

Regional coverage was calculated separately from exposure-region composition. The numerator was the volume of the union of the retained left/right components within each region on the 120-µm acoustic grids. The denominator was the entire bilateral region volume, including atlas descendants, from the 25-µm Allen CCF annotation multiplied by the absolute determinant of the saved atlas-to-skull affine transform. This denominator was not limited to the local acoustic display volume. The ratio was therefore an approximate three-dimensional volume fraction across different grids. One coverage estimate was obtained per skull and summarized across ten skulls; this was not a measure of neuronal activation. Both volumes were expressed in $mm^3$ and regional coverage as a percentage of the bilateral region volume.

##### S3.5. Cohort aggregation, uncertainty estimates, and visualization

Acoustic intensity and exposure-composition metrics were calculated separately for each stimulation side and then averaged within each skull model. Cohort summaries were expressed as mean ± sample standard deviation across ten independent skulls. We estimated 95% confidence intervals for cohort means using the percentile bootstrap, resampling the ten skulls 50,000 times with left- and right-sided predictions kept together [15]. These intervals quantified sampling variability across the modeled skulls and did not incorporate uncertainty in tissue properties or anatomical registration.

For visualization, the native three-dimensional temporal-average intensity fields and the peak-connected pressure-amplitude components defined at −6 dB in S3.4 (ITRUSST pressure-decibel convention [3]) were mapped to display coordinates. Intensity maps used linear interpolation, whereas anatomical labels and component masks used nearest-neighbour interpolation. Quantitative metrics were calculated from the native fields and masks before display interpolation.

Dorsal views (Figure 2B) used a maximum-intensity projection; a projected component mask was positive wherever the component occurred at any depth. Coronal views (Figure 2C) were extracted at AP +1.5 mm. For dorsal and coronal views, the independently simulated left/right intensity fields were combined by their within-skull maximum, and component masks by spatial union. Sagittal views (Figure 2D) at ML −1.5/+1.5 mm used the corresponding left/right field and component. These combinations did not represent simultaneous irradiation or summed intensity. Anatomical outlines and intracranial support were derived from each skull model's existing approximate registration. All individual maps used identical axes, with coronal and sagittal views spanning 0–5.0 mm ventral to the bregma plane, and a common intensity scale of 0–1.3 W/cm$^2$. Intensities below 0.0455 W/cm$^2$ were hidden only for visualization; this threshold did not define the three-dimensional components or imply zero exposure.

Three-dimensional views of representative skull C3H_10M (Figure 2E) were rendered in Blender 5.2.1 LTS using Python scripts. Exposure surfaces were generated from the outer faces of occupied voxels in the native 120-µm, side-specific peak-connected −6-dB pressure masks. No smoothing, mesh decimation, or subdivision was applied to these exposure surfaces. Motor-area (MOp plus MOs) and caudoputamen surfaces were extracted from existing, approximately registered 80-µm anatomical labels. For anatomical context, the brain-plus-effective-CSF mask underwent two-iteration morphological closing and internal-hole filling before sampling at 160 µm. Only anatomical context surfaces were smoothed, simplified, and subdivided for display.

Anterior and lateral oblique views show the two independently calculated exposure regions at the same orthographic magnification, with 2-mm scale bars.

Representative spatial maps were presented alongside quantitative summaries from all ten skulls, a presentation approach also used for human transcranial ultrasound simulations [16]. The selection rule was specified for the present study before ranking the skulls: for each skull, the bilateral-mean water-peak intensity ratio and MOp-plus-MOs composition were compared with their respective ten-skull medians; absolute deviations were divided by each outcome's interquartile range and summed. The lowest-scoring skull (C3H_10M) was used for all Figure 2B–E maps. This rule identifies an example near the centers of the two quantitative outcomes, not a spatially median field. All ten individual skull maps are shown in Supplementary Figure S2, and Figure 2F shows paired peak-intensity ratios with side-specific means and skull-bootstrap 95% confidence intervals, alongside the predicted brain temperature-rise trajectory. The retained MOp-plus-MOs composition is shown in Supplementary Figure S2.

##### S3.6. Acoustic-model verification and limitations

Analytical reference solutions were used to check the propagation method. The measurement-derived source was evaluated against the water scans; numerical consistency was examined in the skull simulations. At the measured XY locations, the modeled peak was 0.961 W/cm$^2$ versus 1.194 W/cm$^2$ measured. The modeled axial profiles peaked 0.8 mm from the outlet at the sampled depths, versus 1.4 mm measured in both planes.

These differences limit depth localization and absolute exposure estimates. Convergence was not established across the full skull-model cohort. An independent tissue-level balance of acoustic power entering, leaving, and being absorbed in the modeled domain was not established. In the reference scenario, the absolute surface energy-balance residual relative to modeled absorption within the evaluation box was 29.740 ± 1.884% across ten skull models (mean ± SD), whereas the discrete balance residual relative to source work was 12.41 ± 5.16 ppm. These diagnostics use different definitions and denominators; the discrete residual does not independently verify the tissue-level physical balance. Neither residual represents a temperature-prediction error. Further limitations included treating attenuation as absorption, omission of skull shear-wave propagation, approximate tissue interfaces and atlas registration, and uncertain scalp coupling. The fields and exposure distributions were interpreted as conditional model predictions.

#### S4. Acoustic–thermal modeling

##### S4.1. Acoustic heat-source calculation

Volumetric heat sources were calculated from the simulated acoustic fields using the viscous dissipation term [17] for the carrier-matched Stokes attenuation model. For each tissue compartment, the time-averaged heat deposition rate was defined as $Q(r) = D\eta(r)|\nabla \cdot \hat{u}(r)|^2/2$, where $\hat{u}$ is the peak-amplitude particle-velocity phasor, $D = 0.5$ is the duty cycle, and $\eta = 2\rho c^3 \alpha / \omega^2$, with $\rho$, $c$, $\alpha$, and $\omega$ denoting tissue density, sound speed, attenuation coefficient, and angular carrier frequency, respectively. The duty cycle was incorporated once in the heat-source calculation. No additional $2\alpha I$ heating term was added for skull tissue, and energy dissipated within perfectly matched layers was excluded from tissue heating. The resulting tissue-specific heat sources were transferred to the 80-µm thermal grid using a conservative source-mapping procedure that preserved the integrated heat deposition within the modeled compartments. Acoustic fields and corresponding heat sources were calculated under the reference effective-scalp absorption assumption.

**Definitions**

$$Q(r) = \frac{D}{2}\eta(r)|\nabla \cdot \hat{u}(r)|^2$$

$$\eta(r) = \frac{2\rho(r)c(r)^3\alpha(r)}{\omega^2}, D = 0.5$$

Here, $Q$ has units of W/m³. The units of u, ∇·u, η, ρ, c, α, and ω are m/s, $s^{-1}$, Pa·s, kg/m³, m/s, Np/m, and rad/s, respectively.

##### S4.2. Bioheat simulation and stimulation sequence

Tissue temperature changes were simulated using a heterogeneous, incremental Pennes bioheat model on an 80-µm spatial grid [18]. Acoustic heat sources were applied according to six alternating 300-s left- and right-sided stimulation blocks, separated by five 10-s pauses, followed by 600 s of cooling. Each switching pause was fixed at 10 s to approximate experimental repositioning. Both left-first and right-first sequences were simulated for each skull under the reference material assumptions.

Temperature rise (ΔT, °C) was calculated relative to a matched no-sonication simulation with identical boundary conditions, with an initial ΔT of zero. Tissue-dependent specific heat capacity (J $kg^{-1}$ $K^{-1}$), thermal conductivity (W $m^{-1}$ $K^{-1}$), and perfusion coefficient ($s^{-1}$) were assigned as summarized in Supplementary Table S1, Part B. An assumed Robin heat-transfer coefficient of 6 W $m^{-2}$ $K^{-1}$ was applied at exterior and otherwise unspecified interfaces, while truncated boundaries were fixed at ΔT = 0. The model did not explicitly account for measured gel-mediated cooling, contact-pressure effects, or transducer self-heating. Time integration used a finite-volume Crank–Nicolson scheme [19] with 1-s steps during the first 30 s and 10-s steps thereafter, aligned to stimulation and cooling events. Two half-sized backward-Euler steps were used after heat-source changes to damp numerical transients [20].

S4.3. Temperature metrics and limitations

Maximum temperature rise was evaluated separately within the modeled brain, the bilateral primary motor area (MOp), the bilateral secondary motor area (MOs), skull, and effective dorsal scalp. For each skull and stimulation starting order, the maximum ΔT across all voxels in each region and all simulated time points was determined. The two starting-order maxima were then averaged within each skull, and results were summarized as mean ± sample standard deviation across ten independent skull models. For time-course visualization, the spatial maximum within each region was determined at each simulated time point and averaged across starting orders and skull models. The curves therefore represented spatial-maximum envelopes rather than temperature trajectories at fixed locations.

The reference effective-scalp absorption coefficient was 22.80832 Np/m at 1.078 MHz. Tissue properties and their sources are given in Supplementary Table S1, Part B. The predicted temperature rises depend on the assumed thermal contact conditions and the acoustic-model limitations described in S3.6.

S5. Wheel activity and body-state measurements

**S5.1. Wheel acquisition and endpoint definitions**

Wheel sensors registered one count per revolution (wheel diameter 140 mm, width 60 mm). Both intervention cohorts used the CIF-4A (Melquest, Toyama, Japan). In all 11 mice in the independent EEG/EMG cohort, a TUSB-S03CN3 counter logger (Turtle Industry, Tsuchiura, Japan) recorded wheel counts throughout Days 1–17, including during EEG/EMG recording. Counts were transferred to a computer via USB, saved in the same 1-min format, and analyzed in Python. Wheel analyses for the 32 behavioral-test mice covered the two dark phases preceding Day 17 testing (S8.3).

Wheel activity was classified into two states using the number of revolutions recorded in each 1-min bin. A bin containing one or more revolutions was classified as running, whereas a bin containing zero revolutions was classified as inactivity. Periods without wheel recording should not be interpreted as observed inactivity. State occupancy was defined as the proportion of valid 1-min bins classified as running or inactivity within the analysis period. Consecutive bins assigned to the same state were treated as a single bout, and bout duration was calculated in minutes. Transitions between states were evaluated using consecutive 1-min bins. The inactivity-to-running transition probability was calculated as the number of inactivity-to-running transitions divided by the total number of valid transitions originating from inactivity. The running-to-inactivity transition probability was calculated analogously using all valid transitions originating from running. Each denominator included all valid pairs of adjacent 1-min bins beginning in the specified state, including pairs in which that state persisted. Thus, the transition probabilities represented the likelihood of switching states in the next minute, conditional on the current wheel-defined state. The distinction between pauses and running intensity follows wheel-running microstructure studies in mice [21]. Conditional transitions between movement states have also been used in flies [22]. Here, both states were defined by wheel counts.

**S5.2. Daily interval totals and timestamp-aligned wheel profiles**

Figure 4A sums recorded rotations within each 24-h interval (ZT0–ZT24). Zeros entered during gaps in recording were not interpreted as observed inactivity.

For Figure 4B, minute timestamps were assigned to days using the 07:00 boundary and aggregated into complete 60-min hourly bins on Days 15–17. ZT1.5–5 was excluded from analysis. Requiring a complete hour therefore omitted the four hourly bins from ZT1 to ZT5. Missing values were neither filled with zeros nor interpolated. Each retained hourly value was averaged across the three days within each mouse before group means and SEM were calculated. These profiles were descriptive; no time-specific or longitudinal inferential tests were performed. The dark-phase endpoints in Figure 5B–F used Days 15–17, ZT12–24, with the definitions and statistical analyses described in the main Methods. Hourly activity was expressed in rotations/h and running intensity in rotations per active minute.

##### S5.3. Inactivity-bout persistence

Inactivity bouts comprised consecutive observed zero-rotation 1-min bins during the dark phases (ZT12–24) of Days 15–17. Runs occupying the first dark-phase bin were excluded because their onset was not observed; known-start bouts reaching dark-phase end were right-censored. Daily discrete Kaplan–Meier-type curves [23] estimated $S(k) = \Pr(T > k)$, where T was bout duration in minutes. At each integer minute k, bouts with observed duration $\geq k$ formed the risk set, and observed exits at k were events. The estimate was updated before plotting the step from k onward; censoring at k followed the event update. After the risk set was exhausted, zero estimates remained zero, whereas positive estimates became undefined.

Daily estimates were averaged equally across the three days within each mouse, followed by group means and SEM across nine equally weighted mice. A mouse-level estimate required all three daily estimates to be defined; a group summary required all nine mouse-level estimates to be defined. Figure 5G displays this descriptive estimator as percentages on a logarithmic 1–240-min axis, with curves restricted to their defined ranges. These descriptive curves were not subjected to time-specific or curve-level group tests. The P values in Figure 5F apply only to the separate median-duration endpoint, which retained window-edge bouts. The 50% crossing of the averaged persistence curve is not the group mean of the individual-mouse median values in Figure 5F.

##### S5.4. Body-weight and food-intake records

Body-weight and food-intake records were summarized for the wheel-activity intervention cohort ($n = 9$ per group). For each mouse, body-weight change was expressed as a percentage of its Day 10 body weight and summarized for Days 10, 13, 14, 15, 16, and 17. Mean daily food intake was derived from mouse-assigned consumption values for each measurement interval and expressed as a percentage change relative to the same mouse's mean daily intake over Days 0–10. The intervals were Days 0–10, 10–13, 13–14, 14–15, 15–16, and 16–17. Individual percentage changes were summarized as group means and SEM. Intervals of different durations were summarized separately. Body weight was recorded in g and mean daily intake in g per mouse per day before calculation of percentage changes.

#### S6. Novel-object-recognition test

##### S6.1. Behavioral apparatus and schedule

The novel-object-recognition test (NORT) was conducted in the behavioral-testing intervention cohort using a cylindrical arena (30.4 cm in diameter and 31 cm in height) under low-light conditions (approximately 0.03 lux). Mice underwent 10-min habituation sessions in the arena without objects on three consecutive days (Days 13–15). The object pairings for training and retention were adapted from Lueptow [24]. On Day 16 at ZT8, mice completed a 10-min training session with two identical cylindrical objects, one on each side of the arena. At ZT8 on Day 17, 24 h after training and after the daily tUS or sham procedure, mice underwent a 10-min retention test with one familiar cylinder and a novel triangular object. Mice were then returned to their cages, and OFT began approximately 2 h after NORT ended. NORT sessions were recorded using an infrared-illuminated USB camera (ELP-USBFHD05MT-DL36-J; ELP; OV2710 CMOS sensor) and OBS Studio at approximately 30 frames/s. Previously generated DeepLabCut keypoint coordinates were used for the subsequent rule-based quantification of object exploration.

##### S6.2. Retained coordinates, session geometry, and software provenance

Previously generated DeepLabCut coordinates [25,26] were used without rerunning pose estimation. Scorer identifiers in the outputs indicated SuperAnimal TopViewMouse with HRNet-W32 [27]. Detector identifiers included Faster R-CNN MobileNetV3 Large FPN and Faster R-CNN ResNet50 FPN v2. DeepLabCut v3.0.0rc6 was used. The output identifiers did not establish video-adaptation training settings.

Object exploration was scored using NORTAnalyzer versions 3.4.1 and 3.5.0. Analysis used each video's recorded frame rate, approximately 30 frames/s. Sentinel coordinates of −1 were treated as missing. When the tracking software returned multiple predictions for a mouse, one was selected using the nose and head-reference landmarks. The selection score was twice mean x-coordinate availability plus three times mean median likelihood for these two landmarks.

The familiar object remained on the left and the novel object on the right throughout retention testing, without counterbalancing. Manually defined image ROIs represented the familiar object as a circle and the novel object as a four-vertex polygon. These image ROIs were geometric analysis boundaries, distinct from the physical shapes of the objects. Pixel distances were scaled using the annotated arena radius divided by 15.2 cm. The left exploration ROI increased the object's radius by 2.5 cm. The right exploration ROI offset each polygon edge outward by 2.5 cm. The spatial calibration factor was expressed in pixels/cm.

NORT and OFT scoring was blinded to group identity. Group allocation, stimulation delivery, and wheel-data processing were unblinded.

##### S6.3. Pose filtering and tracking-speed flags

When both ear coordinates were finite, their midpoint defined the head reference. If only one ear coordinate was finite, it served as the reference. If neither was available, the tracked head-midpoint coordinate was used. When both ears contributed, their minimum likelihood represented head-reference confidence. Nose and head-reference coordinates were filtered independently in physical units using position–velocity Kalman filters [28].

The filter interval was 1/fps seconds, where fps is the frame rate in frames/s. Measurement standard deviation was 0.25 cm, and process acceleration standard deviation was 180 $cm/s^2$. Initial position and velocity standard deviations were 5 cm and 50 cm/s. Measurement variance was $0.25^2/\max(\text{likelihood}, 0.001)^2$ $cm^2$. Finite measurements with likelihood ≥0.70 updated the filters; other frames used prediction without measurement updates. Consequently, the subsequent geometric rules did not require raw nose likelihood ≥0.70 at every counted frame.

Nose displacement between consecutive filtered coordinates was converted to speed (cm/s) and smoothed using a five-frame uniform convolution. Frames exceeding median(speed) + 6 × [median absolute deviation(speed) + $10^{-9}$ cm/s] were flagged. Flags were extended by one frame on each side. The median absolute deviation was unscaled, and no additional fixed speed cap was applied. Flagged frames could not contribute exploration time. They were not subtracted from the observation clock used to accumulate the 300-s analysis window.

##### S6.4. Side-specific exploration rules and exclusion order

Let P denote the filtered nose position and H the filtered head reference. The left base rule required P within the expanded circular ROI and an angle ≤45° between P − H and the object-center vector from H. The right base rule used the angle between P − H and the vector from H to the nearest original polygon-boundary point. The right rule accepted an angle ≤45° within the expanded polygon, or P within the original polygon regardless of angle.

The occlusion rule accepted a frame if raw nose likelihood was below 0.20 or nonfinite and the predicted nose was inside the exploration ROI. A valid head-reference measurement with a likelihood of at least 0.70 was also required, and frames with a speed flag were excluded. It did not require the angular condition. No upper limit was imposed on the number of consecutive frames accepted under the occlusion rule.

An ear-confirmation rule accepted frames when both raw ear coordinates, each with likelihood ≥0.70, were inside the expanded ROI contracted inward by 1 cm. This rule did not require the angular condition. The familiar-object on-object mask required at least three of twelve torso landmarks within the original left circle, each finite with likelihood ≥0.40. The landmarks were left/right hip, midside and shoulder, mid_back, mid_backend, mid_backend2, mid_backend3, mouse_center, and neck.

Both base rules excluded speed-flagged and left on-object frames. Left-side occlusion rescue and ear confirmation also excluded left on-object frames. The corresponding right-side rescue rules excluded speed flags but did not explicitly exclude the left on-object mask. No right-side on-object detector was used. Quality-control records for the primary window showed no overlap between right exploration and left on-object classifications.

Frames satisfying both exploration masks were assigned exclusively by comparing squared distances from P to the left center and nearest right polygon boundary. Ties were assigned to the left object. Consecutive accepted frames formed events, including single-frame events, without a minimum event-duration requirement or gap filling. These asymmetric rules and fixed object positions prevented separation of object novelty from side-dependent classification effects.

##### S6.5. Observation window, inclusion, and discrimination index

For all 32 mice, object-exploration time was the summed duration of accepted object-directed approach behavior. Within the maximum 600-s retention interval, we accumulated 300 s of observation after excluding intervals classified as time on the familiar object. The window could span more than 300 s of elapsed time and was not restricted to frames with high tracking likelihood. Tracking-speed QC flags excluded frames from exploration but did not remove them from the cumulative observation clock.

Inclusion required completion of the 300-s observation window and total familiar-plus-novel exploration of at least 20 s. The included cohort comprised 32 mice: Ctrl/Sham and Ctrl/tUS, n = 7 each; SD/Sham and SD/tUS, n = 9 each. The discrimination index was (T_novel − T_familiar)/(T_novel + T_familiar), using exploration times within each mouse's selected window. Both exploration times were measured in seconds.

All 32 mice met the inclusion criteria. Agreement with independent manual exploration scoring was not evaluated in this cohort. The zero reference line for DI does not indicate a one-sample test.

#### S7. Open-field test

##### **S7.1. Behavioral apparatus and schedule**

On Day 17, each mouse in the behavioral-testing cohort explored a 50 × 50 cm open field for approximately 20 min at approximately 0.03 lux. We assessed locomotor activity by total distance traveled (m) and center exploration by time (s) in the central 30 × 30 cm square [29].

##### S7.2. Open-field coordinate processing and measurement quality control

For all 32 mice, OFT distance and center time were calculated from the mouse_center landmark tracked with DeepLabCut [25,26] using the SuperAnimal TopViewMouse model [27]. Coordinates with likelihood <0.90, sentinel values, or locations outside the video image were treated as missing. Only internal gaps bounded by valid coordinates and lasting no more than 1 s were linearly interpolated.

Remaining missing body-center coordinates could be reconstructed from mid_back, neck, and tail_base, each with likelihood ≥0.90. A within-session linear body-axis model predicted the center from tail-base-relative landmark vectors. Every fifth eligible frame was held out for validation, and at least 300 fitting frames were required. Reconstructions required validation RMSE within the session-specific limit, 20 pixels in the included records. Coordinates were then smoothed using centered five-frame moving medians followed by five-frame moving means. After smoothing, coordinates at unrecovered frames were again marked as missing.

Residual gaps could be filled using video-segmentation centroids after a session-specific validation against directly tracked coordinates. Segmentation used a 600 × 600-pixel rectified field and the pixelwise 90th percentile of up to 80 sampled frames as background. Positive background-minus-frame differences were Gaussian-blurred with a 5 × 5-pixel kernel and thresholded at 35 gray levels. Elliptical morphological opening and closing used 5 × 5- and 11 × 11-pixel kernels, respectively; the outer three image pixels were excluded. Candidate areas were 100–6,000 $px^2$ in the rectified image, with the largest candidate selected. Validation used up to 500 direct-tracking frames after the initial 10 s. Acceptance required a detection proportion ≥0.90 and RMSE ≤20 rectified pixels. Remaining missing frames after the initial 10 s were eligible for recovery. A candidate centroid required proximity within 120 rectified pixels to either the preceding accepted position or next valid position. Recovered runs were smoothed separately using five-frame medians followed by five-frame means.

Four floor corners defined a homography to the 50 × 50 cm arena. Frame-to-frame movements implying speeds >100 cm/s were removed and eligible short internal gaps were interpolated again. This process allowed up to ten iterations, followed by removal of any remaining above-threshold transitions. Total distance summed Euclidean displacements only between adjacent valid frames, with displacements <0.05 cm assigned zero. Unrecovered gaps were not bridged. Distances in centimeters were converted to meters for reporting.

The center was defined by 10 ≤ x ≤40 cm and 10 ≤ y ≤40 cm. Only sequences of consecutive frames classified as inside the center contributed to center time. The minimum sequence length was round(0.1 s × fps) frames, with a minimum of one frame. Center time was the number of retained center frames divided by fps. The retained full recording for each mouse, approximately 20 min, was used rather than the NORT 300-s window. Missing coordinates contributed neither distance increments nor center time. Absolute distance and center time were not extrapolated to replace missing observations.

Final valid-tracking fractions exceeded the 0.75 processing threshold. Validation against available tracked coordinates assessed recovery consistency, not accuracy against an independent manual ground truth. All 32 mice remained in the reported OFT analyses.

#### S8. Mouse-level summaries and statistical inference

##### S8.1. Experimental units, transformations, and model contrasts

The mouse was the experimental unit. The independent non-tUS EEG/EMG cohort comprised four Ctrl and seven SD mice (11 total). The wheel intervention cohort comprised nine mice per group (36 total). The same 32 mice completed NORT and OFT, with seven per Ctrl group and nine per SD group. For bilateral acoustic simulations, the skull was the unit of replication.

Wheel endpoints were calculated within each mouse-day during ZT12–24 and averaged equally across Days 15–17 within each mouse. Group summaries used equally weighted mouse values. The focal Days 15–17 interval was selected after inspection of the daily wheel-activity trajectories in Figure 4A and was subsequently used for the dark-phase analyses in Figure 5B–F. Dark-phase total rotations and median inactivity-bout duration used log(1 + x) transformations. Inactivity-to-running and running-to-inactivity probabilities used arcsin(√p). Running intensity, OFT distance, OFT center time, and NORT DI used identity scales.

Median inactivity-bout duration was calculated separately for each mouse and dark phase (ZT12–24). Consecutive observed zero-rotation 1-min bins formed inactivity runs. Runs touching ZT12 or ZT24 were retained at their observed within-window lengths, without excluding unknown-onset runs or applying censoring adjustments. Missing observations split runs, and the observed portions were retained separately without bridging gaps. The three daily medians for Days 15–17 were averaged equally within each mouse before calculating group means and SEM. All included mice had complete dark-phase records and finite daily medians on all three days.

For each intervention endpoint, a two-factor ordinary least-squares model included SD condition (Ctrl or SD), tUS condition (Sham or tUS), and their interaction (SD condition × tUS):

$$Y = \beta_0 + \beta_H H + \beta_T T + \beta_{HT} HT + \varepsilon.$$

H represented SD condition, coded 1 for SD and 0 for Ctrl; T represented tUS condition, coded 1 for tUS and 0 for Sham. Heteroscedasticity-consistent HC3 covariance estimates [30] were used for contrasts, two-sided t tests, and nominal 95% confidence intervals. Residual degrees of freedom were 32 for wheel outcomes and 28 for OFT/NORT outcomes.

For the five wheel and three behavioral outcomes, model-scale residual distributions were assessed using Shapiro–Wilk [31] and Jarque–Bera [32] tests, and heteroscedasticity using the Koenker version of the Breusch–Pagan test [33]. Internally studentized residuals, leverage, and Cook's distance [34] were examined for influential observations. No observations were excluded on the basis of these diagnostics.

Within-Ctrl and within-SD tUS-minus-Sham contrasts were $\beta_T$ and $\beta_T + \beta_{HT}$, respectively. The SD condition × tUS interaction, $\beta_{HT}$, was the difference between these two contrasts. The SD/Sham-minus-Ctrl/Sham contrast in each intervention outcome was $\beta_H$ from the same model. A significant within-SD contrast alone did not establish an SD-specific tUS effect. For transformed outcomes, model estimates and confidence intervals remained on the transformed scale. Arithmetic group means on the original scale were reported descriptively. Confidence intervals on transformed scales were not presented as differences on the original scale.

The additional 24-h Sleep-occupancy comparison used a two-sided unpaired Student t-test with equal variances and mouse-level percentages (SciPy 1.15.3; Python 3.11.2). The mean difference, 95% confidence interval, and P value were reported for this comparison.

##### S8.2. Multiplicity, missing observations, and descriptive displays

Tests of group differences and SD condition × tUS interactions were two-sided, with a significance threshold of $\alpha = 0.05$. For comparisons subject to Holm adjustment [35], significance was assessed using the adjusted P value.

Separate within-group comparisons of familiar- and novel-object exploration times in Supplementary Figure S15 used two-sided paired t-tests on mouse-level values (Novel minus Familiar; $\alpha = 0.05$). One comparison was performed in each of the four behavioral groups, with no adjustment across these four tests.

For each wheel outcome, Holm adjustment covered three two-sided group contrasts from the same HC3 model: tUS minus Sham within Ctrl, tUS minus Sham within SD, and SD/Sham minus Ctrl/Sham. For each OFT/NORT outcome, the family contained those three contrasts plus SD/tUS minus Ctrl/Sham, giving four comparisons. The additional contrasts and their adjustment families were defined during manuscript revision. All specified group contrasts entered their family regardless of nominal significance. The fourth behavioral contrast compared groups that differed in both SD condition and tUS condition. It did not test equivalence or normalization. For intervention endpoints, families were defined separately for each outcome; outcomes and cohorts were not pooled. The housing × tUS interaction was tested separately for each outcome, without correction across outcomes. Confidence intervals were pointwise nominal 95% intervals, not simultaneous intervals; exclusion of zero by such an interval did not guarantee significance after Holm adjustment. Holm adjustment across Wake, NREM, and REM remained confined to the three dark-minus-light occupancy comparisons in Table S2, using Welch tests as specified in S9.2. The three vigilance-state contrasts jointly assessed light–dark redistribution of the same sleep–wake composition and were therefore treated as one family. The 24-h sleep-architecture contrasts in Table S2B used equal-variance Student tests. Waking band-power contrasts are described in S9.3.

The intervention-model inputs contained finite observations for every included mouse and endpoint. No outcome imputation or additional animal exclusion was introduced for these models. Missing wheel observations were not recoded as inactivity, and transitions or bouts did not cross missing-data or day boundaries. Daily state metrics were averaged across available finite day-level values. All three days were present for every included mouse in the focal source table. Procedure-related unavailable observations in 24-h profiles were kept distinct from observed zero activity. Timestamp coverage alone did not establish observation validity during handling.

Legends identify individual values and summaries. Tables S3 and S4 (Part A) give group means. SEM is the standard error across mice. Figure S13 shows daily within-SD contrasts from separate four-group HC3 models with pointwise 95% CIs (df = 32). We made no time-specific or overall tests of the S5.3 persistence curves and no additional tests of hourly EEG profiles, body weight, food intake, or illustrative spatial displays.

We fitted the intervention models, estimated HC3 contrasts, and applied within-outcome Holm adjustment in Python using NumPy, pandas, SciPy, and statsmodels.

##### S8.3. Wheel activity before behavioral testing

We selected three measures of behavioral persistence and switching based on the tUS wheel-activity experiment: activity-resumption probability, median inactivity-bout duration, and activity–inactivity switches per hour. In the separate behavioral-test cohort, each measure was calculated for each of the last two complete dark phases before testing (Days 15 and 16, ZT12–24). A minute with at least one rotation was active. Resumption probability was the fraction of adjacent minute pairs starting in inactivity that ended in activity. Inactivity bouts comprised consecutive inactive minutes. Switches in either direction were summed and divided by 12 h (switches/h). We averaged the two nightly values equally within each mouse and matched them by animal ID to Day 17 NORT DI and OFT center time. In subsequent comparisons, we paired two measures of running amount and intensity with NORT DI, OFT center time, and total OFT distance. For each dark phase, total rotations were the sum of the minute counts, and running intensity was total rotations divided by the number of active minutes. The two nightly values were averaged equally within each mouse. For each of the 12 pairs, we calculated Spearman's ρ [36] across all 32 mice pooled across groups and a two-sided P value from 99,999 unrestricted permutations of the outcome across mice [37]. Each pair was tested separately.

#### S9. Independent EEG/EMG experiment

##### S9.1. Electrode implantation, placement and recording

EEG/EMG electrodes were implanted under inhaled isoflurane anesthesia in the independent EEG/EMG cohort using a 2-EEG/1-EMG headmount (8201; Pinnacle Technology, Lawrence, KS, USA). Electrode positions were defined relative to bregma. The right-sided EEG2 and EEG1 screw electrodes were placed at anteroposterior (AP) +1.8 mm, mediolateral (ML) +1.1 mm and AP −4.0 mm, ML +1.1 mm, respectively. The left-sided ground (AGND) and common reference (EEG COMMON) electrodes were positioned at AP +1.8 mm, ML −1.1 mm and AP −4.0 mm, ML −1.1 mm, respectively. The electrodes were secured using a self-curing acrylic dental resin (UNIFAST III; GC Corporation, Japan). Two EMG leads were implanted in the nuchal muscles, one on each side. Following electrode implantation, mice were allowed a 1-week postoperative recovery period before the start of the experimental timeline (Day 0). During recovery, mice were habituated to tethering with a dummy cable. This was intended to familiarize them with cable resistance and discourage cable chewing during subsequent EEG/EMG recordings. The combination of screw EEG electrodes and nuchal EMG leads follows established mouse sleep-recording methods [38].

On Day 16 at ZT8 (15:00), mice were connected to the recording cable while remaining in their respective wheel-housing conditions, and continuous EEG/EMG recording was maintained until Day 18 at ZT3 (10:00). During recording, the implanted headmount was connected to a head-mounted preamplifier configured for sleep recording (8202-SL; 2 EEG/1 EMG; Pinnacle Technology). Amplified signals were transmitted through a commutator to a data-conditioning and acquisition system (8206-HR; Pinnacle Technology) and acquired at 400 Hz using SIRENIA ACQUISITION (Pinnacle Technology). The acquisition-stage EEG low-pass filter was set to 40 Hz. The physiological characterization used the continuous 24-h interval from Day 17 at ZT0 (07:00) to Day 18 at ZT0 (07:00). Raw EMG was acquired at 400 Hz and is displayed in μV using the EDF physical calibration. For display only, the minimum and maximum raw samples in each 5-s overview interval or 0.5-s expanded interval were retained at their original timestamps. No additional filtering, rectification, normalization or RMS transformation was applied.

S9.2. EEG/EMG vigilance-state analysis

We calculated overall and hourly vigilance-state occupancy in Ctrl and SD mice. EEG/EMG-derived vigilance-state labels were obtained using FASTER2 (version 0.5.5) [39] and retained without manual correction. Each 8-s epoch was classified as Wake, non-rapid eye movement (NREM) sleep, or rapid eye movement (REM) sleep. Analyses were restricted to the continuous 24-h recording interval from Day 17 at ZT0 to Day 18 at ZT0. The light and dark phases were defined as ZT0–ZT12 and ZT12–ZT24, respectively. For each mouse, vigilance-state occupancy was calculated as the proportion of valid EEG/EMG recording time assigned to each state. Hourly occupancy was calculated by dividing the total duration of valid epochs assigned to each state by the total duration of all valid epochs within the corresponding mouse-hour. Group-level hourly profiles were summarized as the mean ± standard error of the mean (SEM) across mice. Hourly profiles were presented descriptively, without tests at individual time points. Vigilance-state classification was performed without knowledge of Ctrl/SD group assignments.

To characterize the temporal stability of vigilance states beyond their overall occupancy [40], bout duration was evaluated for Wake, NREM sleep, and REM sleep. Direction-specific transition rates and probabilities were also calculated to assess the frequency and direction of switching between vigilance states. Consecutive valid 8-s epochs assigned to the same vigilance state were grouped into bouts, and the median bout duration (s) was calculated for each state and mouse. Missing or unknown epochs were not bridged when identifying bouts or state transitions. Direction-specific transitions among Wake, NREM, and REM sleep were counted using consecutive valid epochs. For a transition from state $i$ to a different state $j$, the transition rate ($R_{i\to j}$) and conditional transition probability ($P_{i\to j}$) were defined as

$$R_{i\to j}=\frac{N_{i\to j}}{T_{\text{valid}}},\; P_{i\to j}=\frac{N_{i\to j}}{\sum_{k\in\{W,N,R\}} N_{i\to k}},$$

where $N_{i\to j}$ denotes the number of valid adjacent epoch pairs with the first epoch in state $i$ and the second in state $j$, and $T_{\text{valid}}$ is the valid recording duration in hours. The denominator of $P_{i\to j}$ included same-state persistence ($i\to i$). Here, W, N, and R denote Wake, NREM sleep, and REM sleep, respectively, and Sleep was defined as NREM and REM combined. For each 24-h outcome listed in Supplementary Table S2, Part B, mouse-level values were compared between Ctrl and SD mice using a two-sided unpaired Student's t-test assuming equal variances. Transition rates are expressed as transitions per valid recording hour.

Combined Sleep bouts joined contiguous NREM and REM epochs across sleep-stage transitions; Wake bouts comprised contiguous Wake epochs. Individual-mouse median durations were calculated separately for ZT0–12 and ZT12–24, retaining observed portions of phase-edge bouts. Bidirectional Sleep↔Wake counts included valid adjacent epoch pairs within each phase and were divided by valid recording hours; the pair crossing ZT12 was excluded from both phase counts. Each mouse contributed one value per phase. These phase-specific analyses were added after examination of the 24-h summaries. Ctrl–SD contrasts used two-sided unpaired equal-variance Student t tests, reporting P values and pointwise 95% CIs for the biologically distinct light and dark phases. The same mice contributed to both phases; connecting lines identify individual mice. No phase-specific treatment effect is inferred from differences in significance between phases.

We evaluated the correspondence between wheel status and EEG/EMG-defined vigilance in both directions, including the sensitivity and specificity of wheel status for vigilance and their balanced accuracy [41].

Each 8-s vigilance epoch was apportioned to 1-min wheel bins according to temporal overlap in seconds. An epoch spanning a minute boundary contributed 4 s to each adjacent bin. Minutes with at least one rotation were classified as active, and minutes with zero rotations as inactive. The timing of individual rotations within that minute was unavailable.

Time without a Wake, NREM or REM label and bins without matched wheel observations were excluded. Scored epochs were retained irrespective of the FASTER2 QC flags. For each mouse, conditional probabilities were calculated as the valid overlapping duration satisfying both conditions divided by the valid overlapping duration of the conditioning state. Group means and SEM were calculated across the mouse-level probabilities, with each mouse contributing equally.

For Wake and Sleep (NREM and REM combined), we defined sensitivity, specificity, and balanced accuracy as follows.

$$\text{Sensitivity} = P\left(\text{Active} \mid \text{Wake}\right)$$

$$\text{Specificity} = P\left(\text{Inactive} \mid \text{Sleep}\right)$$

$$\text{Balanced accuracy} = \frac{\text{Sensitivity} + \text{Specificity}}{2}$$

We also calculated $P\left(\text{Wake} \mid \text{Active}\right)$ and $P\left(\text{Sleep} \mid \text{Inactive}\right)$, conditioning on wheel activity status. Wheel inactivity was treated as a behavioral measure rather than as a direct measure of sleep.

Phase-dependent changes were evaluated to distinguish light–dark redistribution from differences in overall 24-h occupancy of Wake, NREM sleep, and REM sleep under SD conditions. For each mouse, the occupancy of each state was calculated separately for the light (ZT0–ZT12) and dark (ZT12–ZT24) phases. The phase-related change in state occupancy was defined as

$$\Delta O_s = O_{s,\text{Dark}} - O_{s,\text{Light}},$$

where $O_{s,\text{Dark}}$ and $O_{s,\text{Light}}$ represent the occupancy percentages of vigilance state $s$ during the respective phases. Their difference is expressed in percentage points.

For each vigilance state, the untransformed mouse-level dark-minus-light difference in occupancy was compared between Ctrl and SD mice using a two-sided Welch's t-test. Between-group differences were expressed as SD minus Ctrl in percentage points, with nominal 95% confidence intervals calculated using Welch degrees of freedom. Both nominal and Holm-adjusted P values were reported, with adjustment across the three comparisons for Wake, NREM, and REM. The three vigilance-state contrasts jointly assessed light–dark redistribution of the same sleep–wake composition and were therefore treated as one family.

##### S9.3. Absolute EEG power during wakefulness

We analyzed calibrated EEG2 recordings (400 Hz) from Day 17 ZT0 to Day 18 ZT0 in the independent non-tUS cohort. We divided each 8-s FASTER2 Wake epoch with zero EEG missing-sample and outlier fractions into two nonoverlapping 4-s windows. Windows within the calibrated digital range were converted to µV. After mean removal, we estimated a Hann-windowed periodogram [42] and summed spectral density ($\mu V^2$/Hz) multiplied by the 0.25-Hz bin width over delta (0.5–<4 Hz), theta (6–<9 Hz [43]), sigma (10–<15 Hz), and beta (15–<30 Hz).

We averaged absolute power ($\mu V^2$) over valid windows within each mouse. All-wake estimates included windows without a matched wheel count. For the wheel-conditioned analysis, each 4-s window was assigned the wheel status of its enclosing minute (S9.2). Prior comparisons of waking EEG spectra during voluntary wheel running and novel-object exploration informed this comparison [44].

We tested differences in mean log power between housing conditions by enumerating all 330 assignments of four Ctrl and seven SD mice. For the within-mouse active-to-inactive contrast, we tested the mean log ratio across all 11 mice using all 2,048 sign assignments [45]. Between-group differences in log ratios used the same housing-label permutation test. Tests were two-sided. Holm adjustment covered four bands separately for each of these three questions, and eight comparisons for the housing contrasts within active and inactive minutes. We report back-transformed group means and ratios (Table S2, Part C).

#### S10. Independent wheel and activity-rhythm characterization

The independent non-tUS cohort (Ctrl, n = 4; SD, n = 7) had 17 days of minute-level wheel counts. Daily totals and hourly profiles used these longitudinal records. For the simultaneous Day 17 profile, hourly wheel rates were rotations per valid wheel-recording minute multiplied by 60 (54–60 valid minutes per mouse-hour). Vigilance-state occupancy and group summaries followed S9.2.

Individual 30-min actograms were displayed as 48-h double plots using a per-mouse logarithmic scale referenced to the 95th percentile of positive 30-min counts (Supplementary Figure S10C). Double plotting followed the standard actogram convention for rodent wheel records [46].

The five dark-phase wheel endpoints used the same state definitions, daily estimates and arithmetic three-day within-mouse averaging as the intervention experiment (Days 15–17, ZT12–24). Median bouts used window-bounded consecutive inactive runs. Persistence curves used the onset and censoring rules in S5.3; curves were averaged across days within mice and then across mice. Each outcome had one two-sided, equal-variance Student t test (df = 9), using log(1+x) totals and median bouts, arcsine-square-root transition probabilities and original-scale intensity. P values were nominal, without across-outcome adjustment.

For activity-rhythm characterization, we used the last three complete light–dark cycles before SD onset on Day 10 at ZT8 (Days 7–9) and the last three before cable attachment on Day 16 at ZT8 (Days 13–15). The final 48 h (Days 16–17), beginning 8 h before cable attachment at Day 16 ZT8, were summarized descriptively. Dark-phase rotation share and dark-minus-light active-minute occupancy were calculated per mouse. Interdaily stability (IS) and intradaily variability (IV) were calculated from hourly counts. IS compared the variance of the mean 24-h hourly profile with total hourly variance. IV compared successive-hour squared differences with total variance. For each mouse and analysis window, minute-level activity was first averaged across days at each time of day. M10 and L5 were the highest 600-min and lowest 300-min moving averages, respectively, on this circular 24-h profile. Relative amplitude was RA = (M10 − L5)/(M10 + L5) [47,48]. Dark-phase rotation share is expressed as a percentage, the dark-minus-light occupancy difference in percentage points, and M10 and L5 in rotations/min.

A fixed-24-h single-harmonic model (intercept plus sine and cosine terms) was fitted to hourly counts [49]. Fitted amplitude, MESOR and acrophase were summarized descriptively. Circular statistics [50] were used to summarize phase and the start times of the M10 and L5 windows. These indices describe behavior under the imposed light–dark schedule; a free-running circadian period was not estimated. For each of six endpoints, including $R^2$, the within-mouse change from Days 7–9 to Days 13–15 was compared between groups using a two-sided equal-variance Student t test (df = 9), with pointwise 95% CIs and no adjustment of P values across endpoints. These additional analyses were conducted after the principal intervention analysis; their computational specifications were fixed before execution. Amplitude and MESOR are expressed in rotations/h, and acrophase and M10/L5 start times in Zeitgeber hours.

#### Supplementary Results

##### Regional temperature predictions under reference material assumptions

Under the reference material assumptions, the maximum predicted temperature rise (ΔT) within the modeled brain was 0.3759 ± 0.0105°C (mean ± SD, n = 10 skull models; Supplementary Fig. S3). The corresponding maxima within the bilateral MOp and MOs regions were 0.3000 ± 0.0085°C and 0.3161 ± 0.0090°C, respectively. Maximum predicted temperature rises in the skull and effective dorsal scalp were 0.4772 ± 0.0175°C and 0.5078 ± 0.0211°C, respectively.

##### Temporal profiles and spatial distribution of predicted heating

The temporal profiles in Figure 2F (right) represented spatial-maximum temperature envelopes averaged across stimulation starting orders and skull models, rather than temperature changes at fixed anatomical locations. The brain maximum lay outside the approximately registered MOp and MOs regions in 20 of the 20 skull-by-starting-order simulations. The predicted maxima occurred at 1,540 or 1,850 s.

#### EEG/EMG-defined sleep organization

In the independent non-tUS EEG/EMG experiment (Ctrl, n = 4; SD, n = 7), mice in the SD condition spent a greater proportion of the day asleep. Twenty-four-hour sleep occupancy was 61.23% in SD mice and 49.47% in Ctrl mice (difference [SD − Ctrl], 11.76 percentage points; 95% CI, 8.83–14.69; Student's t(9) = 9.080; P = 7.94 × $10^{-6}$; Figure 3A). The light–dark contrast in Wake occupancy was also smaller under SD: the dark-minus-light difference was 15.50 percentage points in SD versus 24.53 in Ctrl (between-group difference, −9.03 percentage points; nominal 95% CI, −13.84 to −4.22; Holm-adjusted P = 0.00693; Supplementary Table S2).

Phase-specific analysis showed lower mean individual-mouse Sleep-bout medians in SD during the light phase (303.4 versus 403.0 s; P = 0.0306), with a less clear dark-phase difference (280.0 versus 310.0 s; P = 0.602; Supplementary Figure S5A). Wake-bout medians were lower in both phases (18.3 versus 30.0 s in light, P = 0.00204; 25.1 versus 60.0 s in dark, P = 0.00158; Supplementary Figure S5B). Sleep↔Wake transitions were more frequent during both light (13.38 versus 8.88 per valid recording hour, P = 0.00299) and dark (10.29 versus 6.83, P = 0.0218; Supplementary Figure S5C). Contrasts and nominal 95% CIs are included in Table S2.

Greater sleep occupancy coexisted with shorter sleep-stage bouts. Mean individual-mouse median NREM-bout duration was 87.4 s in SD mice versus 290.0 s in Ctrl mice (P = 5.37 × $10^{-6}$; Table S2). REM bouts were also shorter: mean individual-mouse medians were 22.86 versus 77.00 s (difference [SD − Ctrl], −54.14 s; 95% CI, −63.82 to −44.47; Student's t(9) = −12.66; P = 4.87 × $10^{-7}$; Table S2). Individual 24-h state sequences illustrated closely interspersed NREM and REM periods in SD mice, with variation in their temporal arrangement across animals (Supplementary Figure S6).

Total vigilance-state transitions were more frequent in SD than in Ctrl mice (31.54 versus 10.54 transitions/h; P = 1.27 × $10^{-5}$; Table S2). Most of the between-group increase involved transitions between NREM and REM: their bidirectional frequency was 19.71 versus 2.68 transitions per valid recording hour (difference, 17.03; 95% CI, 12.39–21.67; P = 1.64 × $10^{-5}$; Table S2). Sleep↔Wake transitions were also more frequent (11.83 versus 7.86 per valid recording hour; difference, 3.97; 95% CI, 1.80–6.14; P = 0.00255; Table S2). A separate measure counted only Sleep→Wake transitions per hour spent in Sleep. This rate was 9.70 in SD and 7.96 in Ctrl, with no statistically significant difference (difference, 1.73; 95% CI, −0.35 to 3.82; P = 0.0919; Table S2).

Simultaneous wheel recordings, raw EMG traces, and classified vigilance states illustrated how wheel behavior related to physiological state over 24 h and within expanded 120-min intervals (Figure 3C). Across mice, Sleep occupied 86.34% of wheel-inactive time under SD and 59.34% under Ctrl (Figure 3D). Wake occupied 87.97% of the time within 1-min bins classified as wheel-active under SD and 99.85% under Ctrl. The individual-mouse decomposition of inactive periods showed lower mean Wake occupancy and higher mean NREM and REM occupancy in SD than in Ctrl mice (Supplementary Figure S7A). In both the light and dark phases, Sleep occupied a greater proportion of wheel-inactive time in SD mice than in Ctrl mice (Supplementary Figure S7B). Wheel inactivity thus captured periods enriched for Sleep, especially under SD conditions, while still including Wake. All of these physiological comparisons came from mice that received no tUS or sham procedures.

Across 24 h, the group means of the mouse-level median Sleep-bout durations were 286.9 s in SD mice and 372.0 s in Ctrl mice. The between-group comparison was not statistically significant (Student's t(9) = −2.160, P = 0.0591; 95% CI for SD − Ctrl, −174.3 to 4.0 s).

#### Absolute EEG power during wakefulness

Across wakefulness, delta power was higher in SD mice (Figure S8A). In wheel-active minutes, waking theta power exceeded that in wheel-inactive minutes in 10 of 11 mice (geometric mean within-mouse ratio, 1.24; Holm-adjusted P = 0.0039; Figure S8B). This association is consistent with greater waking theta activity during voluntary wheel running [43]. Delta, sigma, and beta power also increased in active minutes. The wheel-associated power ratios showed no significant housing difference after Holm adjustment. In wheel-inactive minutes, SD mice had higher delta power than Ctrl mice, and in wheel-active minutes they had lower beta power (Table S2, Part C).

#### Wheel activity across the experimental timeline

Supplementary Figures S9 and S10 show vigilance–wheel correspondence and longitudinal wheel activity in the independent EEG/EMG cohort (Ctrl, n = 4; SD, n = 7; no tUS or sham). For dark phases on Days 15–17 (ZT12–24), group means of within-mouse three-day averages (Ctrl versus SD) were 15,148.6 versus 5,884.1 daily rotations; 48.93 versus 17.99 rotations per active minute; P(I→R), 0.0940 versus 0.1358; P(R→I), 0.0950 versus 0.1780; and daily median inactivity-bout duration, 4.958 versus 2.690 min. Endpoint-specific tests and uncertainty estimates are in the Supplementary Source Data (nominal P values; no across-endpoint adjustment). These mice were not pooled or statistically compared with the intervention cohort. Individual actograms for all 11 mice appear in Supplementary Figure S10C.

#### Activity-rhythm organization before and during SD

Figure S11 shows hourly wheel-activity profiles, and Figure S12 shows each mouse's rhythm indices during the two 72-h windows. Within-mouse changes and their group contrasts are given in Supplementary Table S2, Part D. The between-group differences in within-mouse changes (SD − Ctrl) were −0.331 for interdaily stability (95% CI −0.517 to −0.145, P = 0.00297), +0.400 for intradaily variability (95% CI 0.0386 to 0.762, P = 0.0337), and −0.473 for fixed-24-h single-harmonic $R^2$ (95% CI −0.672 to −0.273, P = 0.000453). The corresponding P values for dark-phase rotation share, dark-minus-light active-minute occupancy and relative amplitude were 0.433, 0.509 and 0.0680.

Before SD onset, dark-phase rotations already accounted for approximately 87.1% of daily rotations in Ctrl mice and 65.0% in SD-assigned mice. Absolute rotations subsequently fell during both phases in SD mice, so a larger light-phase share could arise even as daytime activity declined. The differences in stability, variability and 24-h fit were evident in the pre-cable comparison. A change in the activity waveform can reduce the fit to the imposed 24-h light–dark cycle. The intrinsic circadian period was not measured. The terminal 48-h profiles are descriptive and include both pre- and post-attachment recording.

#### Physiological context

In the independent non-tUS cohort, greater sleep occupancy coexisted with shorter NREM bouts. Much of the increase in transitions involved NREM↔REM switching. Sleep↔Wake transitions also increased per hour of valid recording, while the Sleep→Wake rate per hour of sleep showed no clear group difference (Table S2). Other models illustrate the importance of transition direction. Chronic stress increased Wake→NREM transition counts without a significant change in NREM→REM counts [51]. Mice with Bmal1 selectively deleted in histaminergic neurons had more NREM→REM and REM→NREM transitions during the light phase [52].

### Supplementary Tables

Table S1. Stimulation and measured acoustic exposure parameters, and material properties and assumptions for acoustic–thermal modeling.

Part A gives the stimulation settings and water measurements. Part B gives the model inputs and their sources (Supplementary Methods S2–S4).

Part A — Stimulation and measured acoustic exposure

| **Parameter** | **Experimental specification** |
|---|---|
| Transducer | Custom-made planar piezoelectric transducer, 20-mm diameter, with matching box; Honda Electronics Co., Ltd. Neither geometric nor electronic focusing. |
| Collimator and mounting | 2-mm outlet; 10-mm tapered section; Clear Resin, printed with Form 3B (Formlabs); F44 slide lens holder (Suruga Seiki). |
| Driving equipment | WF1973 multifunction generator and HSA4101 high-speed bipolar amplifier (NF Corporation). |
| Stimulation carrier frequency | 1.078 MHz. |
| Pulse sequence | PRF 40 Hz; 25-ms pulse-repetition period; 12.5 ms ON and 12.5 ms OFF; nominal duty cycle 50%. |
| Nominal targets and positioning | Bilateral motor cortex; AP +1.5 mm, ML ±1.5 mm relative to bregma; stereotaxic positioning (RWD Model 68801). |
| Acoustic coupling | Pro Jelly, Normal Type (Jex), filling the collimator taper and applied to the scalp. |
| Stimulation blocks | Six alternating left/right 5-min blocks; three blocks per hemisphere. Total programmed block duration 30 min; 15 min per hemisphere. |
| Cumulative acoustic ON time | 15 min per daily session. This is distinct from the 30-min programmed block total. |
| Repeated sessions | Once daily on Days 13–17, between ZT2 and ZT4, under the assigned Ctrl or SD condition. |
| Anesthesia | Isoflurane: 5% induction; 1.5–2.5% maintenance. |
| Sham procedure | Same anesthetic conditions, matched anesthesia duration and daily schedule. Separate box; no stereotaxic fixation, transducer, scalp gel or ultrasound. Fixation, contact and coupling were not matched. |

Free-field acoustic measurement

| **Parameter** | **Measured value or definition** |
|---|---|
| Measurement system | Calibrated HNR-0500 needle hydrophone (Onda Corporation); measurements in water. |
| Measured water-field carrier frequency | Approximately 1.078 MHz. |
| Measurement pulse settings | PRF 40 Hz; nominal 12.5-ms ON and 12.5-ms OFF intervals. |

| Parameter | Measured value or definition |
|---|---|
| Pressure conversion | Baseline-corrected hydrophone voltages converted to acoustic pressure using the effective calibration sensitivity. |
| Temporal averaging interval | Complete pulse-repetition intervals, including ON and OFF periods (nominally 25 ms); not a 5-min block or whole-session average. |
| Free-field reference intensity | Maximum $I_{TA}$ within the sampled XY plane at Z = −1.4 mm: 1.193585 W/cm² (1.194 W/cm² in Main Results). |
| Measurement-map inclusion | All 3,147 retained cells in the final acoustic measurement maps; the peak-intensity ratio uses the sampled XY-plane maximum, not an exhaustively determined three-dimensional maximum. |

Notes for Part A

a. The stimulation carrier frequency (1.078 MHz) and measured water-field carrier frequency (approximately 1.078 MHz) are reported separately. Target coordinates describe intended placement and do not establish selective neural activation.

b. The 30-min total comprises programmed stimulation blocks and excludes repositioning intervals. The five fixed 10-s pauses in Part B are simulation assumptions, not measured event timestamps or a fixed experimental session duration.

c. Measured pulse-average and temporal-average intensities were retained separately; their ratio was not forced to equal the nominal duty cycle. No pulse-average intensity, peak negative pressure or mechanical index is inferred from the temporal-average reference in this table.

Part B — Material properties and modeling assumptions

| Tissue and source | ρ<br>kg m⁻³ | c<br>m s⁻¹ | α<br>Np m⁻¹ | $C_p$<br>J kg⁻¹ K⁻¹ | k<br>W m⁻¹ K⁻¹ | w<br>s⁻¹ |
|---|---|---|---|---|---|---|
| Brain [53] | 1030 | 1560 | 2.6259 | 3600 | 0.5 | 0.008 |
| Effective CSF [54] | 1000 | 1500 | 0.029052 | 4180 | 0.6 | 0 |
| Skull [53,55,56] | 1933 | 2425 | 36.123 | 1700 | 0.3 | 0.0003 |
| Effective dorsal scalp [54] | 1109 | 1595 | 22.808 | 3391 | 0.37 | 0.00197 |
| Other head tissue [54] | 1090 | 1580 | 7.6635 | 3421 | 0.49 | 0.000715 |

ρ, density; c, sound speed; α, acoustic attenuation coefficient at 1.078 MHz; $C_p$, specific heat capacity; k, thermal conductivity; w, perfusion coefficient. Values are effective model inputs, not measurements in the experimental mice.

| Model component | Adopted setting or assumption |
|---|---|
| Anatomical model cohort | Ten independent P90 male C3H/HeJ skull models; distinct from experimental C3H/HeNSlc mice. Common MRI-template tissue compartments and approximate atlas registration. |
| Acoustic solver and grid | Linear heterogeneous compressional-wave model; k-space pseudospectral method; 120-µm grid; 128 temporal samples per acoustic cycle; 1.078 MHz. |
| Source and duty cycle | Amplitude calibrated to the measured carrier amplitude along the water depth profile (S3.3); spatial source distribution reconstructed from measured XY intensity and drive-referenced carrier phase. Separate left/right fields; duty cycle 0.5 applied once. |
| Effective scalp absorption | Reference 22.80832 Np/m at 1.078 MHz. Skull coefficient fixed. Values are effective model inputs, not measurements in the experimental mice. |
| Bioheat model and grid | Heterogeneous incremental Pennes model; 80-µm thermal grid; conservative transfer of acoustic heat sources. |
| Simulated sequence | Six alternating 300-s blocks; five assumed 10-s switching pauses; 600-s cooling. Total 2,450 s. Both left-first and right-first starting orders. |
| Thermal reference and boundaries | Initial $\Delta T = 0$ relative to matched no-sonication conditions. Assumed Robin coefficient $h = 6$ W $m^{-2}$ $K^{-1}$ at exterior/otherwise unspecified interfaces; truncated boundaries fixed at $\Delta T = 0$. |
| Blood properties | Density 1,030 kg $m^{-3}$; specific heat capacity 3,617 J $kg^{-1}$ $K^{-1}$. |
| Time integration | Finite-volume Crank–Nicolson scheme: 1-s steps for the first 30 s, then 10-s steps aligned to events; two half-sized backward-Euler steps after heat-source changes. |

Notes for Part B

Attenuation was treated as absorption where tissue-specific absorption data were unavailable. The model omitted bone shear waves, contact-related cooling and transducer self-heating. Tissue-level acoustic power balance remains incompletely verified. Values are rounded here; the source data retain full precision.

Parameter provenance

Brain properties were transferred from Phipps et al. [53], including density, sound speed, heat capacity, thermal conductivity, and perfusion; brain attenuation was converted from the source frequency law at 1.078 MHz. Skull density (1933 kg/m³) and sound speed (2425 m/s) came from Liang et al. [55], while skull heat capacity and conductivity came from Phipps et al. [53]. The skull coefficient used the 1-MHz human-bone absorption value reported by Pinton et al. [56], with quadratic frequency extrapolation as a model assumption. Scalp and other-tissue density, sound speed, heat capacity, and conductivity used the skin and muscle rows of Wang et al. [54]; attenuation coefficients were converted at the carrier frequency. These are transferred surrogates, not mouse-specific measurements. The effective CSF compartment used water-like surrogate properties; its adopted sound speed (1500 m/s), heat capacity (4180 J/kg/K), and quadratic attenuation scaling were model choices rather than unchanged transfers of the water row in Wang et al. [54]. Nonbrain perfusion coefficients were modeling assumptions. Carrier-frequency coefficients and the solver's frequency-squared loss law are distinct from the frequency laws in the source studies.

**Supplementary Table S2. Sleep–wake measures, waking EEG power, and wheel-activity rhythms in the independent EEG/EMG cohort.**

Part A. Dark-minus-light vigilance-state occupancy

| Vigilance state | Ctrl (n = 4), pp | SD (n = 7), pp | SD − Ctrl, pp (95% CI) | Nominal P | Holm-adjusted P |
|---|---|---|---|---|---|
| Wake | +24.53 | +15.50 | −9.03 (−13.84 to −4.22)<br>t = −4.277<br>df = 8.560 | 0.00231 | 0.00693 |
| NREM | −19.89 | −12.67 | +7.23 (+1.56 to +12.89)<br>t = 2.885<br>df = 8.999 | 0.01805 | 0.03609 |
| REM | −4.64 | −2.84 | +1.80 (−2.29 to +5.90)<br>t = 1.073<br>df = 6.080 | 0.32389 | 0.32389 |

pp = percentage points; CI = confidence interval.

Values are group means of mouse-level dark-minus-light occupancy differences (percentage points). SD-minus-Ctrl contrasts use two-sided Welch tests and pointwise nominal 95% CIs. Holm adjustment covers the three states (Wake, NREM and REM).

Part B. Twenty-four-hour and phase-specific EEG/EMG comparisons

| **Outcome (unit)** | **SD − Ctrl** | **95% CI** | **t** | **df** | **P** |
|---|---|---|---|---|---|
| 24-h Sleep occupancy (percentage points) | +11.76 | 8.83 to 14.69 | 9.080 | 9 | $7.94 \times 10^{-6}$ |
| Median NREM-bout duration (s) | −202.57 | −250.69 to −154.45 | −9.523 | 9 | $5.37 \times 10^{-6}$ |
| Total vigilance-state transition frequency (transitions per valid recording hour) | +21.00 | 15.46 to 26.54 | 8.568 | 9 | $1.27 \times 10^{-5}$ |
| Median REM-bout duration (s) | −54.14 | −63.82 to −44.47 | −12.662 | 9 | $4.87 \times 10^{-7}$ |
| Bidirectional NREM↔REM transition frequency (transitions per valid recording hour) | +17.03 | 12.39 to 21.67 | 8.307 | 9 | $1.64 \times 10^{-5}$ |
| Bidirectional Sleep↔Wake transition frequency (transitions per valid recording hour) | +3.97 | 1.80 to 6.14 | 4.132 | 9 | 0.00255 |
| Sleep→Wake transition frequency (transitions per Sleep hour) | +1.73 | −0.35 to 3.82 | 1.886 | 9 | 0.0919 |
| Median combined Sleep-bout duration (s) | −85.14 | −174.33 to 4.04 | −2.160 | 9 | 0.0591 |
| Light: median Sleep-bout duration (s) | −99.57 | −187.51 to −11.64 | −2.561 | 9 | 0.0306 |
| Dark: median Sleep-bout duration (s) | −30.00 | −155.53 to 95.53 | −0.541 | 9 | 0.602 |
| Light: bidirectional Sleep↔Wake transitions (transitions per valid recording hour) | +4.51 | 1.97 to 7.04 | 4.027 | 9 | 0.00299 |
| Dark: bidirectional Sleep↔Wake transitions (transitions per valid recording hour) | +3.45 | 0.63 to 6.27 | 2.769 | 9 | 0.0218 |
| Light: median Wake-bout duration (s) | −11.71 | −17.90 to −5.53 | −4.282 | 9 | 0.00204 |
| Dark: median Wake-bout duration (s) | −34.86 | −52.53 to −17.18 | −4.461 | 9 | 0.00158 |

Part B gives SD-minus-Ctrl contrasts from two-sided equal-variance Student t tests (Ctrl, n = 4; SD, n = 7), with pointwise 95% CIs and P values. The first eight rows cover 24 h, followed by separate light- and dark-phase comparisons. Sleep combines NREM and REM. Each transition-rate label specifies the denominator in hours of valid recording or hours of sleep.

**Part C. Absolute waking EEG power and wheel activity**

| Band | Ctrl ($\mu V^2$) | SD ($\mu V^2$) | SD/Ctrl (P) | A/I (P) |
|---|---|---|---|---|
| δ | 540.0 | 1,215.9 | 2.25 (0.0242) | 1.61 (0.0039) |
| θ | 175.2 | 257.9 | 1.47 (0.2909) | 1.24 (0.0039) |
| σ | 73.5 | 67.4 | 0.92 (0.4424) | 1.10 (0.0322) |
| β | 103.1 | 86.2 | 0.84 (0.2909) | 1.17 (0.0039) |

| Band | SD/Ctrl, inactive (P) | SD/Ctrl, active (P) | A/I, SD/Ctrl (P) |
|---|---|---|---|
| δ | 2.35 (0.0242) | 1.64 (0.4545) | 0.69 (0.3333) |
| θ | 1.32 (0.7455) | 1.38 (0.6061) | 1.04 (0.6424) |
| σ | 0.81 (0.1818) | 0.95 (0.7455) | 1.18 (0.0970) |
| β | 0.87 (0.7455) | 0.71 (0.0424) | 0.82 (0.0970) |

Group means are geometric means ($\mu V^2$). Each ratio is followed by its Holm-adjusted P value. For each mouse, A/I is the ratio of mean waking band power in wheel-active versus wheel-inactive minutes. The upper table gives the geometric mean of this ratio across all 11 mice. The final column of the lower table gives the ratio of the SD and Ctrl geometric means (SD/Ctrl). Tests used mice as the unit (Ctrl, 4; SD, 7). See S9.3 for permutation procedures and adjustment families.

**Part D. Changes in wheel-activity rhythm indices**

| Figure S12 measure | Ctrl change (n = 4) | SD change (n = 7) | SD − Ctrl [95% CI] | P value |
|---|---|---|---|---|
| A. Dark-phase rotation share (pp) | +5.04 ± 2.48 | −3.59 ± 7.65 | −8.63 [−32.41, 15.16] | 0.433 |
| B. Dark-minus-light active-minute occupancy (pp) | +5.83 ± 4.97 | −2.50 ± 8.55 | −8.33 [−35.75, 19.08] | 0.509 |
| C. Interdaily stability (IS) | +0.031 ± 0.039 | −0.300 ± 0.057 | −0.331 [−0.517, −0.145] | 0.00297 |
| D. Intradaily variability (IV) | +0.032 ± 0.118 | +0.432 ± 0.100 | +0.400 [0.0386, 0.762] | 0.0337 |
| E. Relative amplitude (RA) | +0.005 ± 0.005 | −0.152 ± 0.056 | −0.157 [−0.329, 0.014] | 0.0680 |
| F. Fixed-24-h single-harmonic fit ($R^2$) | +0.021 ± 0.038 | −0.452 ± 0.062 | −0.473 [−0.672, −0.273] | 0.000453 |

Change is the Days 13–15 value minus the Days 7–9 value for each mouse. Group changes are arithmetic means ± SEM. SD − Ctrl is the difference in mean change with its pointwise 95% confidence interval. P values are from two-sided equal-variance Student t tests on individual changes (df = 9), one per outcome (Supplementary Methods S10). pp = percentage points.

**Supplementary Table S3. Total wheel activity, wheel-defined activity-state measures, and model-based comparisons during the dark-phase analysis of Days 15–17. Outcomes correspond to Figure 5B–F.**

Part A. Descriptive statistics

| Outcome | Ctrl/Sham | Ctrl/tUS | SD/Sham | SD/tUS |
|---|---|---|---|---|
| Dark-phase total wheel activity (rotations/dark phase) | 14,381.9 | 12,874.6 | 10,054.2 | 7,540.6 |
| P(Inactivity→Running) | 0.09147 | 0.09146 | 0.22434 | 0.13166 |
| P(Running→Inactivity) | 0.06815 | 0.07562 | 0.14011 | 0.15349 |
| Median inactivity bout (min) | 5.06 | 3.63 | 2.48 | 3.81 |
| Running intensity (rotations/active min) | 35.71 | 33.78 | 23.04 | 22.56 |

Part B. Model-based comparisons

| Outcome | Transformation | Comparison | Estimate | 95% CI | Nominal P | Holm P |
|---|---|---|---|---|---|---|
| Dark-phase total wheel activity (rotations/dark phase) | log(1 + x) | SD condition × tUS interaction | −0.17172<br>t = −0.884 | −0.56752 to 0.22407 | 0.3834 | — |
| Dark-phase total wheel activity (rotations/dark phase) | log(1 + x) | tUS − Sham within Ctrl | −0.12166<br>t = −0.817 | −0.42484 to 0.18151 | 0.4197 | 0.4197 |
| Dark-phase total wheel activity (rotations/dark phase) | log(1 + x) | tUS − Sham within SD | −0.29339<br>t = −2.349 | −0.54783 to −0.03895 | 0.0252 | 0.0504 |
| Dark-phase total wheel activity (rotations/dark phase) | log(1 + x) | SD/Sham − Ctrl/Sham | −0.34968<br>t = −2.707 | −0.61283 to −0.08653 | 0.0108 | 0.0324 |
| P(Inactivity→Running) | arcsin(sqrt(p)) | SD condition × tUS interaction | −0.11710<br>t = −2.613 | −0.20838 to −0.02582 | 0.0136 | — |
| P(Inactivity→Running) | arcsin(sqrt(p)) | tUS − Sham within Ctrl | −0.00288<br>t = −0.081 | −0.07497 to 0.06920 | 0.9356 | 0.9356 |
| P(Inactivity→Running) | arcsin(sqrt(p)) | tUS − Sham within SD | −0.11998<br>t = −4.365 | −0.17598 to −0.06399 | 0.000125 | 0.000249 |
| P(Inactivity→Running) | arcsin(sqrt(p)) | SD/Sham − Ctrl/Sham | +0.18806<br>t = 5.613 | 0.11981 to 0.25631 | 3.34e-06 | 1e-05 |
| P(Running→Inactivity) | arcsin(sqrt(p)) | SD condition × tUS interaction | +0.00555<br>t = 0.241 | −0.04147 to 0.05257 | 0.8115 | — |
| P(Running→Inactivity) | arcsin(sqrt(p)) | tUS − Sham within Ctrl | +0.01310<br>t = 0.959 | −0.01473 to 0.04094 | 0.3448 | 0.6470 |
| P(Running→Inactivity) | arcsin(sqrt(p)) | tUS − Sham within SD | +0.01865<br>t = 1.003 | −0.01924 to 0.05655 | 0.3235 | 0.6470 |
| P(Running→Inactivity) | arcsin(sqrt(p)) | SD/Sham − Ctrl/Sham | +0.11887<br>t = 8.852 | 0.09151 to 0.14622 | 4.1e-10 | 1.23e-09 |

| Outcome | Transformation | Comparison | Estimate | 95% CI | Nominal P | Holm P |
|---|---|---|---|---|---|---|
| Median inactivity bout (min) | log(1 + duration) | SD condition × tUS interaction | +0.60282<br>t = 2.692 | 0.14665 to 1.05898 | 0.0112 | — |
| Median inactivity bout (min) | log(1 + duration) | tUS − Sham within Ctrl | −0.26206<br>t = −1.352 | −0.65686 to 0.13274 | 0.1858 | 0.1858 |
| Median inactivity bout (min) | log(1 + duration) | tUS − Sham within SD | +0.34076<br>t = 3.037 | 0.11224 to 0.56928 | 0.00472 | 0.0130 |
| Median inactivity bout (min) | log(1 + duration) | SD/Sham − Ctrl/Sham | −0.51659<br>t = −3.070 | −0.85931 to −0.17387 | 0.0043 | 0.0130 |
| Running intensity (rotations/active min) | identity | SD condition × tUS interaction | +1.44801<br>t = 0.404 | −5.85285 to 8.74886 | 0.6889 | — |
| Running intensity (rotations/active min) | identity | tUS − Sham within Ctrl | −1.93142<br>t = −0.776 | −7.00297 to 3.14013 | 0.4436 | 0.8872 |
| Running intensity (rotations/active min) | identity | tUS − Sham within SD | −0.48341<br>t = −0.187 | −5.73526 to 4.76843 | 0.8525 | 0.8872 |
| Running intensity (rotations/active min) | identity | SD/Sham − Ctrl/Sham | −12.66700<br>t = −4.900 | −17.93264 to −7.40136 | 2.66e-05 | 7.97e-05 |

Part A reports untransformed arithmetic group means of mouse-level measures averaged across the dark phases (ZT12–24) of Days 15–17 (Ctrl/Sham, Ctrl/tUS, SD/Sham, and SD/tUS; n = 9 per group). Part B reports estimates and pointwise nominal 95% confidence intervals from two-factor linear models with HC3 standard errors (residual df = 32), on the log(1 + x) scale for total rotations and median inactivity-bout duration, the arcsine-square-root scale for both transition probabilities, and the original scale for running intensity. Holm adjustment applied to the three group contrasts within each outcome (Supplementary Methods S8).

Supplementary Table S4. Novel-object-recognition and open-field outcomes: descriptive statistics and model-based comparisons.

Part A: Descriptive statistics

| Outcome | Ctrl/Sham | Ctrl/tUS | SD/Sham | SD/tUS |
|---|---|---|---|---|
| NORT discrimination index | 0.1869 | 0.2166 | −0.0067 | 0.3601 |
| OFT center time (s) | 107.83 | 136.01 | 52.36 | 102.08 |
| OFT total distance (m) | 83.15 | 79.76 | 56.43 | 66.79 |

Part B: Model-based comparisons

| Outcome | Transformation | Comparison | Estimate | 95% CI | Nominal P | Holm P |
|---|---|---|---|---|---|---|
| NORT discrimination index | identity | SD condition × tUS interaction | +0.3370<br>t = 1.496 | −0.1246 to 0.7987 | 0.1459 | — |
| NORT discrimination index | identity | tUS − Sham within Ctrl | +0.0298<br>t = 0.188 | −0.2950 to 0.3545 | 0.8524 | 0.8524 |
| NORT discrimination index | identity | tUS − Sham within SD | +0.3668<br>t = 2.290 | 0.0388 to 0.6949 | 0.0297 | 0.1189 |
| NORT discrimination index | identity | SD/Sham − Ctrl/Sham | −0.1935<br>t = −1.106 | −0.5521 to 0.1650 | 0.2783 | 0.8349 |
| NORT discrimination index | identity | SD/tUS − Ctrl/Sham | +0.1733<br>t = 1.014 | −0.1767 to 0.5233 | 0.3193 | 0.8349 |
| OFT center time (s) | identity | SD condition × tUS interaction | +21.55<br>t = 0.642 | −47.20 to 90.29 | 0.5261 | — |
| OFT center time (s) | identity | tUS − Sham within Ctrl | +28.18<br>t = 0.988 | −30.23 to 86.59 | 0.3315 | 0.6630 |
| OFT center time (s) | identity | tUS − Sham within SD | +49.73<br>t = 2.810 | 13.47 to 85.98 | 0.00894 | 0.0294 |
| OFT center time (s) | identity | SD/Sham − Ctrl/Sham | −55.48<br>t = −2.891 | −94.79 to −16.17 | 0.00734 | 0.0294 |
| OFT center time (s) | identity | SD/tUS − Ctrl/Sham | −5.7528<br>t = −0.281 | −47.6197 to 36.1140 | 0.7804 | 0.7804 |
| OFT total distance (m) | identity | SD condition × tUS interaction | +13.75<br>t = 1.356 | −7.02 to 34.52 | 0.1858 | — |
| OFT total distance (m) | identity | tUS − Sham within Ctrl | −3.39<br>t = −0.535 | −16.35 to 9.57 | 0.5966 | 0.5966 |
| OFT total distance (m) | identity | tUS − Sham within SD | +10.36<br>t = 1.308 | −5.86 to 26.59 | 0.2014 | 0.4029 |
| OFT total distance (m) | identity | SD/Sham − Ctrl/Sham | −26.72<br>t = −3.864 | −40.88 to −12.55 | 0.000604 | 0.0024 |
| OFT total distance (m) | identity | SD/tUS − Ctrl/Sham | −16.3539<br>t = −2.127 | −32.1024 to −0.6055 | 0.0424 | 0.1271 |

Part A reports untransformed arithmetic group means for NORT discrimination index, OFT center time, and OFT total distance (Ctrl/Sham and Ctrl/tUS, n = 7 each; SD/Sham and SD/tUS, n = 9 each). Part B reports estimates and pointwise nominal 95% confidence intervals on the original outcome scales from two-factor linear models with HC3 standard errors (residual df = 28). Holm adjustment covered the four group contrasts for each outcome, including SD/tUS − Ctrl/Sham (Supplementary Methods S8).

Supplementary Table S4, Part C. Wheel activity before behavioral testing

| Wheel measure | Ctrl/Sham (n = 7) | Ctrl/tUS (n = 7) | SD/Sham (n = 9) | SD/tUS (n = 9) |
|---|---|---|---|---|
| Total rotations per dark phase | 14,463.50 [13,618.25, 15,723.50] | 13,783.50 [12,964.50, 14,324.00] | 9,304.00 [8,749.50, 9,829.50] | 6,932.00 [6,239.00, 8,963.50] |
| Running intensity (rotations/active min) | 35.91 [34.81, 37.05] | 35.25 [32.58, 37.31] | 21.58 [18.17, 26.40] | 21.02 [19.52, 23.57] |
| Activity-resumption probability | 0.093 [0.072, 0.116] | 0.082 [0.062, 0.120] | 0.198 [0.173, 0.324] | 0.127 [0.117, 0.136] |
| Median inactivity-bout duration (min) | 4.50 [3.38, 4.75] | 3.50 [2.88, 4.25] | 2.50 [2.00, 3.00] | 4.00 [3.00, 4.00] |
| Activity–inactivity switches per hour | 4.83 [4.10, 5.29] | 4.50 [3.88, 5.52] | 9.75 [9.50, 11.25] | 8.21 [6.88, 9.25] |

For each mouse, nightly values were averaged across the dark phases (ZT12–24) of Days 15 and 16. Values are group medians [25th, 75th percentiles] of these two-night means. Percentiles were calculated by linear interpolation.

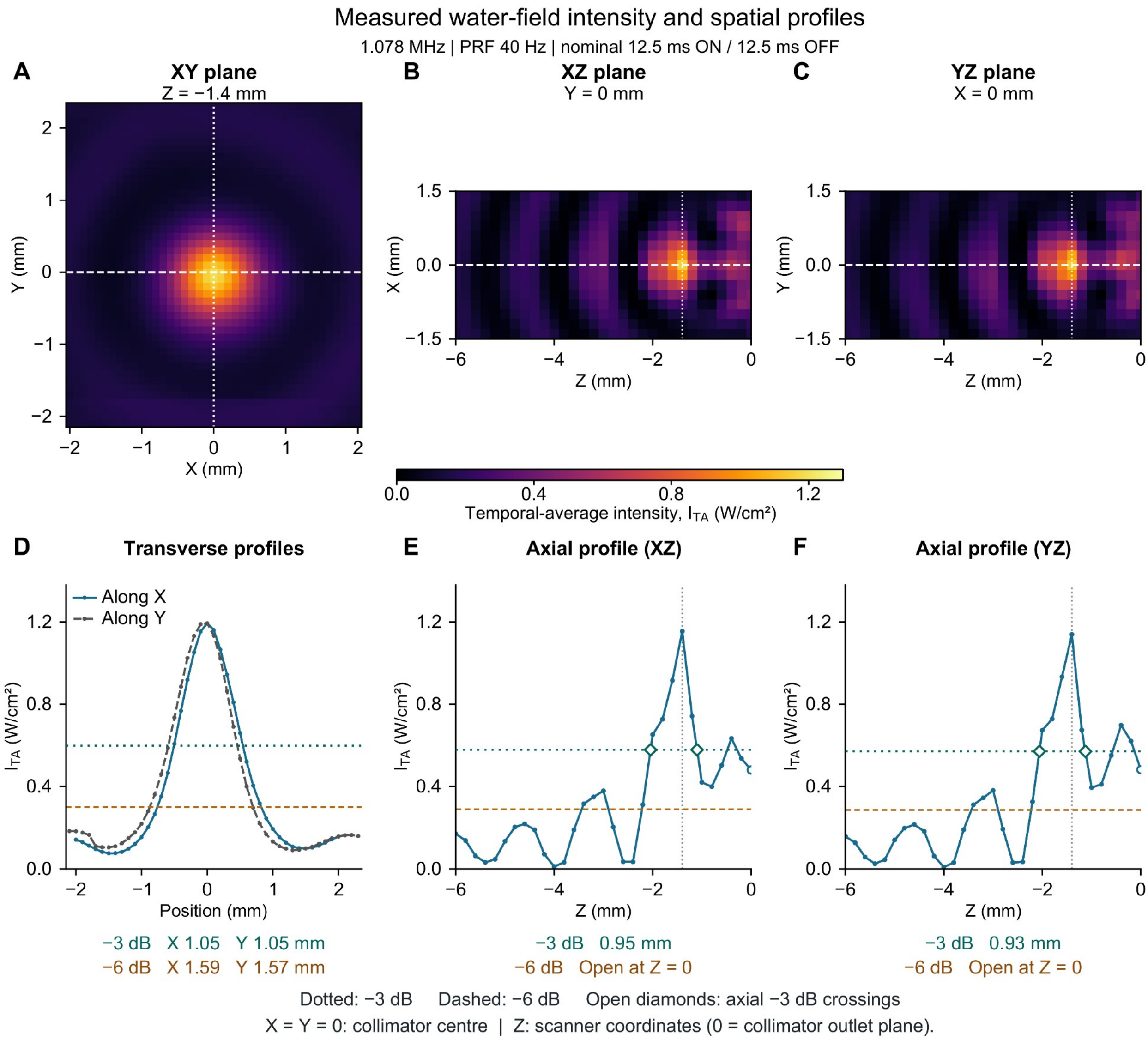


**Figure S1. Measured water-field intensity maps and spatial profiles.**

(A–C) Hydrophone measurements of temporal-average intensity (I_TA) in the XY plane at Z = −1.4 mm (A), XZ plane at Y = 0 mm (B), and YZ plane at X = 0 mm (C). Measurements used approximately 1.078 MHz, a pulse repetition frequency of 40 Hz, and nominal 12.5-ms ON and OFF intervals. I_TA was calculated over complete pulse-repetition intervals, including OFF periods. Maps share an absolute intensity scale and show measured cells without smoothing or interpolation. The source data comprise 1,845 XY positions at 0.1-mm spacing and 651 positions in each axial plane at 0.2-mm spacing.

X = 0 and Y = 0 denote the collimator center. The vertical display ranges in B and C are −1.5 to +1.5 mm; the Z axes in B, C, E, and F span −6 to 0 mm, with scanner Z = 0 mm aligned with the collimator outlet plane. White dashed lines identify the profiles below; white dotted lines mark the remaining peak coordinates. All measured positions are retained in the source data.

(D) X- and Y-direction profiles through the XY-plane intensity maximum, plotted relative to the collimator center. (E, F) Z-direction profiles at X = 0 and Y = 0 mm, extracted separately from XZ (E) and YZ (F). The axial intensity peaks were at Z = −1.4 mm. Points are measurements joined for visualization. Teal dotted and orange dashed lines indicate the −3 and −6 dB intensity thresholds, respectively. In D, the −3 dB widths were 1.05 mm along both X and Y; the −6 dB widths were 1.59 and 1.57 mm, respectively. In E and F, open diamonds mark the −3 dB crossings enclosing the main axial peak (widths, 0.95 and 0.93 mm, respectively); separate suprathreshold intervals nearer the outlet were excluded. Both axial profiles remained above the −6 dB threshold at Z = 0 mm, so their complete −6 dB widths could not be determined.

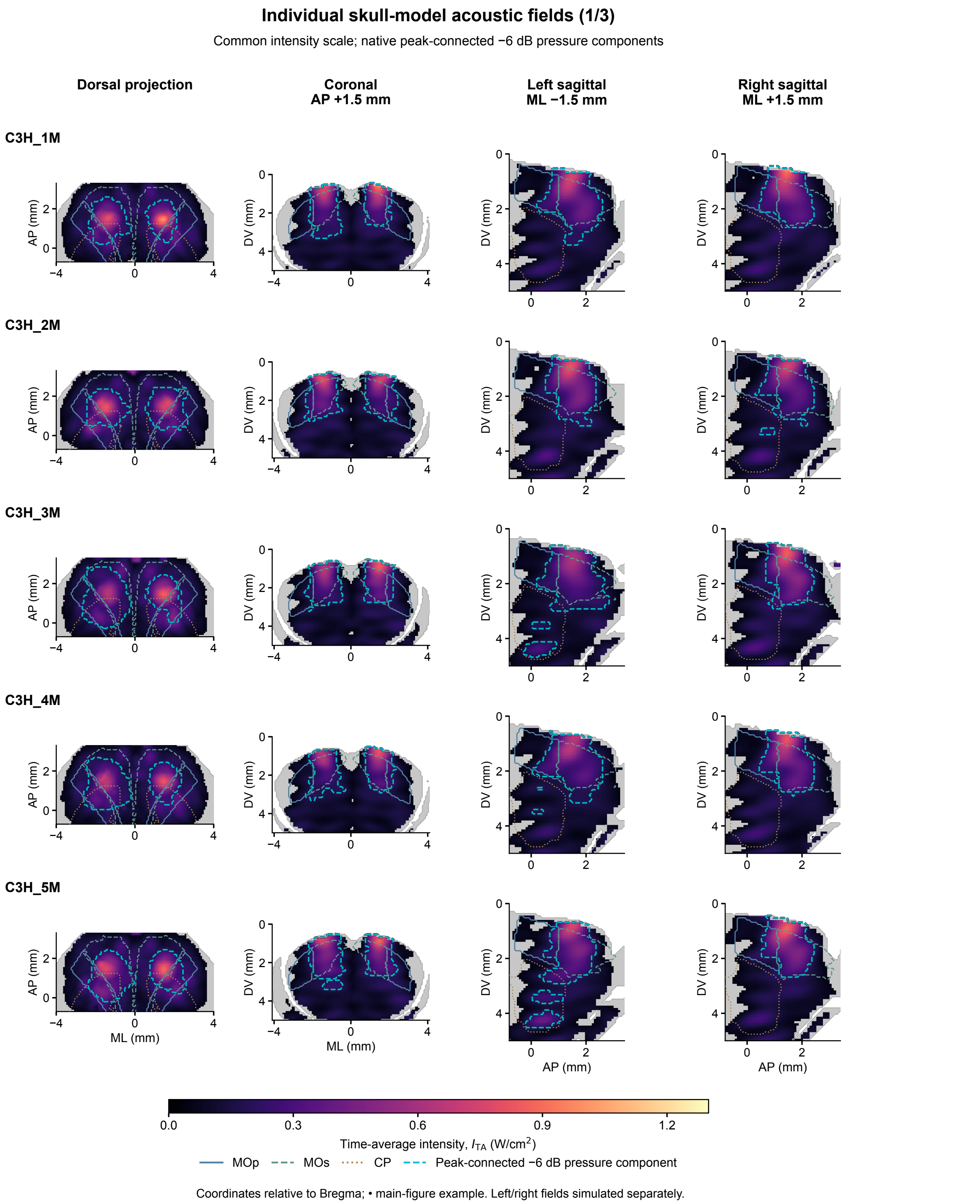


**Figure S2. Individual skull-model predictions of transcranial acoustic exposure. (1/3)**

Each row shows one of ten postnatal day 90 male C3H/HeJ skull models, with five models per page. Columns show a dorsal maximum projection, a coronal section at anteroposterior +1.5 mm, a left sagittal section at mediolateral −1.5 mm, and a right sagittal section at mediolateral +1.5 mm, relative to bregma. Left and right exposures were simulated independently with duty factor 0.5. Dorsal and coronal maps display the higher of the two independently modeled intensities at each location. Each sagittal map displays the field for the corresponding stimulation side. These displays do not represent simultaneous irradiation or intensity summation.

All models use the same axes, with coronal and sagittal views spanning 0–5.0 mm ventral to the bregma plane, the same intensity scale (0–1.3 W/cm²), and the same display threshold: temporal-average intensities below 0.0455 W/cm² are hidden for display only. Gray indicates the modeled intracranial support of each skull. MOp, MOs, and CP outlines use the existing approximate atlas registration for that skull. Cyan outlines show the projection or section of the native, peak-connected −6-dB pressure-amplitude components. For each stimulation side, the component was defined within physical brain plus effective CSF by $|\hat{p}| \geq 10^{-6/20} |\hat{p}|_{\max}$ and six-neighbour connectivity to the pressure peak. Dorsal and coronal component masks combine the separately calculated left/right components by spatial union; each sagittal map uses its corresponding side. The cyan contours are individual-model component boundaries, not across-skull occupancy contours or thresholds of the displayed intensity maps. In the dorsal projection, a component is present at a planar location if it occurs at any depth along that projection.

Figure 2B–D shows the intensity maps for C3H_10M, and Figure 2E shows the corresponding 3D exposure regions. Figure 2F summarizes all ten skulls.

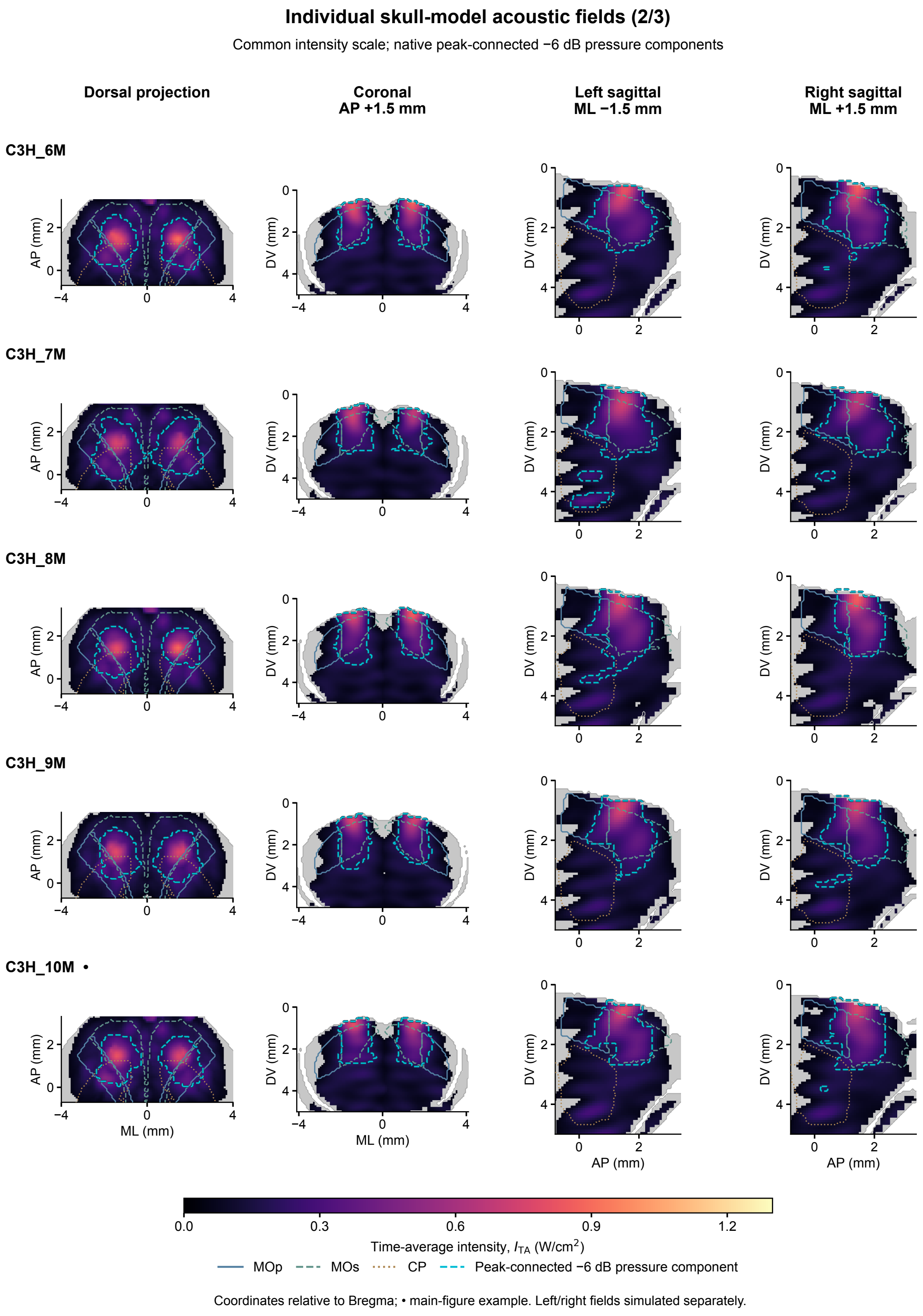


**Figure S2. Individual skull-model predictions of transcranial acoustic exposure. (2/3)**
The remaining five skull models are shown with the same axes, intensity scale, anatomical overlays, and pressure-component definitions as in (1/3).

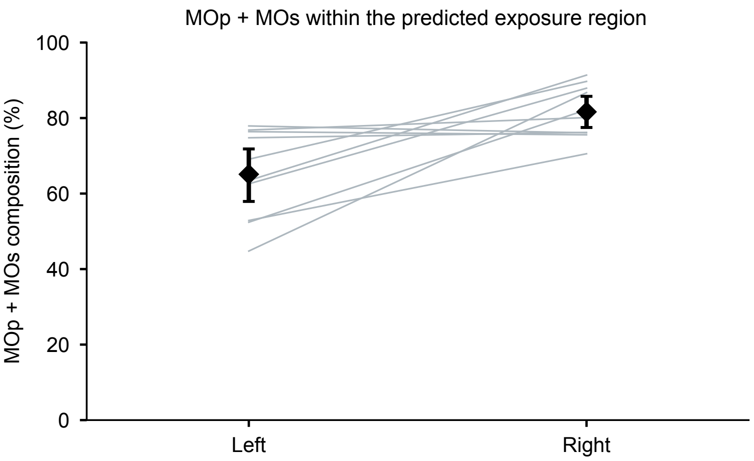


**Figure S2. Individual skull-model predictions of transcranial acoustic exposure. (3/3)**

MOp-plus-MOs composition of the predicted exposure region. The numerator counts MOp and MOs voxels; the denominator counts all nonzero atlas-annotated voxels within each retained peak-connected −6-dB component, including annotated fiber tracts. Circles denote left-sided fields and squares denote right-sided fields. Lines connect the two sides within each skull. Black diamonds show side-specific means and error bars show skull-bootstrap 95% confidence intervals (n = 10 skulls). The bilateral mean was 73.37 ± 6.12% (mean ± SD), averaging sides within each skull before cohort summarization. Side-specific motor-area composition is provided in motor_composition_by_side.csv. Coverage of each bilateral atlas region, expressed as a percentage of its total volume, is provided in region_coverage_by_skull.csv.

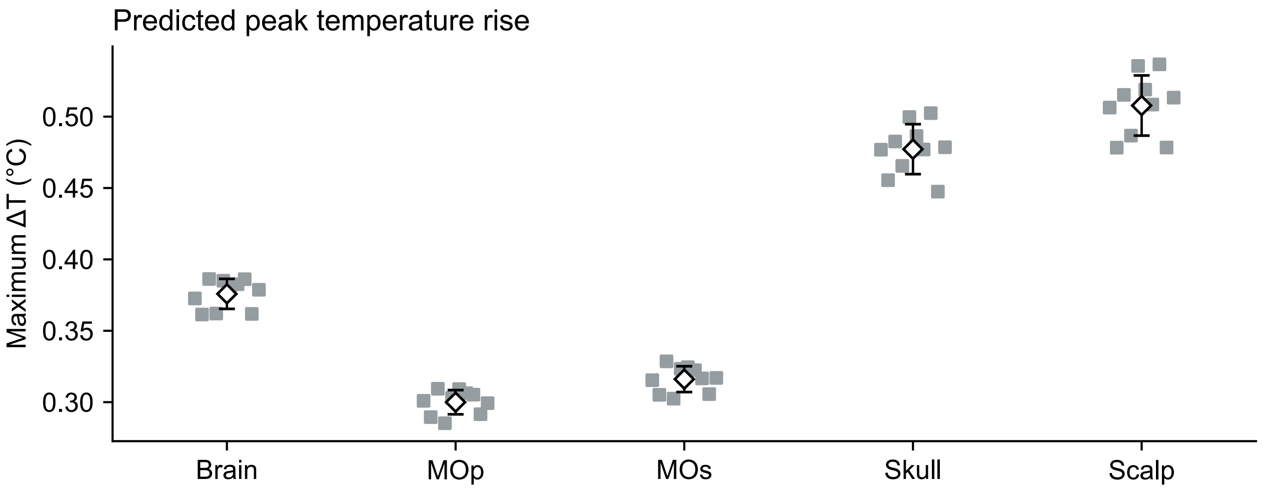


**Figure S3. Predicted tissue temperature changes under reference material assumptions.**
Maximum predicted temperature rise (ΔT) in the modeled brain, the bilateral primary motor area (MOp), the bilateral secondary motor area (MOs), skull, and effective dorsal scalp. Gray squares represent individual skull models, whereas open diamonds and error bars indicate mean ± standard deviation (n = 10 independent skull models). For each skull, the maximum ΔT across all voxels and simulated time points was determined separately for the left-first and right-first stimulation sequences, and the two maxima were averaged within each skull.

Temperature changes were calculated relative to matched no-sonication simulations.

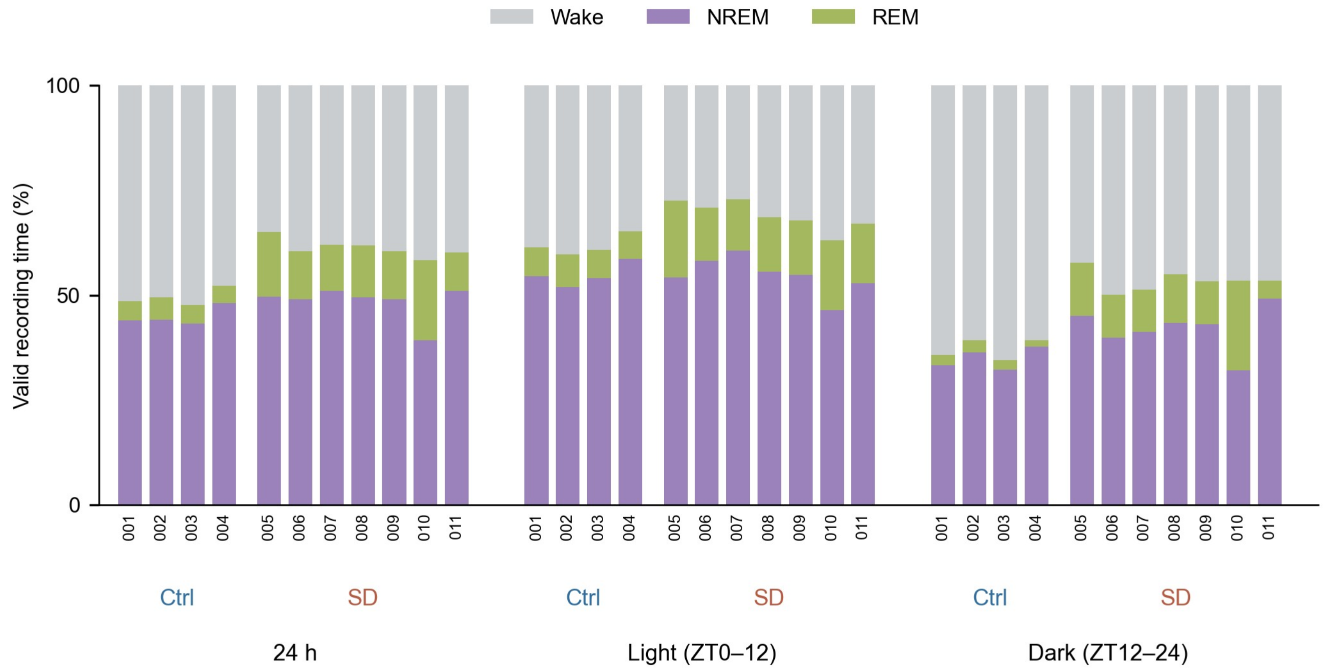


Figure S4. Individual vigilance-state composition under Ctrl and SD conditions.

Stacked bars show Wake, NREM and REM as percentages of valid EEG/EMG recording time for each mouse over 24 h (Day 17 ZT0 to Day 18 ZT0), and separately during the light (ZT0–12) and dark (ZT12–24) phases. Each bar represents one mouse. Within each phase, Ctrl mice appear in public-ID order EEG001–EEG004, followed by SD mice EEG005–EEG011; tick labels show the numeric part of each ID. Gray, purple and yellow-green indicate Wake, NREM and REM, respectively. Ctrl, n = 4; SD, n = 7. These independent mice received no tUS or sham procedures.

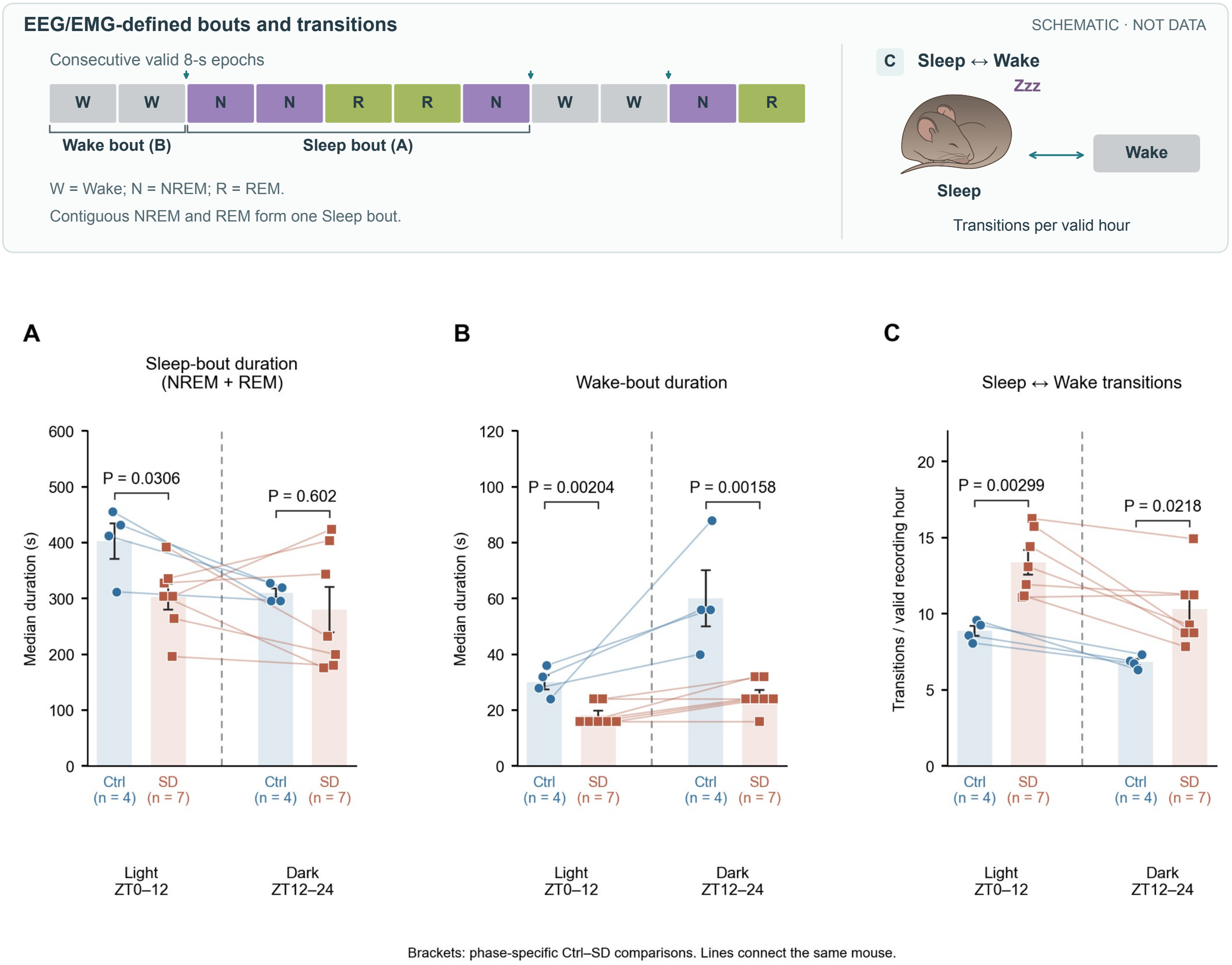


**Figure S5. Phase-specific sleep–wake bout duration and transitions.**

(A) Individual-mouse median Sleep-bout duration, with contiguous NREM and REM combined as Sleep. (B) Individual-mouse median Wake-bout duration. (C) Bidirectional Sleep↔Wake transitions per valid recording hour. Light denotes ZT0–12 and dark ZT12–24. Phase-edge bouts retain their observed within-phase lengths. Points show mice (Ctrl n = 4; SD n = 7), lines connect the same mouse, pale bars show group means, and error bars show SEM. Dashed vertical lines separate phases.

Brackets show P values from two-sided unpaired equal-variance Student t tests comparing Ctrl and SD separately within each phase. These independent mice received no tUS.

The schematic shows how scored EEG/EMG epochs define bouts and transitions between Sleep and Wake.

SciDraw illustrations adapted and recolored under CC BY 4.0 (https://creativecommons.org/licenses/by/4.0/). Sleep adapted from “Mouse Sleeping Curled” by Matias Andina (https://doi.org/10.5281/zenodo.7041090).

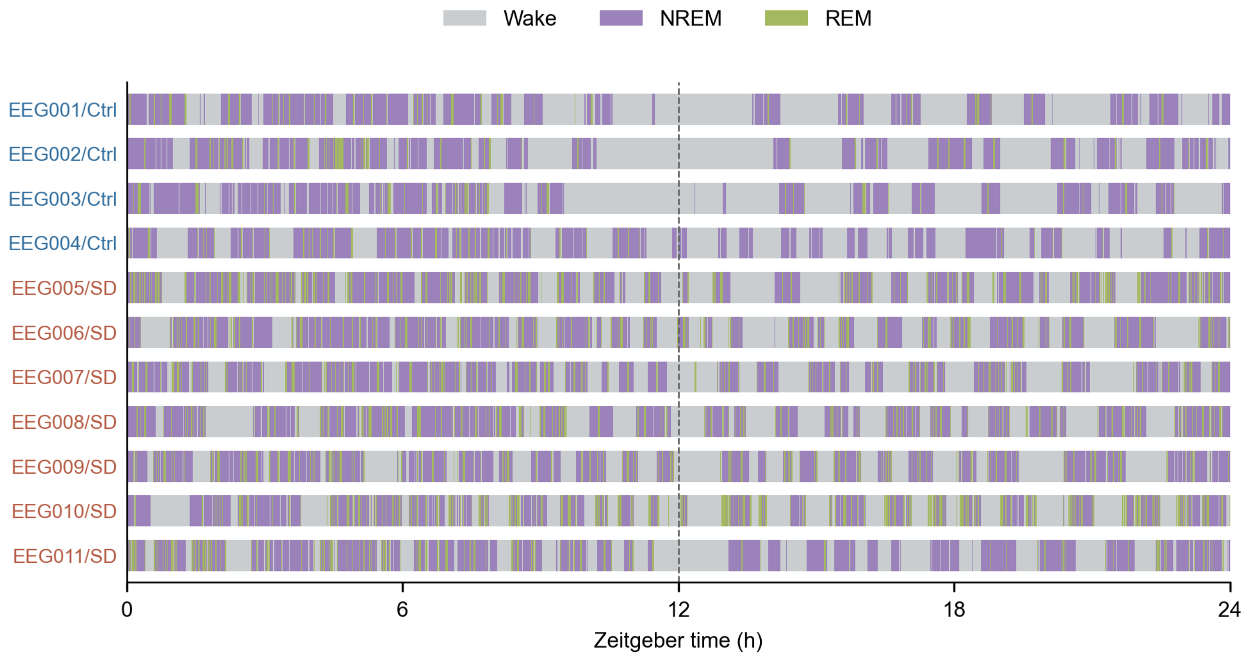


**Figure S6. Twenty-four-hour EEG/EMG-defined vigilance-state sequences in individual mice.**
The 24-h vigilance-state sequences are shown for all mice in the independent non-tUS cohort (Ctrl, n = 4; SD, n = 7). Wake, NREM and REM states were classified in consecutive 8-s epochs using FASTER2. Each horizontal row represents one mouse, with Ctrl mice displayed above SD mice. Gray, purple and yellow-green indicate Wake, NREM and REM, respectively. The vertical dashed line marks ZT12, separating the light (ZT0–12) and dark (ZT12–24) phases.

Colors show the 8-s vigilance-state classifications; brief episodes may be unresolved at the displayed scale.

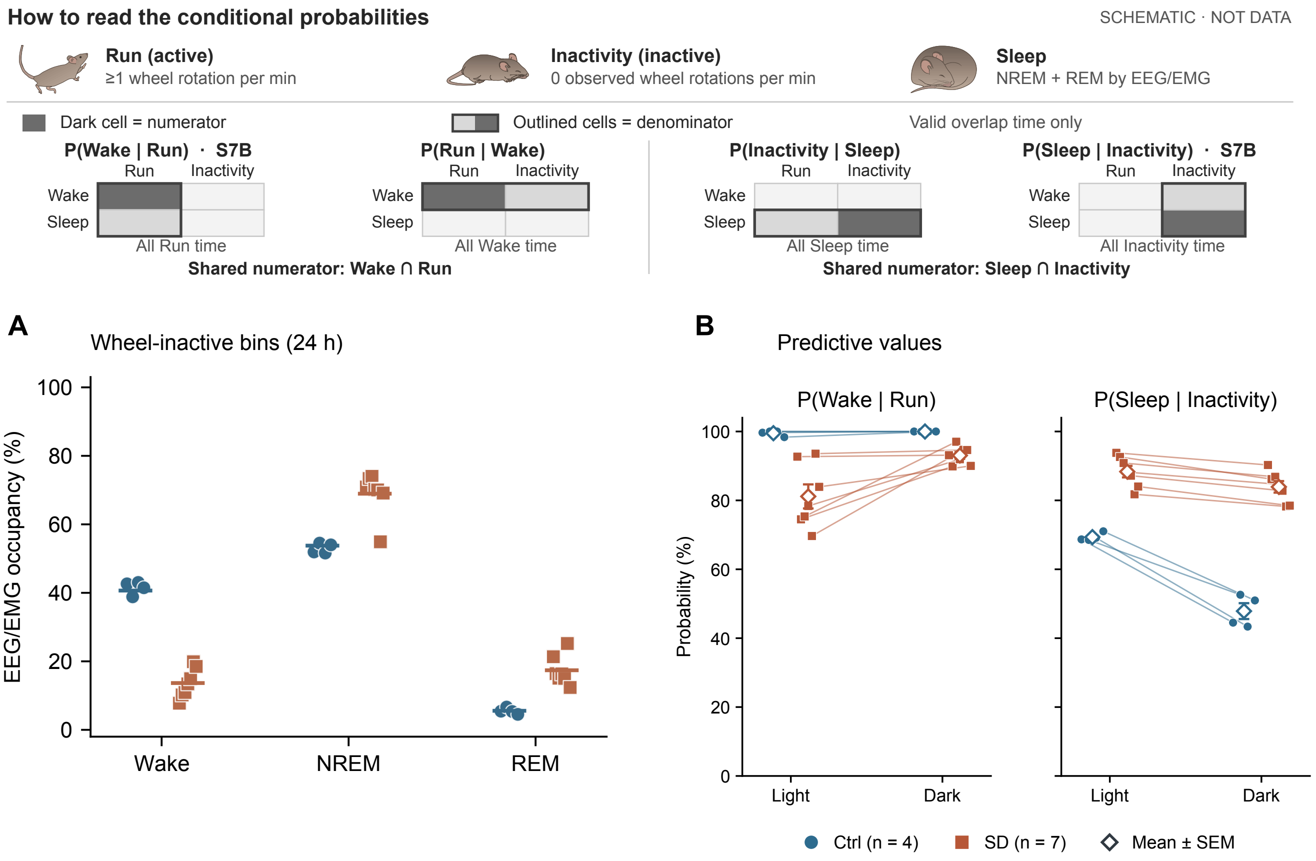


Figure S7. State composition during wheel inactivity and phase-specific wheel–vigilance correspondence.

(A) Wake, NREM and REM composition during wheel-inactive bins. The denominator was valid EEG/EMG time overlapping these bins. (B) Light- and dark-phase values for P(Wake | Run) and P(Sleep | Inactivity). Within each plot, a line connects the two observations from the same mouse. Open diamonds and error bars show group means ± SEM.

Blue circles indicate Ctrl (n = 4) and rust squares SD (n = 7). In A, small symbols show individual mice, and horizontal lines indicate group means. All mice belonged to the independent non-tUS experiment. Each mouse contributed equally. Both panels show descriptive mouse-level summaries.

The schematic shows the shared numerator and distinct denominators within each probability pair. S7B identifies the two probabilities plotted in panel B.

SciDraw illustrations adapted and recolored under CC BY 4.0 (https://creativecommons.org/licenses/by/4.0/). Run adapted from “Mouse on a Running Wheel” by Andrew Hardaway (https://doi.org/10.5281/zenodo.4742583). Inactivity adapted from “mouse sleeping” by Ethan Tyler and Lex Kravitz (https://doi.org/10.5281/zenodo.3925977). Sleep adapted from “Mouse Sleeping Curled” by Matias Andina (https://doi.org/10.5281/zenodo.7041090).

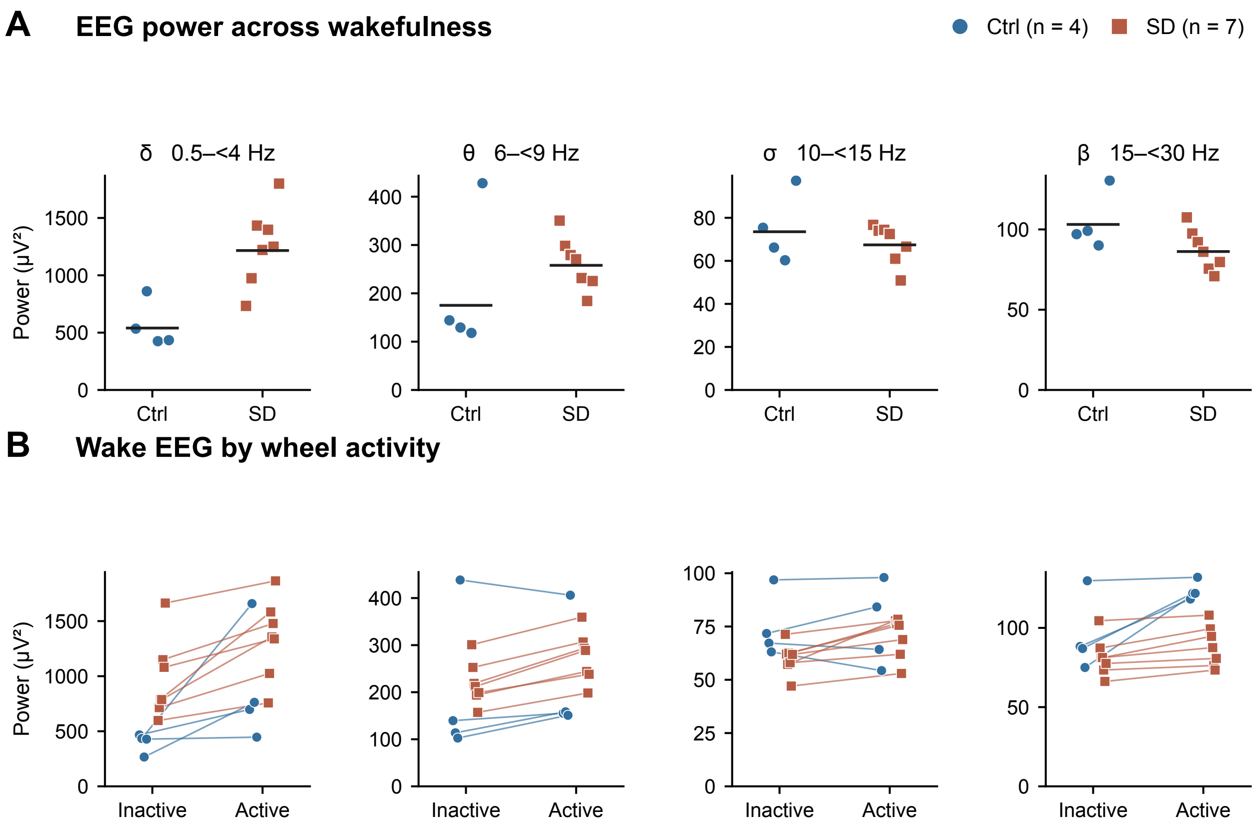


**Figure S8. Absolute waking EEG power and its association with wheel activity.**

(A) Each point represents the mean absolute EEG2 power across valid wake windows for one mouse. Horizontal bars indicate group geometric means. (B) Lines connect the same mouse across wheel-inactive and wheel-active minutes. Ctrl (blue circles, n = 4) and SD (rust squares, n = 7) mice received neither tUS nor sham. Each column represents the same band in both panels. Band limits and statistical comparisons are given in Supplementary Methods S9.3 and Table S2, Part C.

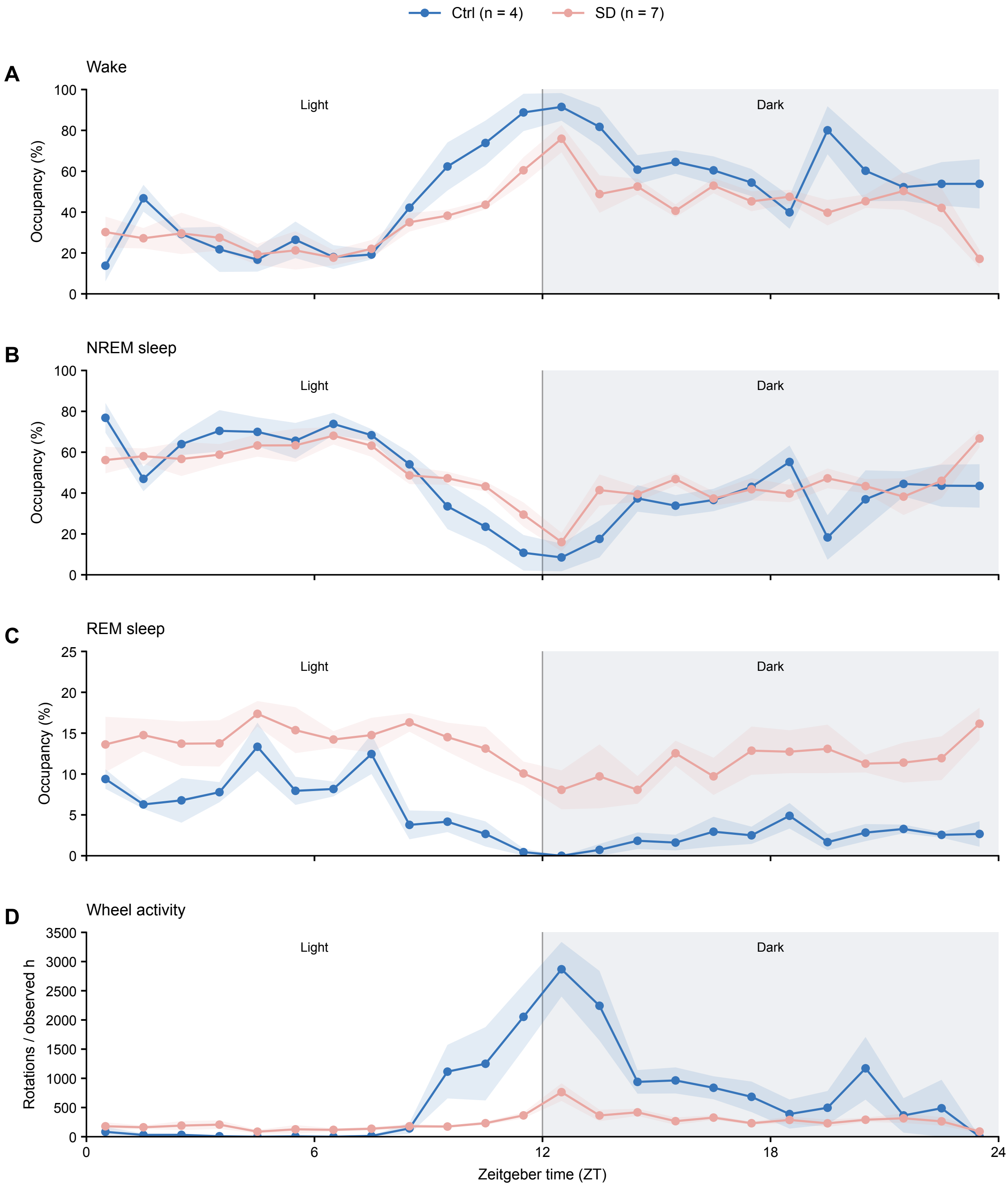


Figure S9. Matched 24-h vigilance and wheel profiles.

(A–C) Hourly Wake, NREM and REM occupancy during the Day 17 ZT0 to Day 18 ZT0 analysis window. Occupancy was calculated from the duration of valid vigilance-state records within each mouse-hour. (D) Wheel-rotation rate during the same 24-h window, calculated as the sum of observed counts in each mouse-hour divided by its valid recorded minutes and multiplied by 60. Valid wheel coverage was 54–60 min per mouse-hour; missing minutes were not replaced with zero. Panel D therefore reports a rate per 60 min of valid recording. Lines show group means and shading shows SEM across equally weighted mice (Ctrl n = 4, SD n = 7). Points are placed at the centers of identical 1-h ZT intervals in all four panels. Gray backgrounds denote the dark phase, ZT12–24. Wake and NREM use a 0–100% scale; REM uses a 0–25% scale to resolve its lower occupancy. Wheel rate retains its own units. No hourly significance tests are shown.

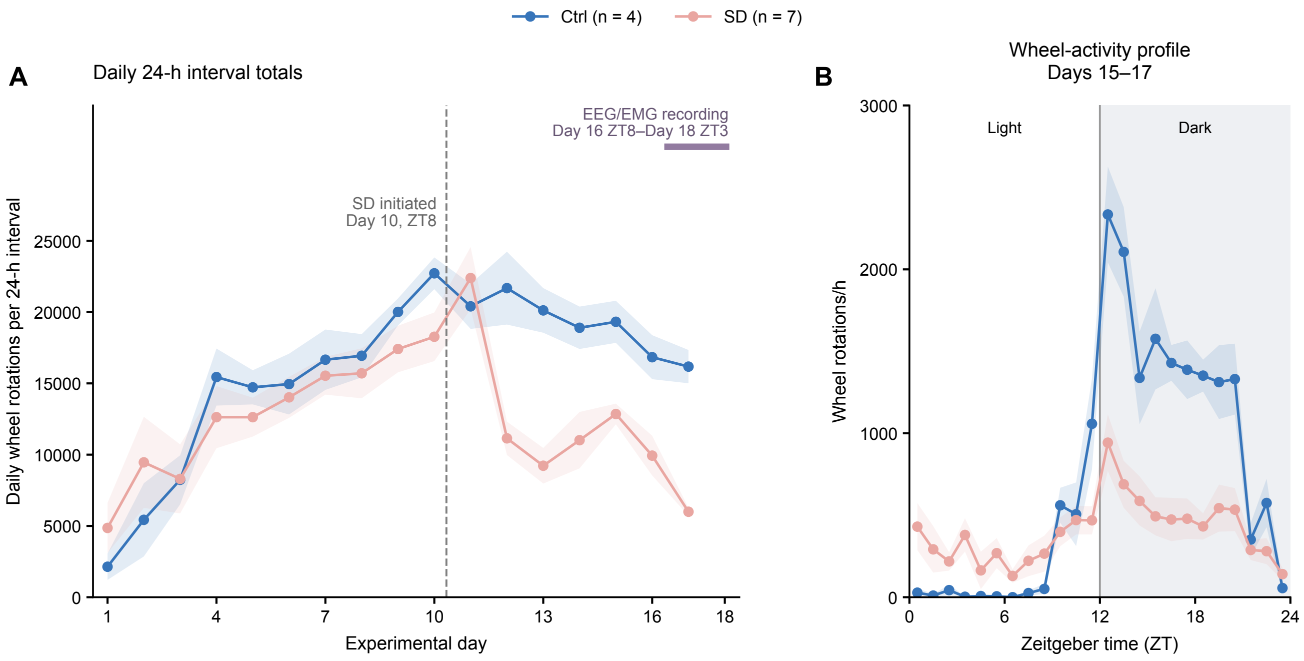


Figure S10. Longitudinal wheel activity and individual actograms.

(A) Daily observed wheel-rotation totals over 24-h intervals on Days 1–17. The vertical dashed line indicates SD onset at Day 10 ZT8. The purple bar indicates the EEG/EMG recording period from Day 16 ZT8 to Day 18 ZT3. The horizontal axis extends to Day 18 to show the recording end; wheel daily totals are plotted only through Day 17. (B) Hourly wheel-activity profiles averaged across Days 15–17 within each mouse before group summaries. Panel B summarizes three days. Supplementary Figure S9D shows the wheel rate for the single 24-h interval with simultaneous EEG/EMG recording (Day 17 ZT0 to Day 18 ZT0). Lines and shading indicate mean ± SEM (Ctrl n=4; SD n=7). Neither group received tUS or sham procedures. No day-wise or hourly significance tests are shown.

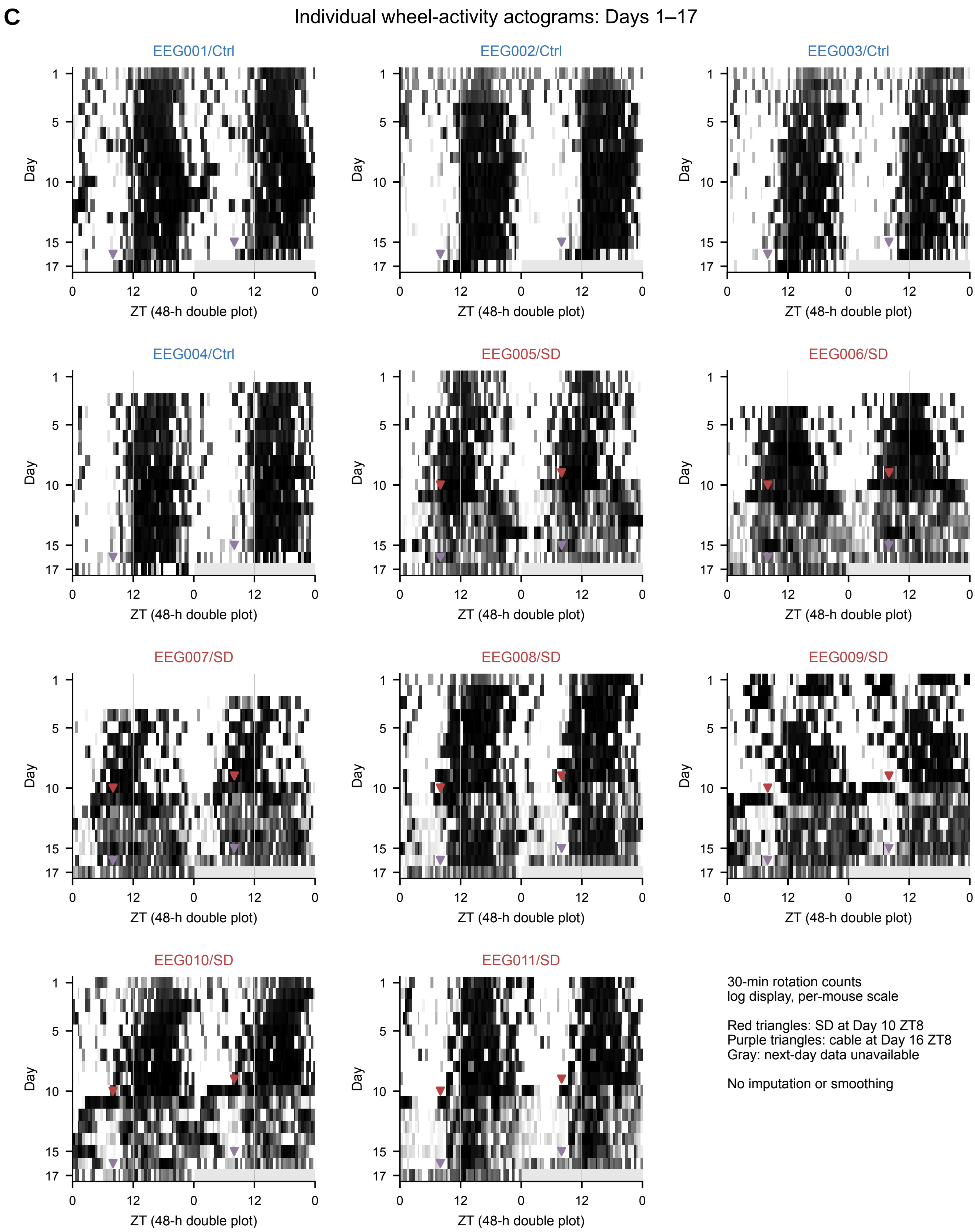


Figure S10 (continued). Individual actograms.

(C) Double-plotted 48-h actograms for all 11 mice. Pixels show observed 30-min rotation counts on a logarithmic scale referenced to each mouse's 95th percentile of positive counts. Shading is comparable only within a mouse. Red triangles mark SD onset (Day 10 ZT8) in SD mice. Purple triangles mark cable attachment (Day 16 ZT8). Double plotting shows each event twice. Gray marks unrecorded Day 18 in the final row. Missing counts were not imputed. Blue and red labels identify Ctrl (n = 4) and SD (n = 7) mice, respectively. Neither group received tUS or sham procedures.

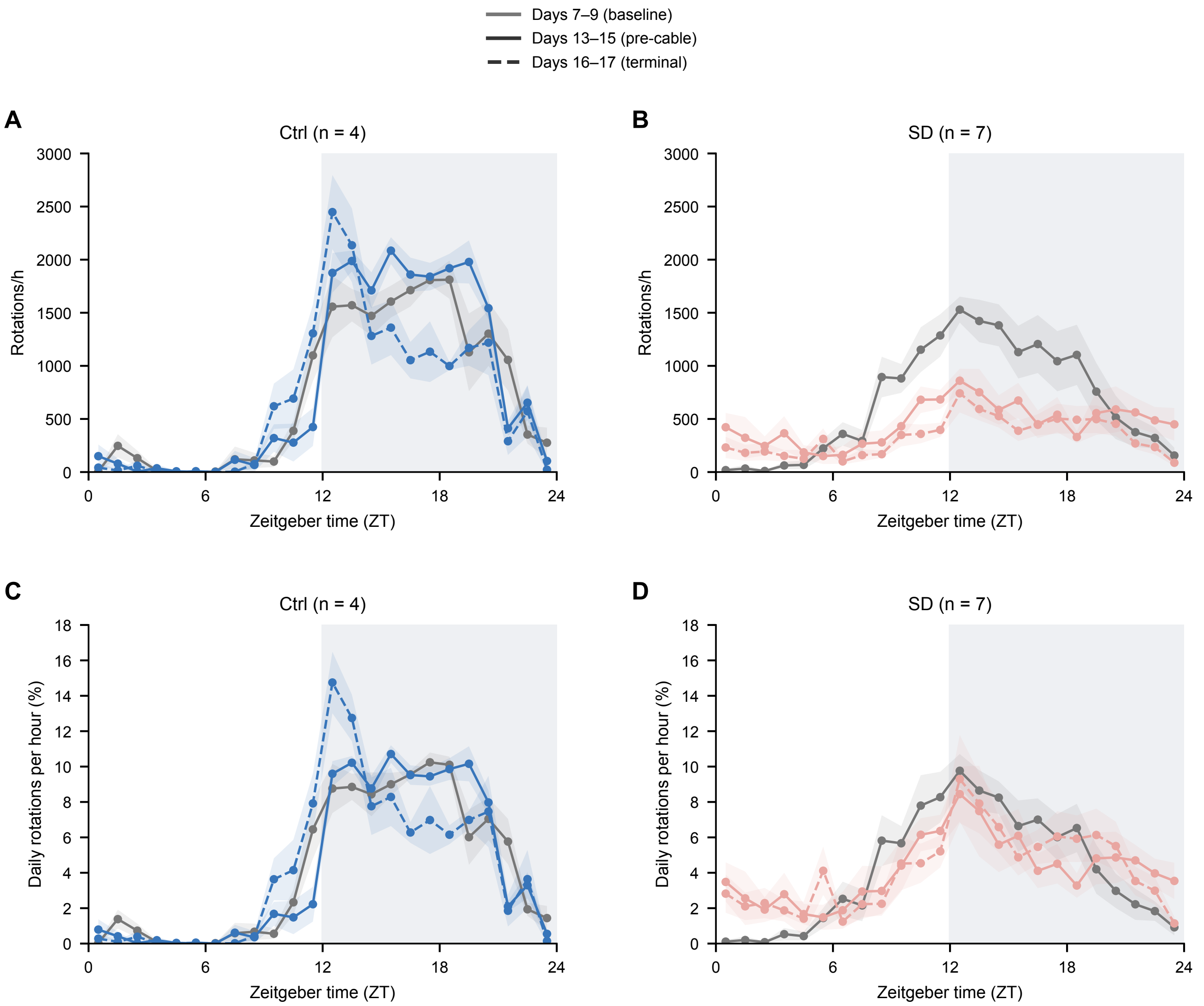


Figure S11. Absolute and relative 24-h profiles.

(A,B) Wheel rotations per hour. (C,D) Hourly rotation share as a percentage of each mouse's average daily total. Ctrl and SD are shown in separate columns with matching axes. Gray solid lines represent Days 7–9; colored solid lines represent Days 13–15; colored dashed lines represent Days 16–17. Lines show group means of the mouse-level values, and shading shows SEM across mice. Gray backgrounds identify the dark phase. The terminal 48 h include 8 h before cable attachment (Day 16 ZT8) and 40 h after attachment.

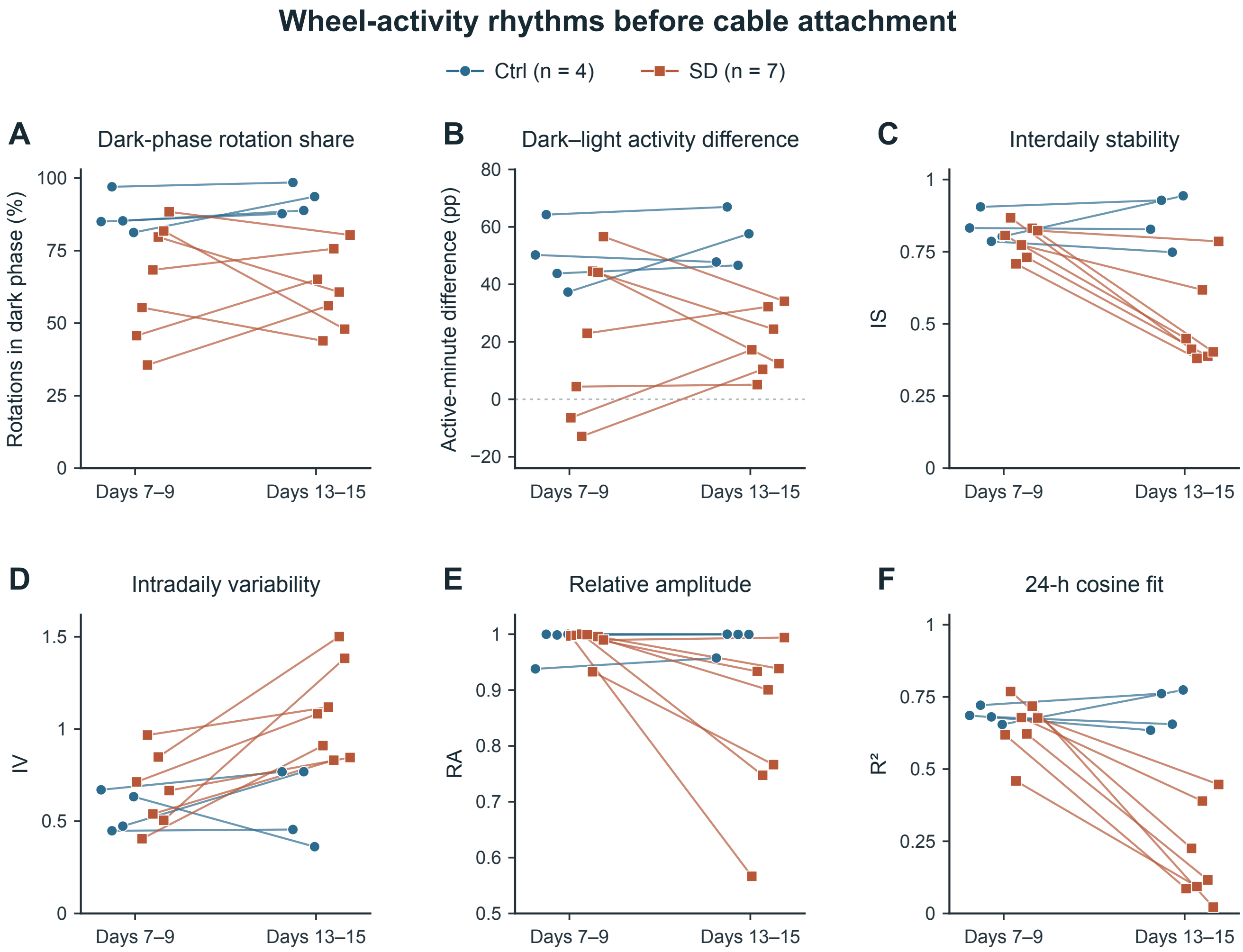


Figure S12. Individual wheel-activity rhythms across two 72-h windows.

Each line joins measurements from the same mouse. Days 7–9 were the last three complete days before SD mice moved to wheel-confined housing (Day 10, ZT8). Days 13–15 were the last three complete days before EEG/EMG cable attachment (Day 16, ZT8). Ctrl mice remained in their original cages. Both windows included three light and three dark phases. Panel letters correspond to the rows in Supplementary Table S2, Part D.

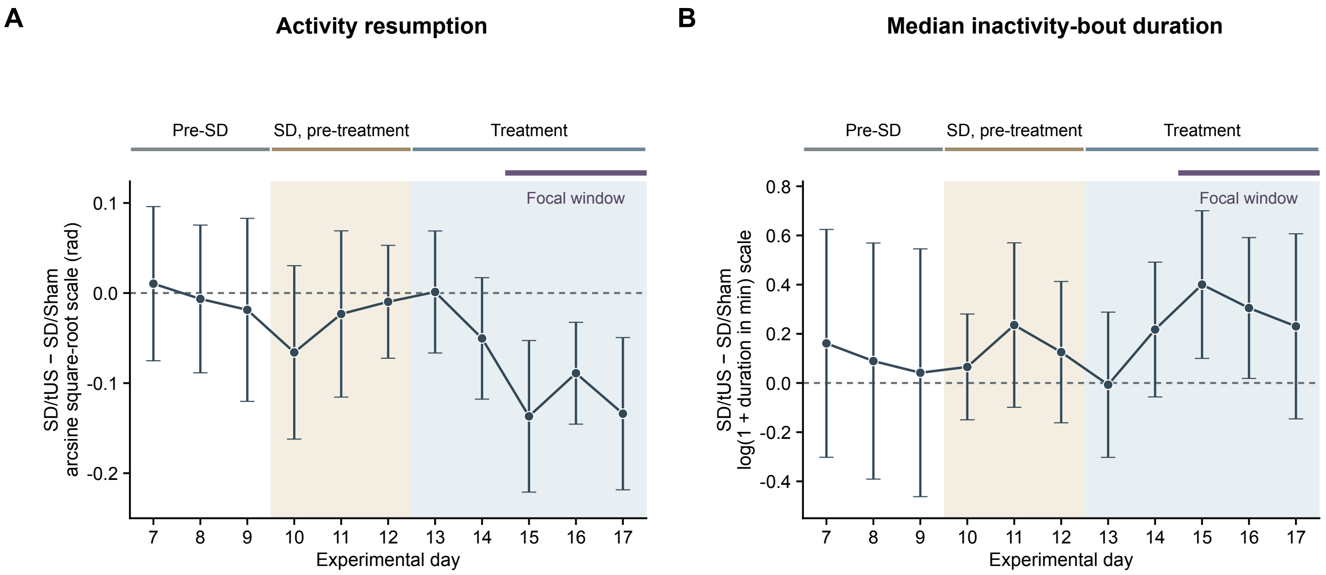


**Figure S13. Daily dark-phase differences in running pauses between SD/tUS and SD/Sham mice.**

Each point shows the estimated SD/tUS minus SD/Sham difference during one dark phase (ZT12–24), with the same nine mice per group across days. Error bars are pointwise 95% CIs. (A) Activity-resumption probability, P(I→R), on the arcsine square-root scale. (B) Median inactivity-bout duration on the log(1 + duration in minutes) scale. Zero denotes equal group means on the plotted scale. Groups are labeled by eventual assignment. SD housing began on Day 10 and treatment on Day 13. The upper bars mark these phases and the Days 15–17 window used in Figure 5, selected after inspection of the activity trajectories.

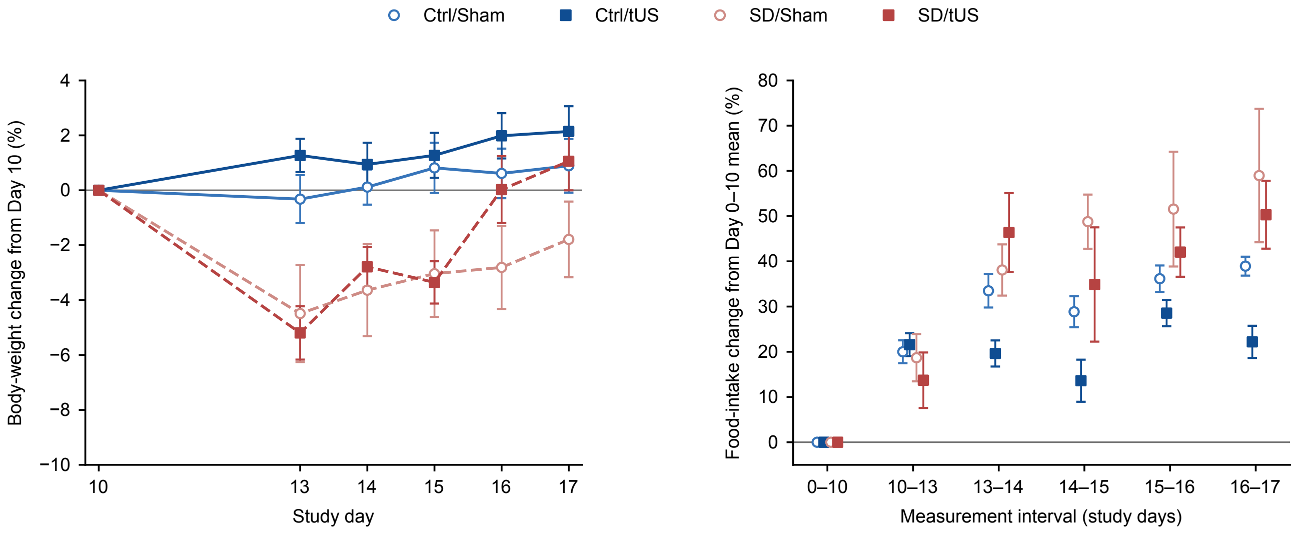


**Figure S14. Body-weight and food-intake changes in the wheel-activity intervention cohort.**

Body weight and food intake were monitored in the independent wheel-activity intervention cohort comprising four groups of mice: Ctrl/Sham, Ctrl/tUS, SD/Sham, and SD/tUS (n = 9 per group).

**Left:** Percentage change in body weight relative to each mouse's body weight on Day 10, shown for Days 10, 13, 14, 15, 16, and 17. Lines connect group means on measurement days.

**Right:** Percentage change in mean daily food intake relative to each mouse's average daily intake over Days 0–10. Values are shown for the measurement intervals Days 0–10, 10–13, 13–14, 14–15, 15–16, and 16–17. Intervals of different durations are presented as separate measurements and are not connected by lines.

Symbols and error bars indicate group means ± SEM, with nine mice contributing to each displayed group summary. Blue symbols indicate Ctrl groups, and rust-colored symbols indicate SD groups; open circles denote Sham and filled squares denote tUS. In the body-weight panel, solid and dashed lines indicate Ctrl and SD groups, respectively. Baseline values of 0% are defined by normalization rather than independent observations of zero change.

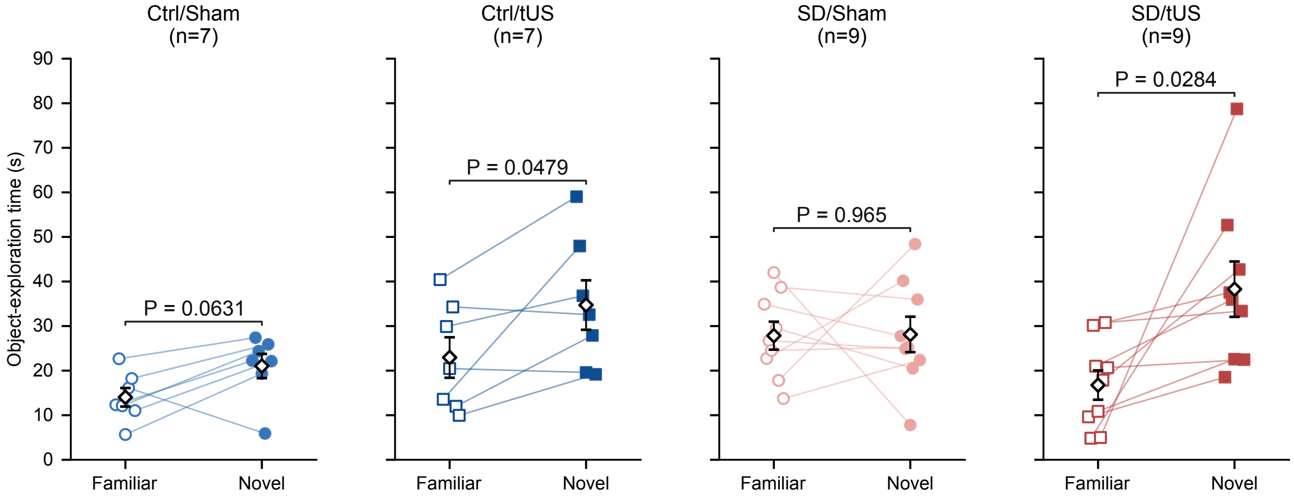


**Figure S15. Individual familiar- and novel-object exploration times during the novel object recognition test.**

Object exploration times during the Day 17 retention test are shown for Ctrl/Sham (n = 7), Ctrl/tUS (n = 7), SD/Sham (n = 9), and SD/tUS (n = 9) mice. Familiar-object and novel-object exploration times were quantified using automated object-directed approach classifications derived from DeepLabCut-based tracking. The analysis used the first 300 s of cumulative observation after excluding on-object intervals. This window could extend beyond the first 300 s of elapsed recording.

Open symbols represent familiar-object exploration times, filled symbols represent novel-object exploration times, and lines connect measurements from the same mouse. Diamonds with error bars indicate mean ± SEM. Brackets show P values from separate within-group comparisons of familiar- and novel-object exploration using two-sided paired t-tests.

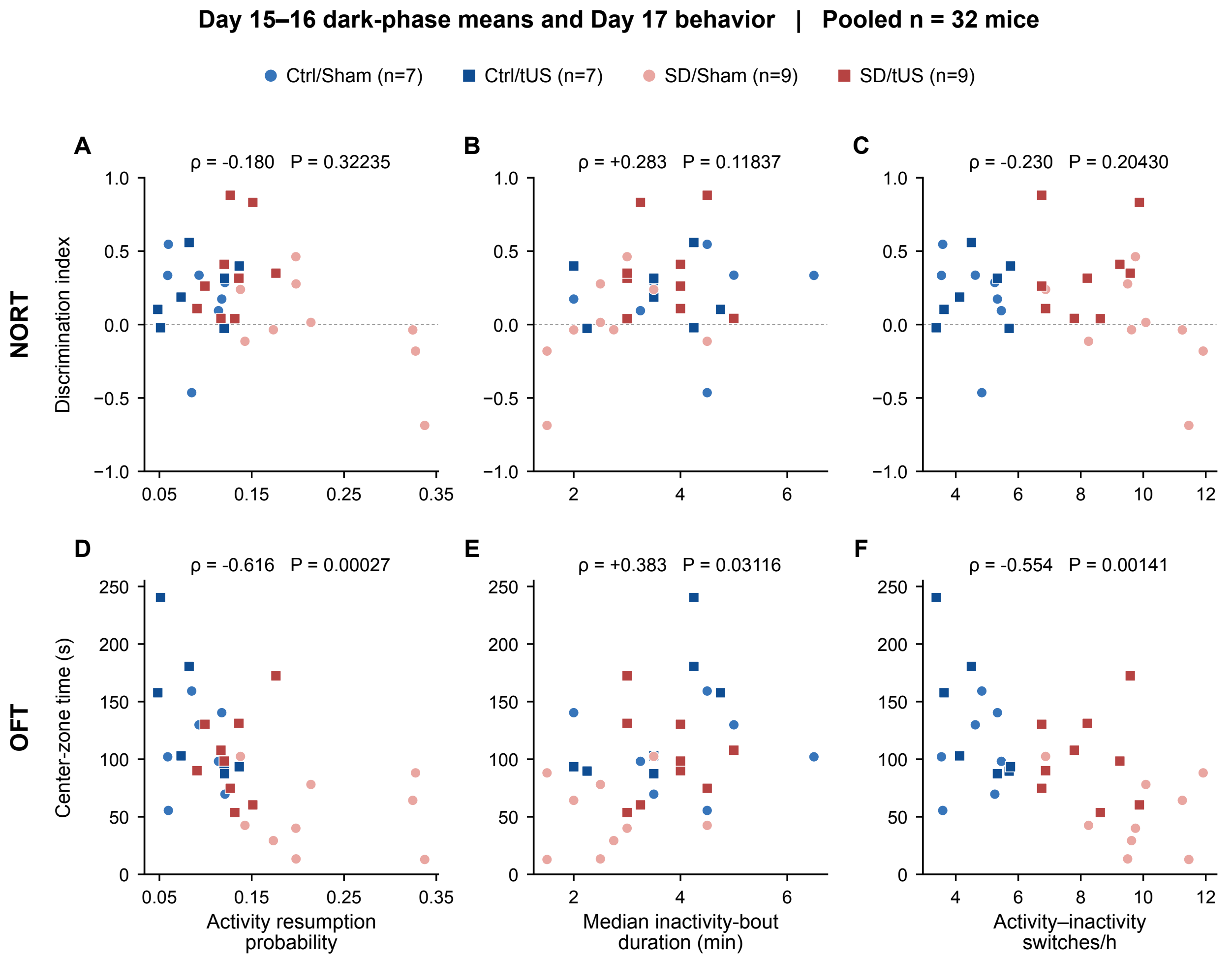


Figure S16. Wheel-activity characteristics on Days 15–16 and behavioral outcomes on Day 17 (1/2).

Two-night dark-phase means are plotted against NORT discrimination index (A–C) and OFT center time (D–F). Each point represents one mouse. Colors and symbols identify the four groups. Spearman's ρ and two-sided permutation P values were calculated across all 32 mice pooled across groups. The dotted line at DI = 0 marks equal exploration time for the novel and familiar objects (A–C). Group medians and quartiles for the wheel measures are given in Supplementary Table S4, Part C.

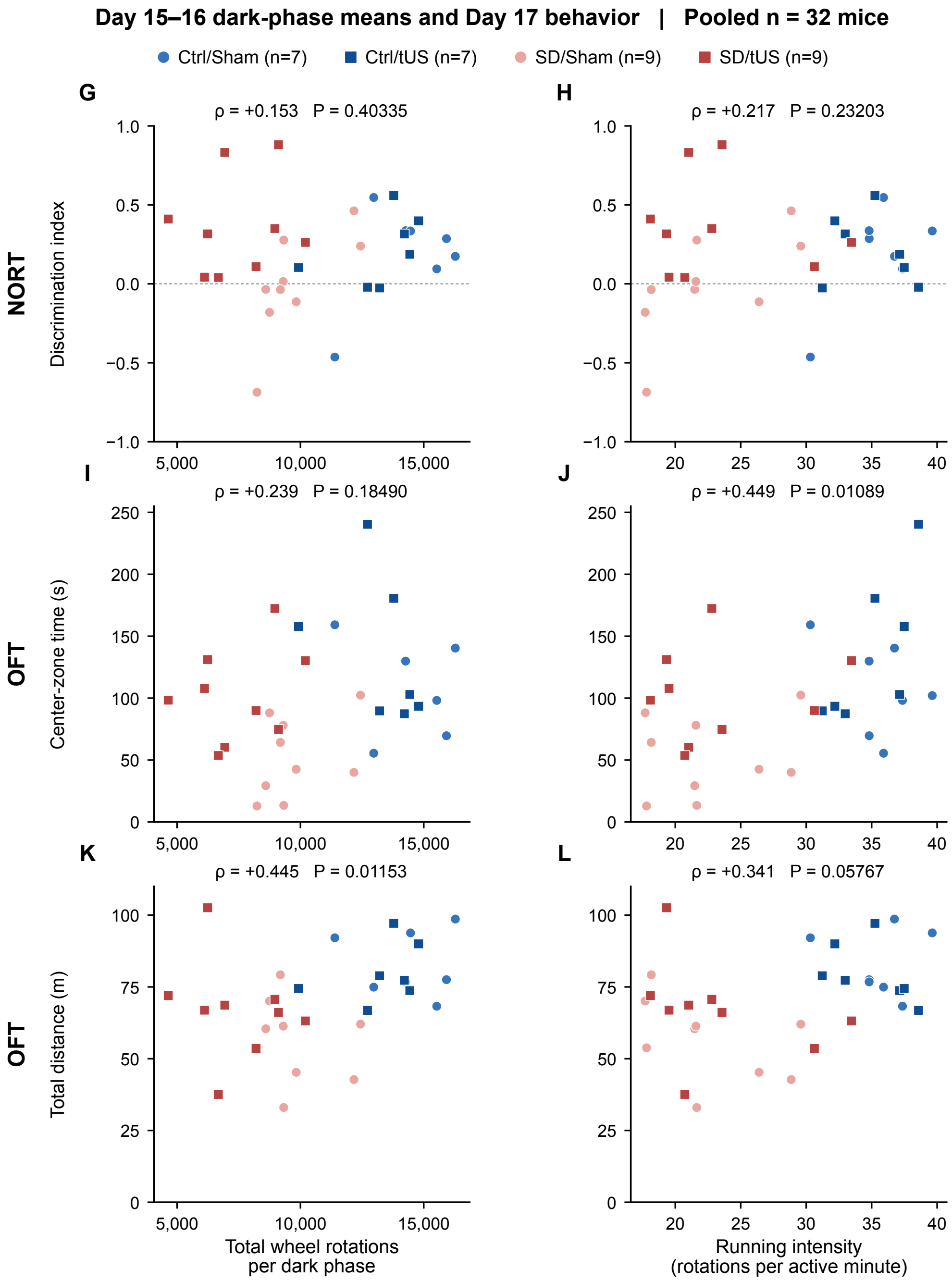


Figure S16. Wheel-activity characteristics on Days 15–16 and behavioral outcomes on Day 17 (2/2).

Total wheel rotations per dark phase (G, I, K) and running intensity in rotations per active minute (H, J, L) are shown against NORT DI (G, H), OFT center time (I, J), and total OFT distance (K, L). Each point pairs one mouse's two-night mean with its Day 17 outcome. Colors and symbols identify the four groups. Spearman's ρ and two-sided permutation P values were calculated across all 32 mice pooled across groups. The dotted line at DI = 0 marks equal exploration time for the novel and familiar objects (G, H). Group medians and quartiles are given in Supplementary Table S4, Part C.

#### Supplementary references

[1] Minakawa EN, Miyazaki K, Maruo K, Yagihara H, Fujita H, Wada K, et al. Chronic sleep fragmentation exacerbates amyloid β deposition in Alzheimer's disease model mice. Neurosci Lett. 2017;653:362-369. https://doi.org/10.1016/j.neulet.2017.05.054

[2] Sakamoto K, Higo-Yamamoto S, Egi Y, Miyazaki K, Oishi K. Memory dysfunction and anxiety-like behavior in a mouse model of chronic sleep disorders. Biochem Biophys Res Commun. 2020;529(2):175–179. https://doi.org/10.1016/j.bbrc.2020.05.218

[3] Martin E, Aubry JF, Schafer M, Verhagen L, Treeby B, Pauly KB. ITRUSST consensus on standardised reporting for transcranial ultrasound stimulation. Brain Stimul. 2024;17(3):607-615. https://doi.org/10.1016/j.brs.2024.04.013

[4] Franklin KBJ, Paxinos G. Paxinos and Franklin's The Mouse Brain in Stereotaxic Coordinates, Compact: The Coronal Plates and Diagrams. 5th ed. Academic Press; 2019. ISBN 978-0-12-816159-3.

[5] Clinard S, Wettstone E, Moore D, Snell J, Padilla F, Eames M. Low-cost 3-D hydrophone scanning tank with MATLAB GUI control. Ultrasound Med Biol. 2022;48(1):157–163. https://doi.org/10.1016/j.ultrasmedbio.2021.09.022

[6] International Electrotechnical Commission. IEC 62127-1:2022. Ultrasonics – Hydrophones – Part 1: Measurement and characterization of medical ultrasonic fields. 2nd ed. Geneva: IEC; 2022. https://webstore.iec.ch/en/publication/60227

[7] Devine J, Vidal-García M, Liu W, Neves A, Lo Vercio LD, Green RM, et al. MusMorph, a database of standardized mouse morphology data for morphometric meta-analyses. Sci Data. 2022;9(1):230. https://doi.org/10.1038/s41597-022-01338-x

[8] Hallgrimsson B, Devine J. Strain Comparison [dataset]. Version 1. FaceBase; 2021. https://doi.org/10.25550/3-JZ9J

[9] Hikishima K, Komaki Y, Seki F, Ohnishi Y, Okano HJ, Okano H. In vivo microscopic voxel-based morphometry with a brain template to characterize strain-specific structures in the mouse brain. Sci Rep. 2017;7(1):85. https://doi.org/10.1038/s41598-017-00148-1

[10] Wang Q, Ding SL, Li Y, Royall J, Feng D, Lesnar P, et al. The Allen Mouse Brain Common Coordinate Framework: A 3D Reference Atlas. Cell. 2020;181(4):936-953.e20. https://doi.org/10.1016/j.cell.2020.04.007

[11] Treeby BE, Cox BT. k-Wave: MATLAB toolbox for the simulation and reconstruction of photoacoustic wave fields. J Biomed Opt. 2010;15(2):021314. https://doi.org/10.1117/1.3360308

[12] Treeby BE, Jaros J, Rendell AP, Cox BT. Modeling nonlinear ultrasound propagation in heterogeneous media with power law absorption using a k-space pseudospectral method. J Acoust Soc Am. 2012;131(6):4324-36. https://doi.org/10.1121/1.4712021

[13] Sapozhnikov OA, Tsysar SA, Khokhlova VA, Kreider W. Acoustic holography as a metrological tool for characterizing medical ultrasound sources and fields. J Acoust Soc Am. 2015;138(3):1515-32. https://doi.org/10.1121/1.4928396

[14] Krokhmal A, Simcock IC, Treeby BE, Martin E. A comparative study of experimental and simulated ultrasound beam propagation through cranial bones. Phys Med Biol. 2025;70(2):025007. https://doi.org/10.1088/1361-6560/ada19d

[15] Efron B, Tibshirani R. Bootstrap methods for standard errors, confidence intervals, and other measures of statistical accuracy. Stat Sci. 1986;1(1):54–75. https://doi.org/10.1214/ss/1177013815

[16] Yaakub SN, White TA, Roberts J, Martin E, Verhagen L, Stagg CJ, et al. Transcranial focused ultrasound-mediated neurochemical and functional connectivity changes in deep cortical regions in humans. Nat Commun. 2023;14:5318. https://doi.org/10.1038/s41467-023-40998-0

[17] Nyborg WL. Sonically produced heat in a fluid with bulk viscosity and shear viscosity. J Acoust Soc Am. 1986;80(4):1133-9. https://doi.org/10.1121/1.393803

[18] Pennes HH. Analysis of tissue and arterial blood temperatures in the resting human forearm. J Appl Physiol. 1948;1(2):93-122. https://doi.org/10.1152/jappl.1948.1.2.93

[19] Crank J, Nicolson P. A practical method for numerical evaluation of solutions of partial differential equations of the heat-conduction type. Proc Cambridge Philos Soc. 1947;43(1):50–67. https://doi.org/10.1017/S0305004100023197

[20] Rannacher R. Finite element solution of diffusion problems with irregular data. Numer Math. 1984;43:309–327. https://doi.org/10.1007/BF01390130

[21] Hoffman DJ, Newland MC. A microstructural analysis distinguishes motor and motivational influences over voluntary running in animals chronically exposed to methylmercury and nimodipine. Neurotoxicology. 2016;54:127-139. https://doi.org/10.1016/j.neuro.2016.04.009

[22] Wiggin TD, Goodwin PR, Donelson NC, Liu C, Trinh K, Sanyal S, et al. Covert sleep-related biological processes are revealed by probabilistic analysis in Drosophila. Proc Natl Acad Sci U S A. 2020;117(18):10024–10034. https://doi.org/10.1073/pnas.1917573117

[23] Kaplan EL, Meier P. Nonparametric Estimation from Incomplete Observations. J Am Stat Assoc. 1958;53(282):457–481. https://doi.org/10.1080/01621459.1958.10501452

[24] Lueptow LM. Novel Object Recognition Test for the Investigation of Learning and Memory in Mice. J Vis Exp. 2017;(126):55718. https://doi.org/10.3791/55718

[25] Mathis A, Mamidanna P, Cury KM, Abe T, Murthy VN, Mathis MW, et al. DeepLabCut: markerless pose estimation of user-defined body parts with deep learning. Nat Neurosci. 2018;21(9):1281-1289. https://doi.org/10.1038/s41593-018-0209-y

[26] Nath T, Mathis A, Chen AC, Patel A, Bethge M, Mathis MW. Using DeepLabCut for 3D markerless pose estimation across species and behaviors. Nat Protoc. 2019;14(7):2152-2176. https://doi.org/10.1038/s41596-019-0176-0

[27] Ye S, Filippova A, Lauer J, Schneider S, Vidal M, Qiu T, et al. SuperAnimal pretrained pose estimation models for behavioral analysis. Nat Commun. 2024;15(1):5165. https://doi.org/10.1038/s41467-024-48792-2

[28] Kalman RE. A New Approach to Linear Filtering and Prediction Problems. J Basic Eng. 1960;82(1):35–45. https://doi.org/10.1115/1.3662552

[29] Seibenhener ML, Wooten MC. Use of the Open Field Maze to measure locomotor and anxiety-like behavior in mice. J Vis Exp. 2015;(96):e52434. https://doi.org/10.3791/52434

[30] MacKinnon JG, White H. Some heteroskedasticity-consistent covariance matrix estimators with improved finite sample properties. J Econom. 1985;29(3):305–325. https://doi.org/10.1016/0304-4076(85)90158-7

[31] Shapiro SS, Wilk MB. An analysis of variance test for normality (complete samples). Biometrika. 1965;52(3–4):591–611. https://doi.org/10.2307/2333709

[32] Jarque CM, Bera AK. Efficient tests for normality, homoscedasticity and serial independence of regression residuals. Econ Lett. 1980;6(3):255–259. https://doi.org/10.1016/0165-1765(80)90024-5

[33] Koenker R. A note on studentizing a test for heteroscedasticity. J Econom. 1981;17(1):107–112. https://doi.org/10.1016/0304-4076(81)90062-2

[34] Cook RD. Detection of influential observation in linear regression. Technometrics. 1977;19(1):15–18. https://doi.org/10.1080/00401706.1977.10489493

[35] Holm S. A simple sequentially rejective multiple test procedure. Scand J Stat. 1979;6(2):65–70. https://www.jstor.org/stable/4615733

[36] Spearman C. The Proof and Measurement of Association between Two Things. Am J Psychol. 1904;15(1):72–101. https://doi.org/10.2307/1412159

[37] Phipson B, Smyth GK. Permutation P-values should never be zero: calculating exact P-values when permutations are randomly drawn. Stat Appl Genet Mol Biol. 2010;9:Article39. https://doi.org/10.2202/1544-6115.1585

[38] Mang GM, Franken P. Sleep and EEG Phenotyping in Mice. Curr Protoc Mouse Biol. 2012;2(1):55–74. https://doi.org/10.1002/9780470942390.mo110126

[39] Yamada RG, Matsuzawa K, Ode KL, Ueda HR. An automated sleep staging tool based on simple statistical features of mice electroencephalography (EEG) and electromyography (EMG) data. Eur J Neurosci. 2024;60(7):5467-5486. https://doi.org/10.1111/ejn.16465

[40] Mochizuki T, Crocker A, McCormack S, Yanagisawa M, Sakurai T, Scammell TE. Behavioral state instability in orexin knock-out mice. J Neurosci. 2004;24(28):6291-300. https://doi.org/10.1523/JNEUROSCI.0586-04.2004

[41] Brodersen KH, Ong CS, Stephan KE, Buhmann JM. The balanced accuracy and its posterior distribution. 2010 20th International Conference on Pattern Recognition. IEEE; 2010. p. 3121–3124. https://doi.org/10.1109/ICPR.2010.764

[42] Harris FJ. On the use of windows for harmonic analysis with the discrete Fourier transform. Proc IEEE. 1978;66(1):51–83. https://doi.org/10.1109/proc.1978.10837

[43] Fisher SP, Cui N, McKillop LE, Gemignani J, Bannerman DM, Oliver PL, et al. Stereotypic wheel running decreases cortical activity in mice. Nat Commun. 2016;7:13138. https://doi.org/10.1038/ncomms13138

[44] Milinski L, Fisher SP, Cui N, McKillop LE, Blanco-Duque C, Ang G, et al. Waking experience modulates sleep need in mice. BMC Biol. 2021;19:65. https://doi.org/10.1186/s12915-021-00982-w

[45] Hemerik J, Goeman J. Exact testing with random permutations. TEST. 2018;27(4):811–825. https://doi.org/10.1007/s11749-017-0571-1

[46] Jud C, Schmutz I, Hampp G, Oster H, Albrecht U. A guideline for analyzing circadian wheel-running behavior in rodents under different lighting conditions. Biol Proced Online. 2005;7:101–116. https://doi.org/10.1251/bpo109

[47] Blume C, Santhi N, Schabus M. nparACT package for R: A free software tool for the non-parametric analysis of actigraphy data. MethodsX. 2016;3:430–435. https://doi.org/10.1016/j.mex.2016.05.006

[48] Van Someren EJ, Swaab DF, Colenda CC, Cohen W, McCall WV, Rosenquist PB. Bright light therapy: improved sensitivity to its effects on rest-activity rhythms in Alzheimer patients by application of nonparametric methods. Chronobiol Int. 1999;16(4):505-18. https://doi.org/10.3109/07420529908998724

[49] Nelson W, Tong YL, Lee JK, Halberg F. Methods for cosinor-rhythmometry. Chronobiologia. 1979;6(4):305-23. https://pubmed.ncbi.nlm.nih.gov/548245/

[50] Berens P. CircStat: a MATLAB toolbox for circular statistics. J Stat Softw. 2009;31(10):1–21. https://doi.org/10.18637/jss.v031.i10

[51] Huang L, Chen X, Tao Q, Wang X, Huang X, Fu Y, et al. Bright light treatment counteracts stress-induced sleep alterations in mice, via a visual circuit related to the rostromedial tegmental nucleus. PLoS Biol. 2023;21(9):e3002282. https://doi.org/10.1371/journal.pbio.3002282

[52] Yu X, Zecharia A, Zhang Z, Yang Q, Yustos R, Jager P, et al. Circadian factor BMAL1 in histaminergic neurons regulates sleep architecture. Curr Biol. 2014;24(23):2838-44. https://doi.org/10.1016/j.cub.2014.10.019

[53] Phipps MA, Mishra A, Donovan CL, Newton AT, McKnight CD, Dockum AQ, et al. Magnetic Resonance Imaging Monitoring of the Safety of Repeated Low-Intensity Focused Ultrasound Exposure at Three Brain Locations. Neuromodulation. 2026;29(2):243-254. https://doi.org/10.1016/j.neurom.2025.09.313

[54] Wang H, Sun Y, Wang Y, Chen Y, Ge Y, Yuan J, et al. Temperature-Controlled Hyperthermia with Non-Invasive Temperature Monitoring through Speed of Sound Imaging. Applied Sciences. 2023;13(12):7317. https://doi.org/10.3390/app13127317

[55] Liang B, Liu W, Zhan Q, Li M, Zhuang M, Liu QH, et al. Impacts of the murine skull on high-frequency transcranial photoacoustic brain imaging. J Biophotonics. 2019;12(7):e201800466. https://doi.org/10.1002/jbio.201800466

[56] Pinton G, Aubry JF, Bossy E, Muller M, Pernot M, Tanter M. Attenuation, scattering, and absorption of ultrasound in the skull bone. Med Phys. 2012;39(1):299-307. https://doi.org/10.1118/1.3668316
